# Lateral entorhinal cortex inputs modulate hippocampal dendritic excitability by recruiting a local disinhibitory microcircuit

**DOI:** 10.1101/2022.01.13.476247

**Authors:** Olesia M. Bilash, Spyridon Chavlis, Panayiota Poirazi, Jayeeta Basu

## Abstract

The lateral entorhinal cortex (LEC) provides information about multi-sensory environmental cues to the hippocampus through direct inputs to the distal dendrites of CA1 pyramidal neurons. A growing body of work suggests that LEC neurons perform important functions for episodic memory processing, coding for contextually-salient elements of an environment or the experience within it. However, we know little about the functional circuit interactions between LEC and the hippocampus. In this study, we combine functional circuit mapping and computational modeling to examine how long-range glutamatergic LEC projections modulate compartment-specific excitation-inhibition dynamics in hippocampal area CA1. We demonstrate that glutamatergic LEC inputs can drive local dendritic spikes in CA1 pyramidal neurons, aided by the recruitment of a disinhibitory vasoactive intestinal peptide (VIP)-expressing inhibitory neuron microcircuit. Our circuit mapping further reveals that, in parallel, LEC also recruits cholecystokinin (CCK)-expressing inhibitory neurons, which our model predicts act as a strong suppressor of dendritic spikes. These results provide new insight into a cortically-driven GABAergic microcircuit mechanism that gates non-linear dendritic computations, which may support compartment-specific coding of multi-sensory contextual features within the hippocampus.

**HIGHLIGHTS:**

1. Slice electrophysiology experiments investigate how lateral entorhinal cortex influences hippocampal area CA1
2. LEC drives local spikes in distal dendrites but not in somata of CA1 pyramidal neurons
3. LEC inputs recruit VIP IN and CCK IN populations in CA1, but not SST INs
4. Computational modeling and circuit manipulation experiments identify a VIP IN-mediated disinhibitory microcircuit for gating local dendritic spike generation

**IN BRIEF:** Bilash et al. found that a distal cortical input is capable of driving local dendritic spikes in hippocampal pyramidal neurons. This dendritic spike generation is promoted by cortical recruitment of a local VIP interneuron-mediated disinhibitory microcircuit. Their results highlight new circuit mechanisms by which dynamic interaction of excitation, inhibition, and disinhibition support supralinear single-cell computations.

## INTRODUCTION

The formation of multi-sensory episodic memories relies on the functional interactions between the entorhinal cortex and dorsal hippocampus (Basu and Siegelbaum, 2015; Brun et al., 2008; Brun et al., 2002; Igarashi et al., 2014; Kitamura et al., 2014; Remondes and Schuman, 2004). The entorhinal cortex is the main source of integrated multimodal sensory input to the hippocampus, and is anatomically and functionally subdivided into the medial entorhinal cortex (MEC) and lateral entorhinal cortex (LEC) (Hargreaves et al., 2005; Kerr et al., 2007; Witter et al., 2017). Recent work has begun exploring the functional interactions between MEC, the major source of spatial information, and hippocampal area CA1, the major output area of the hippocampus (Bittner et al., 2015; Hales et al., 2014; Jun et al., 2020; Kitamura et al., 2014; Kitamura et al., 2015; Masurkar et al., 2017; Schlesiger et al., 2018; Suh et al., 2011; Zhang et al., 2013). On the other hand, we know little about the circuit organization and function of LEC inputs in shaping CA1 activity, despite LEC being the major source of contextual sensory information. Here, we define how direct glutamatergic inputs from LEC to distal dendrites of CA1 pyramidal neurons (Desmond et al., 1994; Kajiwara et al., 2008; Yeckel and Berger, 1995) sculpt compartment-specific information processing in the hippocampus by recruiting a distinct GABAergic microcircuit.

The lateral entorhinal cortex is a key interaction partner of the hippocampus (Basu et al., 2016; Cui et al., 2013; Li et al., 2017; Masurkar et al., 2017; Nilssen et al., 2018; Ruth et al., 1988; van Groen et al., 2003; Witter et al., 2017; Witter et al., 1989) and is therefore poised to have significant, widespread impact on single-cell computations, circuit dynamics, and behavior. *In vivo* experiments in rodents have demonstrated that LEC codes for important environmental features and sensory experiences (Hargreaves et al., 2005). These include odors (Igarashi et al., 2014; Leitner et al., 2016; Li et al., 2017; Xu and Wilson, 2012), object novelty (Basu et al., 2016; Deshmukh and Knierim, 2011; Tsao et al., 2013; Wilson et al., 2013), object-place associations (Kuruvilla et al., 2020; Tsao et al., 2013; Wang et al., 2018), contextual salience (Basu et al., 2016), temporal structure (Tsao et al., 2018) and cue-reward associations (Lee et al., 2021) during sensory learning experiences. These behavior studies demonstrate that LEC input may carry multi-sensory information to the hippocampus that is critical in supporting episodic memory representations. At the circuit level, studies have uncovered that LEC sends glutamatergic (Li et al., 2017; Masurkar et al., 2017) and GABAergic (Basu et al., 2016) projections directly to hippocampal area CA1, which play crucial roles in associative, context-dependent learning. However, we still lack a basic understanding of how excitatory drive from LEC supports dendritic and somatic computations and circuit dynamics in service of memory processing in the hippocampus.

How might distal dendritic inputs from LEC shape hippocampal activity? We hypothesize that glutamatergic LEC-CA1 projections sculpt nonlinear neuronal computations by tuning compartment-specific excitation-inhibition balance. Dendritic spikes are a prime example of such nonlinear computations and can be effectively modulated by long-range and local GABAergic circuits (Basu et al., 2016; Grienberger et al., 2017; Lovett-Barron et al., 2012; Milstein et al., 2015; Moore et al., 2021; Palmer et al., 2013; Stuart and Spruston, 2015). They are known to underlie vital neural processes such as long-term plasticity (Gambino et al., 2014; Golding et al., 2002; Kim et al., 2015) and feature-selective tuning (Bittner et al., 2017; Cichon and Gan, 2015; Jia et al., 2010; Smith et al., 2013). However, there is a major gap in our mechanistic understanding of the neural circuits that gate input-driven dendritic spikes in principal neurons. This is, in part, due to the experimental difficulty in combining distal dendritic recordings and circuit manipulations. Past studies used compartmental modeling to infer the ionic, synaptic, and anatomical determinants of dendritic spikes, as well as their contribution to single-neuron computations (Gasparini et al., 2004; Park et al., 2019; Pissadaki et al., 2010; Poirazi et al., 2003a, b; Smith et al., 2013). However, such models often fail to capture important dendritic, microcircuit, and input pathway characteristics due to a lack of experimental data, which limits the process of constraining model parameters. As a result, the excitatory-inhibitory circuit interactions that shape dendritic nonlinearities have remained especially elusive. GABAergic inhibitory neurons (INs) serve as rapid and precise gain modulation switches to shape single-cell computations within a neural circuit (Basu et al., 2016; Grienberger et al., 2017; Lovett-Barron et al., 2012; Milstein et al., 2015; Moore et al., 2021). Given their functional diversity and wiring specificity (Klausberger and Somogyi, 2008; Pelkey et al., 2017), INs can boost or suppress compartment-specific activity, and can therefore expand the computational coding capacity of single neurons and neural circuits as a whole. Various single-neuron models have examined the effects of inhibition on local spiking (Bloss et al., 2016; Gidon and Segev, 2012; Jadi et al., 2012; Wilmes et al., 2016), while network-level models have predicted the role of specific interneuron populations in hippocampal place cell formation (Pedrosa and Clopath, 2020; Shuman et al., 2020; Turi et al., 2019). However, these models were not constrained in such a way that they could investigate how a specific input shapes compartment-specific excitation-inhibition balance within a single neuron. Combining experimental and modeling approaches to study LEC-CA1 circuit interactions allows for a powerful investigation of how input-driven GABAergic microcircuit motifs can shape the local input-output transformation within dendritic branches and across the neuron. Dendritic integration of excitatory and inhibitory inputs is a fundamental feature of principal neurons across the brain. Thus, an approach combining experiments and computational modeling can be widely applied to investigate input-driven excitation-inhibition balance and nonlinear computations in other brain regions.

In the present study, we used *in vitro* electrophysiology, optogenetics, and multi-compartmental modeling to examine the cellular and circuit mechanisms by which long-range glutamatergic LEC projections influence the local activity in hippocampal area CA1. We photostimulated glutamatergic LEC axons and recorded light-evoked responses from various neuronal populations in area CA1. Photostimulation of LEC inputs could drive local dendritic spikes in CA1 PN distal dendrites without eliciting somatic action potentials. To uncover the circuit mechanism gating these dendritic spikes, we explored the role of local GABAergic inhibitory neurons. LEC inputs were found to directly recruit vasoactive intestinal peptide (VIP)- and cholecystokinin (CCK)-expressing IN populations. We used our data to develop and constrain a circuit model of an LEC-driven multi-compartment CA1 pyramidal neuron and surrounding GABAergic microcircuitry to simulate the deletion of specific interneuron populations and infer their role in shaping compartment-specific LEC-driven activity in CA1 PNs. The model predicted that a disinhibitory, VIP IN-mediated microcircuit permits LEC-driven dendritic spikes. Optogenetic silencing of VIP INs led to a decreased incidence of LEC-driven dendritic spikes, verifying the model prediction. Taken together, our findings demonstrate for the first time that a VIP IN-mediated disinhibitory microcircuit gates cortically-driven, local dendritic spikes in area CA1. Therefore, our study establishes a circuit mechanism by which LEC promotes supralinear dendritic computations in the hippocampus.

## RESULTS

### LEC directly excites CA1 pyramidal neurons, driving depolarization of dendrites and somata

To explore the synaptic nature of LEC excitatory inputs, we performed optogenetic functional circuit mapping of the direct, glutamatergic projections from LEC to pyramidal neurons in area CA1 of the hippocampus in acute hippocampal slices. We unilaterally injected a recombinant adeno-associated virus (rAAV-CaMKII-ChR2-eYFP) into the superficial layers of LEC to express ChR2-eYFP specifically in glutamatergic LEC neurons and their axons (**Figure 1A**, **Figure S1A-B**). High-resolution confocal imaging confirmed that expression of ChR2-eYFP was restricted to LEC (**Figure 1B**, **Figure S1B**). Local photostimulation over LEC confirmed adequate expression and activation of ChR2 in LEC neurons (**Figure S1C-D**).

**Figure 1.**
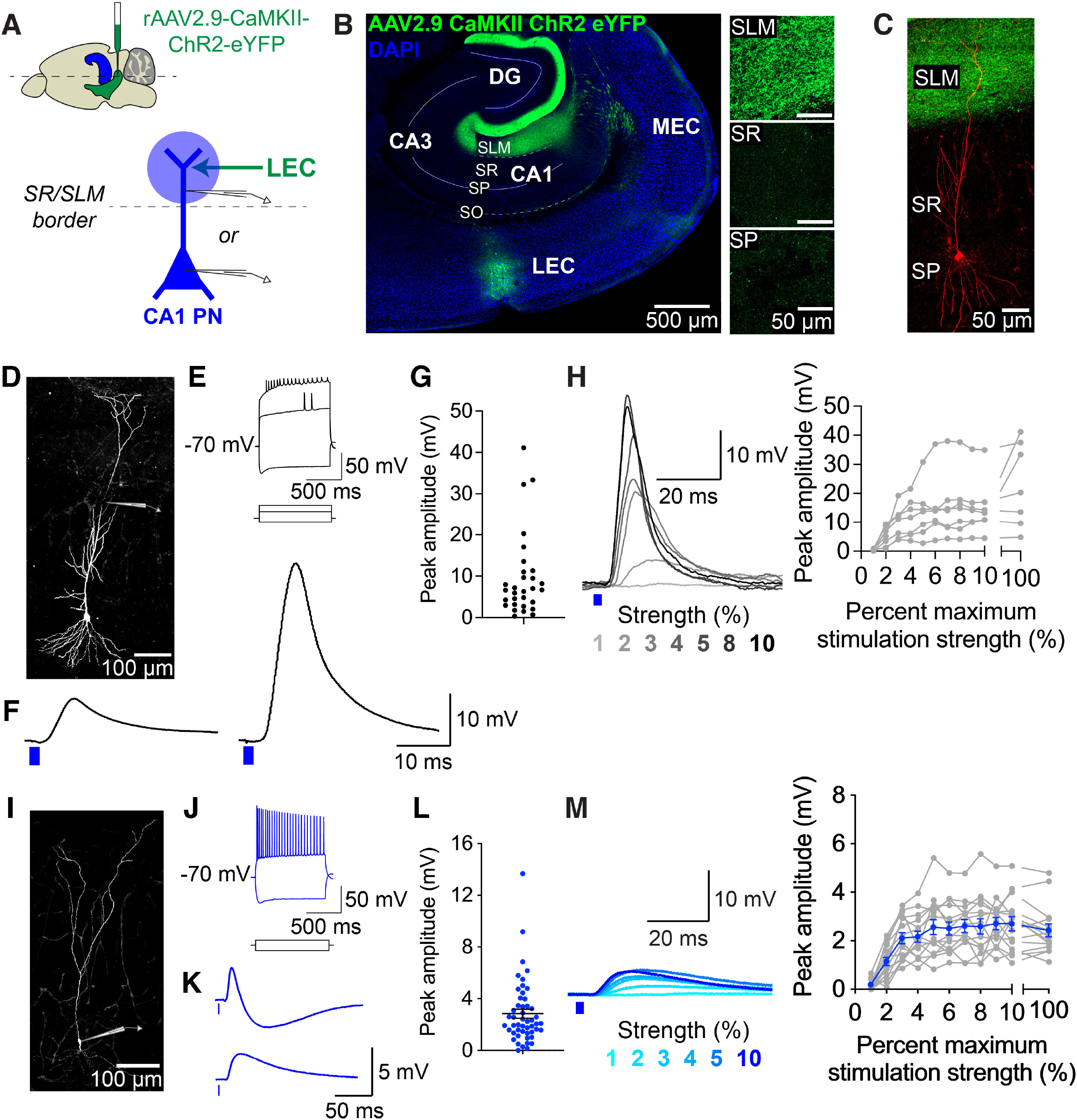
LEC inputs provide direct excitation to CA1 pyramidal neuron dendrites and somata. (A) Schematic of experimental design. Top: sagittal view of a mouse brain, illustrating the LEC injection site (green) and HC (blue). Bottom: CA1 PN (in blue) depicting the location of photostimulation (blue shaded circle) to activate ChR2^+^ LEC axons (green arrow) in SLM. Whole-cell recording sites on the CA1 PN dendrite or soma. (B) Confocal image of a horizontal brain slice demonstrating the LEC injection site and ChR2^+^ LEC axons within HC. Right insets: ChR2^+^ LEC axons in the various CA1 layers showing their high concentration in SLM. (C) Confocal image of a typical CA1 pyramidal neuron morphology (in red). Recorded CA1 PN was filled intracellularly with biocytin and counterstained with streptavidin conjugated with Alexa 594. The ChR2-eYFP^+^ glutamatergic LEC axons and axon terminals were labeled with anti-GFP (green). (D) Confocal image of a CA1 PN, recorded at the dendrite (patch pipette shows recording site). The CA1 PN was filled intracellularly with biocytin and was later counterstained with streptavidin conjugated with Alexa 647. (E) Example traces illustrating the intrinsic electrophysiological properties of a CA1 PN dendrite. The firing and sag were measured from a distal dendrite in response to 1000 ms-long depolarizing (+700 and +400 pA) and hyperpolarizing (-325 pA) dendritic current injections, respectively. The dendritic membrane potential was maintained at -70 mV. Note the filtered spikelets, which are typical of distal dendritic recordings. (F) Example traces of LEC-driven post-synaptic responses recorded in CA1 PN distal dendrites in current clamp mode in response to a 2 ms photostimulation of LEC axons with 470 nm light over SLM. Left: putative subthreshold dendritic response. Right: putative suprathreshold dendritic response. (G) Scatterplot of the peak amplitudes of LEC-driven responses in CA1 PN distal dendrites. Min to Max amplitude: 1.76 to 41.1 mV. Average ± SEM: 12.9 ± 2.7 mV. Total n = 17 dendrites, 16 slices, 13 mice. (H) Input-output transformation. Left: Example traces of the LEC-driven responses in CA1 PN dendrites as photostimulation strength increases from 1-10% of the maximum. Putative suprathreshold dendritic response seen at 8% maximum photostimulation strength. Right: Summary graph of the input-output transformation amplitudes. n = 7 dendrites, 7 slices, 7 mice. (I) Confocal image of a CA1 PN, recorded at the soma (patch pipette shows recording site). The CA1 PN was filled intracellularly with biocytin and was later counterstained with streptavidin conjugated to Alexa 647. (J) Electrophysiological properties of a CA1 PN. While the somatic membrane potential was maintained at -70 mV, the firing and sag were measured in response to depolarizing (+400) and hyperpolarizing (-375 pA) somatic current injections, respectively. (K) Example traces of LEC-driven post-synaptic potentials (PSPs) recorded in CA1 PN somata, using the same protocols as described in F. Example somata shown that respond with (top) or without (bottom) a putative IPSP. (L) Scatterplot of the peak amplitudes of LEC-driven responses in CA1 PN soma. Average ± SEM: 2.84 ± 0.35 mV. Total n = 50 pyramidal neuron somata, 48 slices, 33 mice. (M) Input-output transformation. Left: Example traces of the LEC-driven responses in CA1 PN somata, generated using the same protocols as described in H. Right: Summary graph of the input-output transformation amplitudes. Data from individual somata shown in gray. Average ± SEM data shown in blue. n = 15 pyramidal neuron somata, 14 slices, 11 mice.

Next, we examined LEC projections to hippocampal area CA1, a major target region for the direct excitatory input from LEC and the output region of the hippocampus. As expected (Masurkar et al., 2017; van Groen et al., 2003; Witter et al., 1989), glutamatergic LEC axons project directly into the distal dendrite layer of hippocampal area CA1 (**Figure 1B-C**). The projections follow a density gradient along the proximal-distal axis of CA1 (Masurkar et al., 2017; Witter et al., 1989), with higher levels of fluorescent ChR2-eYFP^+^ axonal fibers in the distal part of area CA1, close to the subiculum. To determine the synaptic connectivity and strength of the LEC glutamatergic inputs, we locally photostimulated the ChR2^+^ LEC axon terminals in *stratum lacunosum moleculare* (SLM) of area CA1 with 470 nm light (single, 2 ms pulse, 100% LED strength, 56 mW, 50 µm diameter beam spot) and recorded light-evoked responses from CA1 pyramidal neuron (PNs) (**Figures 1**-**2**). Using intracellular, whole-cell recordings in current clamp mode, we recorded robust, light-evoked post-synaptic responses in CA1 PN somata, which persisted in the presence of tetrodotoxin (TTX, 1 μM, to prevent sodium spikes) and 4-aminopyridine (4-AP, 100 μM, to block repolarizing potassium channels), a pharmacological condition that eliminates polysynaptic inputs and feed-forward inhibition (Petreanu et al., 2009) (**Figure S2I**), indicating mono-synaptic connectivity between LEC and CA1. Under these conditions, we sometimes observed an increase in PSP amplitude, which may have resulted from the elimination of feed-forward inhibition and/or other hyperpolarizing ionic conductances.

**Figure 2.**
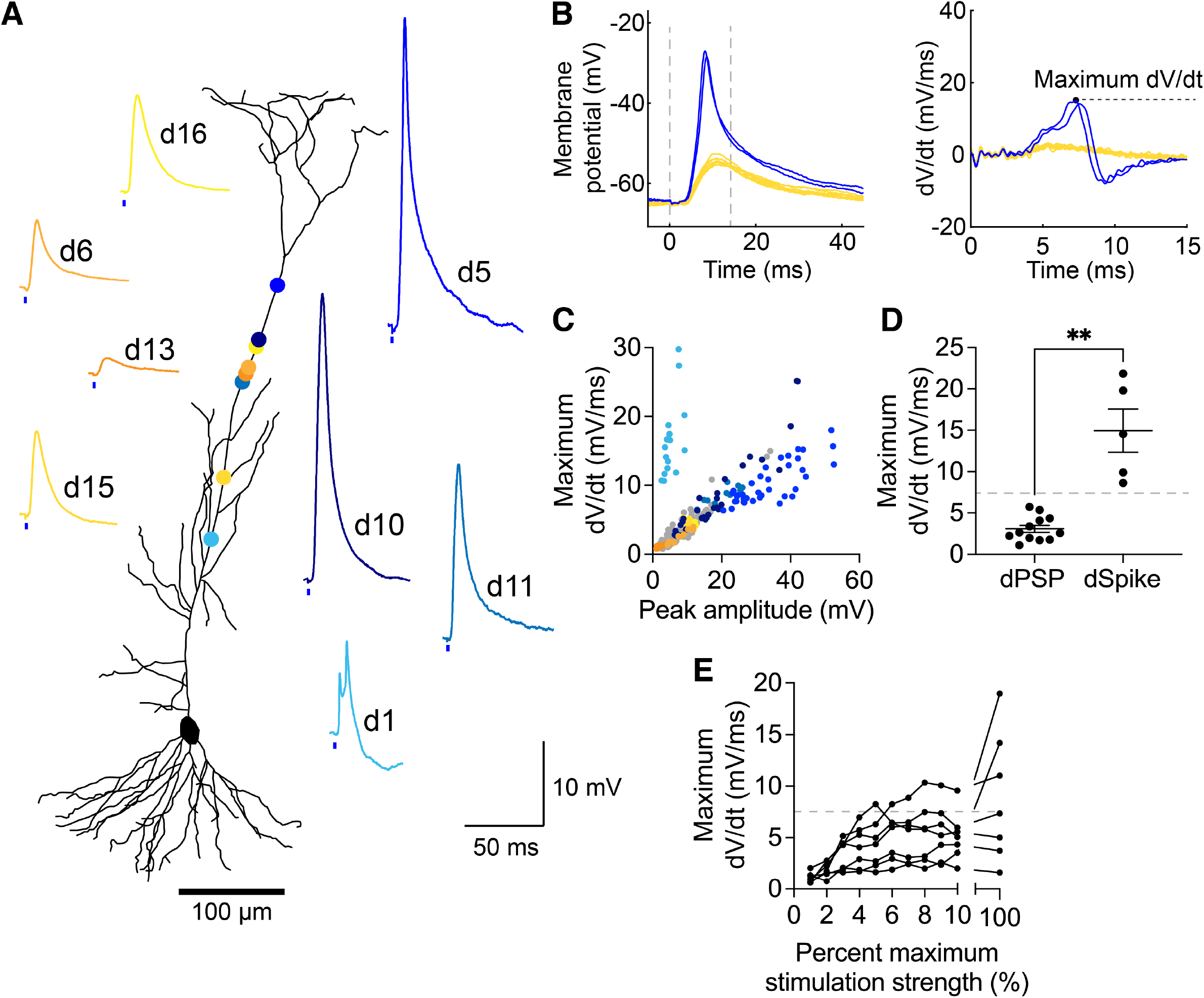
Optogenetic stimulation of LEC inputs can drive dendritic spikes in CA1 pyramidal neurons. (A) Example traces of LEC-driven dendritic responses recorded in current clamp mode in CA1 PN dendrites and the corresponding recording location on the dendritic tree. Dendrites are numbered in the order they were recorded (‘d1’ and so forth). Putative subthreshold dendritic post-synaptic potentials (dPSPs) are shown in shades of yellow, whereas putative dendritic spikes (dSpikes) are shown in shades of blue. Neuronal morphology was reconstructed from the neuron shown in Figure 1D. Total n = 17 dendrites. (B) Left: Example traces of LEC-driven dSpikes (blue) and dPSPs (yellow). Right: Derivative of the LEC-driven dendritic responses, shown from t = 0 to t = 15 ms (gray dashed line in left graph). Maximum dV/dt is calculated as the peak amplitude of the derivative traces. (C) Scatterplot of the peak amplitude versus maximum dV/dt values of every LEC-driven dendritic response recorded. Total n = 17 dendrites (∼15 sweeps recorded/dendrite, on average). Data from the 8 example dendrites are color coded, as in A; data from the remaining 9 dendrites are in gray. (D) Average maximum dV/dt values of LEC-driven dPSPs versus dSpikes. p = 0.0097, Unpaired t-test with Welch’s correction, n = 14 dPSPs and 5 dSpikes. Dashed line indicates the threshold value from our dendritic data set that most accurately categorizes dPSPs versus dSpikes. (E) Input-output transformation, demonstrating the maximum dV/dt values of the LEC-driven dendritic responses as photostimulation strength increases. Notably, some LEC-driven responses transition from subthreshold to suprathreshold as the input-output curve crosses the threshold (dV/dt = 7.5 mV/ms, dashed line). n = 7 dendrites, 7 slices, 7 mice.

After confirming the monosynaptic connectivity between glutamatergic LEC projections and CA1 pyramidal neurons, we characterized the light-evoked post-synaptic responses under more physiological, drug-free conditions in CA1 PN dendrites and soma, while local inhibition and polysynaptic connections remained intact (**Figure 1**). As long-range extrahippocampal inputs, glutamatergic LEC axons synapse onto the distal dendrites of CA1 PNs (**Figure 1B-C**). To directly access the LEC-driven activity in the dendritic compartment, we performed dendritic patch-clamp recordings from CA1 PN distal dendrites, 225-400 µm from the soma, around the *stratum radiatum*/*stratum lacunosum moleculare* (SR/SLM) border (**Figure 1A, D-E**, **Figure S2A-D**). Photostimulation with 100% LED power resulted in dendritic post-synaptic responses that ranged in amplitude and kinetics (Peak amplitude: 1.8 to 41.1 mV, Time to peak: 4.2 to 12.6 ms, Half-width: 2.3 to 32.5 ms, n = 17 dendrites) (**Figure 1F-G**). To measure the input-output (I/O) transformation of the responses, we photostimulated the LEC axons across a range of lower strength light intensities to recruit an increasing number of ChR2^+^ LEC axons (**Figure 1H**). The amplitudes of LEC-driven dendritic responses saturated after 6% maximum stimulation strength, with some amplitudes ultimately exceeding 30 mV at higher stimulation strengths. Some LEC-driven dendritic responses exhibited a shape reminiscent of suprathreshold dendritic spikes previously described in the literature (Gasparini et al., 2004; Hausser et al., 2000; Remy et al., 2009; Remy and Spruston, 2007).

To assess how the LEC-driven responses propagate to the soma, we also performed whole-cell recordings from CA1 PN somata (**Figure 1I-J**, **Figure S2E-H**), using the same photostimulation paradigms as for dendrites. The somatic post-synaptic responses also exhibited a range of amplitudes and kinetics (Peak amplitude: 1.5 to 13.7 mV (2.9 ± 0.3 mV), Time to peak: 9.8 to 40.5 ms, Half-width: 7.7 to 115 ms, n = 51 somata) (**Figure 1K-L**). The amplitudes of LEC-driven somatic responses also saturated after 6% maximum stimulation strength (**Figure 1M**). Although the net depolarization driven by glutamatergic LEC inputs in the CA1 PN soma was larger than the depolarization resulting from electrical stimulation of the perforant path (includes MEC and LEC axonal bundles; 0.5 to 2 mV (Basu et al., 2013; Dudman et al., 2007; Dvorak-Carbone and Schuman, 1999), compare to **Figure 1L**), it was never enough to elicit LEC-driven action potentials in the CA1 PN soma. In summary, we demonstrated that glutamatergic LEC inputs drive depolarization in the dendritic and somatic compartments of CA1 PNs, albeit without leading to action potential generation.

### LEC drives supralinear output in dendrites but not in the soma

As stated, the shape of some LEC-driven dendritic responses resembled suprathreshold dendritic spikes previously described in the literature (Basu et al., 2016; Hausser et al., 2000; Remy et al., 2009), such as putative sodium spikes (Gasparini et al., 2004; Golding and Spruston, 1998; Golding et al., 2002; Losonczy and Magee, 2006; Remy and Spruston, 2007) (**Figure 2A**, examples d5, d10, d11) and complex calcium spikes (Golding et al., 1999; Stuart et al., 1997) (**Figure 2A**, example d1). This was surprising, as single inputs rarely elicit dendritic spikes without the aid of coincident inputs (Basu et al., 2013; Pissadaki et al., 2010; Takahashi and Magee, 2009), burst activity (Golding et al., 2002; Kim et al., 2015; Takahashi and Magee, 2009), inhibition blockade (Golding et al., 2005; Golding and Spruston, 1998; Golding et al., 2002; Takahashi and Magee, 2009), and simultaneous depolarizing current injections (Bittner et al., 2015; Golding et al., 2002; Milstein et al., 2015).

To validate that these were indeed LEC-driven dendritic spikes, rather than large-amplitude dendritic PSPs, we analyzed the derivative traces (Gasparini et al., 2004; Losonczy and Magee, 2006; Muller et al., 2012; Remy et al., 2009) and phase plots of all LEC-driven responses. Both the derivative traces and phase plots revealed obvious differences between LEC-driven dendritic PSPs and dendritic spikes (**Figure 2B-C**, **Figure S3**). The derivative traces of LEC-driven dendritic spikes exhibited a recognizable, high amplitude peak that reached a maximum dV/dt value before dipping back below zero. In contrast, the derivative traces of LEC-driven dendritic PSPs exhibited a smooth, flattened curve (**Figure 2B**, **Figure S3B**). Based on these characteristics, we set a dendritic spike classification criterion for our slice electrophysiology dataset: light-evoked dendritic responses with maximum dV/dt > 7.5 mV/ms were categorized as suprathreshold dendritic spikes (dSpikes), while those with maximum dV/dt < 7.5 mV/ms were categorized as subthreshold dendritic post-synaptic potentials (dPSPs). Compared to other measurements, the maximum dV/dt value provided the clearest, quantifiable delineation of subthreshold versus suprathreshold LEC-driven dendritic responses (**Figure 2D**, **Figure S4**). Interestingly, among the recorded dendritic responses, the incidence of LEC-driven dendritic spikes did not appear to be strictly distance dependent within the distal dendritic compartment (**Figure 2A**, **Figure S4B**). Eliciting LEC-driven dendritic spikes was possible at lower photostimulation strengths (4-10% maximum stimulation strength), and was not limited to the maximum (100%) stimulation strength protocol (**Figure 2E**). LEC-driven PSPs recorded at the dendrite were significantly larger in amplitude and had faster kinetics than those recorded in the soma of CA1 PNs (Peak amplitude p < 0.0001, Time of peak p < 0.0001, Half-width p < 0.0001, Mann-Whitney test, n = 14 dPSPs and 52 somatic PSPs, **Figure S5**). This is expected, given the proximity of the dendritic recordings to the LEC axon terminals and the biophysical properties of the distal dendrites. Taken together, our results indicate that glutamatergic LEC inputs are capable of driving suprathreshold dendritic spikes in CA1 PN distal dendrites. The complete lack of LEC-driven action potentials recorded in over 50 somata (**Figure 1**) suggests that these local dendritic spikes do not propagate forward to drive suprathreshold somatic activity.

We next explored the synaptic properties of LEC-CA1 inputs using photostimulation trains. Presynaptic short-term plasticity and postsynaptic summation could provide insight into the facilitatory versus depressing nature of LEC inputs and elucidate whether the repeated activity of LEC inputs enables the propagation of dendritic spikes to drive somatic firing (**Figures S6-7**). First, we measured the presynaptic short-term plasticity dynamics of the LEC inputs in CA1 PN somata using trains of light pulses delivered across physiologically-relevant frequencies (1-10 Hz, ≤ theta frequency) (Frank et al., 2001; Pilkiw et al., 2017), at low light intensities (2-3% of maximal strength) (**Figure S6**). Photostimulation at 4, 8, and 10 Hz was particularly effective at facilitating the synaptic inputs from the ChR2^+^ LEC axon terminals in area CA1 (**Figure S6D-E**). Simplistically, this paired pulse facilitation suggested that the glutamatergic LEC axons may have a low release probability, although we cannot rule out the contribution of shifts in excitation-inhibition balance. Nevertheless, a low release probability in glutamatergic LEC axons may explain the lack of LEC-driven action potentials in CA1 PN somata in response to a single light pulse. So, we tested whether trains of photostimulation at maximum (100%) strength could evoke summation-driven action potentials in CA1 PN somata. Surprisingly, we did not observe any LEC-driven action potentials in the CA1 PN somata, although the maximum photostimulation trains elicited significant summation of the post-synaptic responses (**Figure S7A-F**, **J**). Meanwhile, 8 Hz photostimulation at maximum strength produced significant summation of LEC inputs in the distal dendrites, and was capable of increasing dendritic spike probability (**Figure S7G-J**).

### LEC inputs recruit strong feed-forward inhibition onto both compartments of CA1 PNs

What are the local circuit mechanisms underlying LEC-driven dendritic spikes in CA1 PN distal dendrites? Given that we observe LEC-driven dendritic spikes when inhibition is intact, could GABAergic inhibition be actively gating the incidence of distal dendritic spikes or their spread to the soma? GABAergic inputs synapse onto the entire somato-dendritic axis of CA1 PNs (Glickfeld et al., 2009; Megias et al., 2001) and can thus shape compartment-specific activity (Bloss et al., 2016; Pouille and Scanziani, 2001). To assess the role of feed-forward inhibition (FFI) in regulating LEC-driven depolarization and coupling between the dendrite and soma, we pharmacologically blocked GABA_A_ and GABA_B_ receptors with bath application of antagonists SR95531 (2 μM) and CGP55845 (1 μM) (**Figure 3A, F**) while holding the CA1 PN at resting membrane potential. We directly measured the LEC-driven excitatory post-synaptic potential (EPSP) from the dendrite or soma in current clamp mode, and derived the underlying inhibitory PSP (inferred IPSP) for subthreshold responses (Basu et al., 2013; Basu et al., 2016; Pouille and Scanziani, 2001). The inferred IPSP was calculated by subtracting the EPSP (measured after GABA receptor blockade) from the net PSP (EPSP+IPSP; measured with inhibition intact in the control condition) (**Figure 3B, G**). This strategy was necessary to specifically investigate LEC-mediated FFI onto CA1 PN dendrites. Synaptic voltage clamp is not effective in dendrites (Beaulieu-Laroche and Harnett, 2018; To et al., 2021), so isolating inhibitory post-synaptic currents (IPSCs) by holding the dendrite at a membrane potential close to AMPA/NMDA reversal potential (0 to +10 mV) in voltage clamp is not feasible. Previous studies and computational modeling have validated that this approach gives a reliable estimate of the IPSP at the somatic and dendritic compartment (Basu et al., 2013; Basu et al., 2016).

**Figure 3.**
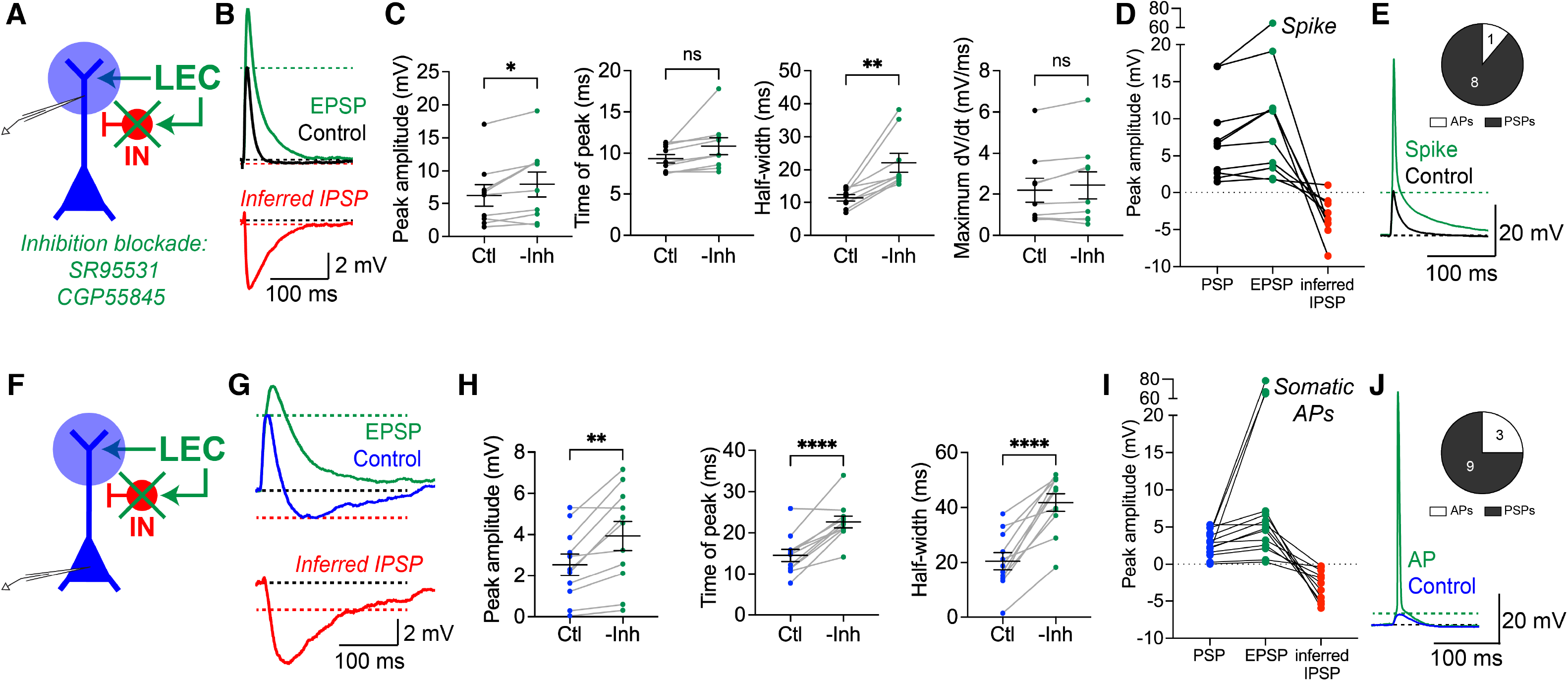
LEC inputs recruit strong feed-forward inhibition onto the somatic and dendritic compartments of CA1 PNs. (A) Experimental strategy. Photostimulation (blue circle) of ChR2^+^ LEC axons (green arrow) in SLM during whole-cell recordings from a CA1 PN dendrite. SR95531 and CGP55845 were perfused onto the brain slice to block inhibition (green X). (B) Example traces from a current clamp recording of an LEC-driven PSP in a CA1 PN dendrite before and after pharmacological inhibition blockade. The control PSP (black) was recorded in drug-free conditions, followed by an EPSP (green) recorded after inhibition blockade. Post-hoc subtraction of the two traces resulted in an LEC-driven *inferred IPSP* trace (red). (C) LEC-driven responses in CA1 PN distal dendrites before and after inhibition blockade. From left to right: Peak amplitude (p = 0.025), Time of peak (p = 0.0612), Half-width (p = 0.0022), Maximum dV/dt (p = 0.0866). Paired t test, n = 9 CA1 PN dendrites. (D) Peak amplitude of the LEC-driven responses per recorded dendrite. For subthreshold responses (n = 8), PSP, EPSP, and *inferred IPSP* amplitudes are shown. For suprathreshold responses (n = 1), only PSP and spike amplitudes are shown. Total n = 9 dendrites, 9 slices, 5 mice. (E) Example traces of LEC-driven post-synaptic responses before (subthreshold, black) and after (suprathreshold, green) inhibition blockade recorded in current clamp mode in a CA1 PN dendrite. Pharmacological inhibition blockade can lead to LEC-driven dendritic spikes. Inset: Proportion of dendrites where we recorded an LEC-driven spike upon removal of inhibition. Total n = 9 (1 dendrite exhibited dSpikes, 8 dendrites exhibited dPSPs). (F) Experimental strategy. Photostimulation (blue circle) of ChR2^+^ LEC axons (green arrow) in SLM during whole-cell recordings from a CA1 PN soma. SR95531 and CGP55845 were perfused onto the brain slice to block inhibition (green x). (G) Example traces from a current clamp recording of an LEC-driven PSP in a CA1 PN soma before and after pharmacological inhibition blockade. The control PSP (blue) was recorded in drug-free conditions, followed by an EPSP (green) recorded after inhibition blockade. Post-hoc subtraction of the two traces resulted in an LEC-driven *inferred IPSP* trace (red). (H) LEC-driven responses in CA1 PN somata before and after inhibition blockade. From left to right: Peak amplitude (p = 0.0023), Time of peak (p < 0.0001), Half-width (p < 0.0001). Paired t test, n = 11 CA1 PN somata. (I) Peak amplitudes of LEC-driven responses in recorded somata. For subthreshold responses (n = 11), PSP, EPSP and *inferred IPSP* amplitudes are shown. For suprathreshold responses (n = 3), only PSP and action potential amplitudes are shown. Total n = 12 CA1 PN somata, 12 slices, 9 mice. (J) Example traces of LEC-driven post-synaptic responses before (subthreshold, blue) and after (suprathreshold, green) inhibition blockade, recorded in current clamp mode in a CA1 PN soma. Pharmacological inhibition blockade can lead to LEC-driven somatic action potentials. Inset: Proportion of cells that fired an LEC-driven action potential upon removal of inhibition. Total n = 12 (3 somata exhibited APs, 9 somata exhibited PSPs).

As expected, blocking GABA receptors demonstrated that LEC-driven FFI sculpts LEC-driven responses in both neuronal compartments (**Figure 3C, H**). After removing inhibition, LEC-driven PSPs exhibited significantly higher amplitudes and significantly wider half-widths in both neuronal compartments (dendrites: 27% increase in peak amplitude (p = 0.025), 95% increase in half-width (p = 0.002); somata: 55% increase in peak amplitude (p = 0.0023), 105% increase in half-width (p < 0.0001), paired t-test, n = 9 dendrites and 11 somata) (**Figure 3C, H**). Moreover, the time of the peak of the PSPs showed significant delays in CA1 PN somata (56% increase in time to peak, p < 0.0001) (**Figure 3H**). There were no significant changes to the time of peak or maximum dV/dt of PSPs recorded at the dendrite (time of peak p = 0.0612, maximum dV/dt p = 0.087) (**Figure 3C**). While the amount of inhibition recruited by glutamatergic LEC inputs varied across recorded pyramidal neurons, on average, the inferred IPSP was comparable in CA1 PN distal dendrites (-3.33 ± 0.91 mV) and somata (-3.11 ± 0.66 mV). Thus, FFI (IPSPs) recruited simultaneously by LEC excitatory inputs did significantly curb the coincident monosynaptic excitation (EPSPs), but together resulted in a net overall depolarization LEC-driven PSPs (**Figure 3D, I**). Furthermore, removing inhibition enabled LEC-driven action potentials in CA1 PN somata in some cases (25% of somata, 11% of dendrites, **Figure 3E, J**), pointing to a complex interaction between the excitation-inhibition balance within the circuit and attenuation of dendritic signals. Together, these results demonstrate that inhibition plays a prominent role in the LEC-to-CA1 circuit. Glutamatergic LEC inputs drive strong feed-forward inhibition onto the CA1 PN, which limits the degree and length of depolarization in the dendrites and somata, thereby sculpting excitability and coupling dynamics between both neuronal compartments.

### LEC inputs directly recruit VIP and CCK interneuron populations in CA1

After demonstrating that inhibition can shape LEC-driven responses in CA1 PNs, we wanted to identify specific GABAergic microcircuit(s) that were likely to modulate LEC-driven dendritic excitability. There are over 20 types of local INs in hippocampal area CA1, each with distinct inputs and physiological functions (Bezaire and Soltesz, 2013; Geiller et al., 2020; Klausberger and Somogyi, 2008; Pelkey et al., 2017). We investigated three genetically defined (Taniguchi et al., 2011) candidate IN populations in CA1 that were likely to be targeted by LEC projections and could feasibly modulate LEC-driven dendritic activity. These were the vasoactive intestinal polypeptide (VIP)-expressing INs and cholecystokinin (CCK)-expressing INs, whose somata and dendritic arbors are located in SLM where the LEC axons terminate (Acsady et al., 1996a; Basu et al., 2013; Basu et al., 2016; Cope et al., 2002; Francavilla et al., 2015; Klausberger, 2009; Pawelzik et al., 2002; Tyan et al., 2014), as well as the somatostatin (SST)-expressing INs, which classically modulate the dendritic activity of pyramidal neurons (Leao et al., 2012; Lovett-Barron et al., 2014; Lovett-Barron et al., 2012). Although parvalbumin (PV)-expressing interneurons are known to mediate FFI within area CA1, they were not investigated because they classically modulate somatic and axonal activity (Dudok et al., 2021a; Pawelzik et al., 2002; Sik et al., 1995; Takacs et al., 2015) and because there is mixed evidence on whether PV INs receive input from the entorhinal cortex (Dudok et al., 2021b; Milstein et al., 2015).

To test whether the three candidate IN populations in CA1 are recruited by glutamatergic LEC inputs, we photostimulated glutamatergic LEC axons, as previously described, and performed targeted patch-clamp recordings from tdTomato-labeled VIP, CCK, or SST INs in hippocampal slices derived from IN-specific transgenic mouse lines. IN-specific tdTomato labeling was achieved by crossing transgenic VIP-Cre, SST-Cre, or CCK-Cre/Dlx-Flp (Basu et al., 2013; Taniguchi et al., 2011) driver mouse lines with the appropriate recombinase-dependent tdTomato reporter lines. This circuit mapping strategy allowed us to systematically probe whether LEC inputs recruit particular candidate interneuron populations in area CA1.

TdTomato-positive VIP INs in CA1 were located primarily in SLM, where LEC axons innervate, and in *stratum pyramidale* (SP) (**Figure 4A-B** (top), **Figure S8A**). We locally photostimulated the ChR2^+^ LEC axons in SLM and recorded from tdTomato^+^ VIP IN somata across the SLM and SP layers under current clamp mode. Light-evoked responses persisted in the presence of TTX and 4-AP in all VIP INs tested (100%, n = 8 VIP INs, **Figure 4E** (top)), confirming that glutamatergic LEC inputs indeed monosynaptically recruit VIP INs in area CA1. Recordings under drug-free conditions revealed robust LEC-driven excitation of VIP INs, with 20% of the recorded VIP INs exhibiting LEC-driven action potentials (93.6% VIP INs responded, 20.5% of them driven to spike, Total n = 47 VIP INs, **Figure 4F** (top), **G-I**,). Notably, the majority of VIP INs driven to spike by glutamatergic LEC inputs were located in the SLM layer (**Figure S8C**).

**Figure 4.**
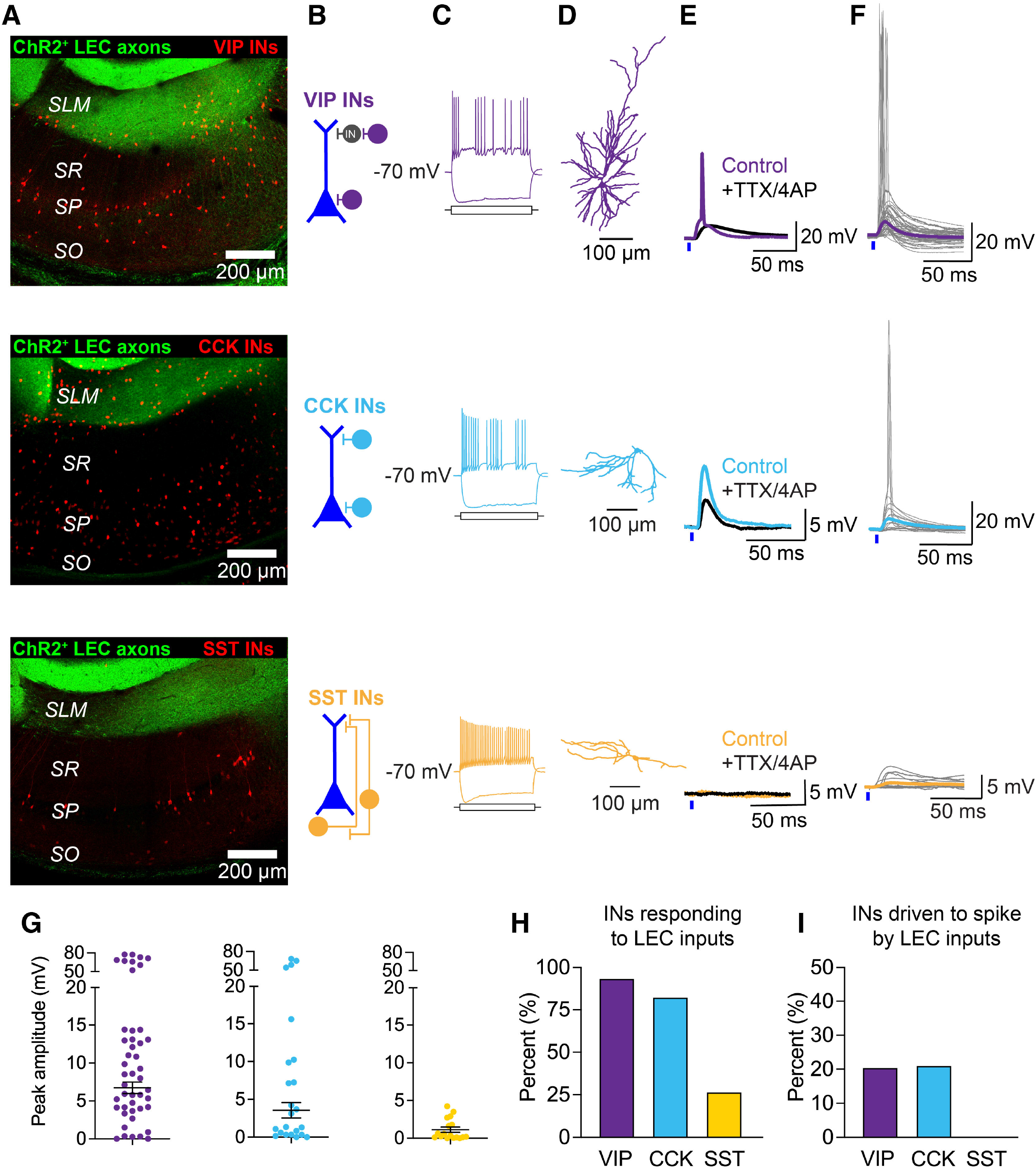
LEC inputs directly recruit VIP IN and CCK IN populations in CA1. Top to bottom: VIP INs (purple), CCK INs (light blue), SST INs (yellow). (A) Confocal images of hippocampal area CA1 with ChR2^+^ LEC axons (green) and tdTomato-labeled inhibitory neurons (red). Top to bottom: VIP-Cre/Ai14, CCK-Cre/Dlx-Flp/Ai65, SST-Cre/Ai14 mouse. (B) Schematic illustrating the predominant location of the recorded IN somata in CA1. (C) Electrophysiological signature of the example IN. Firing and sag properties were measured in response to depolarizing and hyperpolarizing current injections into the IN soma, respectively. Neurons held at -70 mV. VIP IN: +175 pA and -150 pA, CCK IN: +100 pA and -200 pA, SST IN: +175 pA and -250 pA somatic current injection. (D) Reconstruction of the recorded example IN. (E) Example traces of LEC-driven activity in the example IN before (color) and after (black) application of tetrodotoxin (TTX) and 4-aminopyridine (4-AP). (F) LEC-driven post-synaptic responses recorded in current clamp mode in response to a 2 ms photostimulation of LEC axons with 470 nm light over SLM. Traces from individual somata shown in gray. Average PSP trace shown in color. n = 46 VIP INs, 23 CCK INs, and 15 SST INs. (G) Left to right: Scatterplots of all LEC-driven post-synaptic response amplitudes from VIP INs, CCK INs, and SST INs recorded in area CA1. LEC-driven PSP amplitude mean ± SEM shown in black. (H) Summary graph demonstrating the percent of recorded INs that responded to photostimulation of glutamatergic LEC axons. 44/47 VIP INs, 19/23 CCK INs, and 4/15 SST INs responded to photostimulation of LEC axons. (I) Summary graph demonstrating the percent of responding INs that exhibited LEC-driven action potentials. Of the INs that responded to photostimulation of LEC axons, 9/44 VIP INs, 4/19 CCK INs, and 0/4 SST INs spiked.

TdTomato-positive CCK INs in CA1 were located around the SR/SLM border and in SP (**Figure 4A-B** (middle), **Figure S8A**). We recorded from tdTomato^+^ CCK IN somata across these regions and confirmed that glutamatergic LEC inputs monosynaptically recruit CCK INs in area CA1 (**Figure 4E** (middle)). Under drug-free conditions, the majority of CCK INs responded to photostimulation of LEC inputs, with 21% of the recorded CCK INs exhibiting LEC-driven action potentials (82.6% CCK INs responded, 21.1% of them driven to spike, Total n = 23 CCK INs, **Figure 4F** (middle), **G-I**). Interestingly, the CCK INs driven to spike by glutamatergic LEC inputs were located exclusively at the SR/SLM border region in area CA1 (**Figure S8C**). These belong to the interneuron subpopulation that has been shown to mediate feed-forward inhibition onto CA1 PN dendrites, and that is targeted by long-range GABAergic projections from LEC (Basu et al., 2016).

In contrast to the other candidate INs, tdTomato-positive SST INs in CA1 were found almost exclusively in *stratum oriens* (SO) and SP (**Figure 4A-B** (bottom), **Figure S8A**) and were not monosynaptically recruited or driven to spike by the photostimulation of LEC inputs (26.7% SST INs responded, 0% of them driven to spike, Total n = 15 SST INs, **Figure 4E-F** (bottom), **G-I**, **Figure S8B-C**). SST INs in CA1 fall into two major categories: the oriens-lacunosum moleculare INs (OLM INs), which are powerful mediators of feedback inhibition (Leao et al., 2012), and the bistratified SST INs, which modulate Schaffer Collateral-driven FFI (Lovett-Barron et al., 2012). The OLM INs are recruited by spiking CA1 PNs and inhibit the distal dendrites of CA1 PNs. Given that glutamatergic LEC inputs do not recruit CA1 SST INs, and do not drive CA1 PNs to spike, our results thus eliminated SST INs from the pool of candidate INs that are likely to modulate the generation of local LEC-driven dendritic spikes.

Taken together, our results highlight that VIP-expressing and CCK-expressing interneuron populations in hippocampal area CA1 are directly recruited by glutamatergic LEC inputs and likely contribute to shaping LEC-driven activity in the LEC-to-CA1 circuit.

### LEC inputs recruit disinhibitory and inhibitory VIP IN subpopulations in CA1

VIP INs mediate disinhibition in the hippocampus (Acsady et al., 1996b; Chamberland and Topolnik, 2012; Francavilla et al., 2018; Tyan et al., 2014) and cortex (David et al., 2007; Lee et al., 2013; Pfeffer et al., 2013; Pi et al., 2013), which poses an intriguing possibility that VIP INs may gate LEC-driven dendritic spike generation. However, studies have also uncovered that the hippocampal VIP IN population is extremely heterogeneous. The co-expression of molecular markers calretinin (CR) or CCK roughly splits the VIP INs into disinhibitory and inhibitory subpopulations, respectively (Acsady et al., 1996b; Tyan et al., 2014). Although we demonstrated that LEC inputs target VIP INs, it was unclear whether this included one or both functional subpopulations.

To investigate the recruitment of the VIP IN subpopulations, we photostimulated glutamatergic LEC axons, as previously described, and performed targeted patch-clamp recordings from either tdTomato-expressing CR^+^ VIP INs or CCK^+^ VIP INs in hippocampal slices derived from intersectional triple transgenic mice (VIP-Flp/CR-Cre/Ai65 or VIP-Flp/CCK-Cre/Ai65) (**Figure 5**). We found that glutamatergic LEC inputs recruit *both* CR^+^ VIP INs and CCK^+^ VIP INs, and are capable of driving action potential firing in each subpopulation (91.3% CR^+^ VIP INs responded, 14.3% of them driven to spike, Total n = 23 CR^+^ VIP INs; 84% CCK^+^ VIP INs responded, 33.3% of them driven to spike, Total n = 25 CCK^+^ VIP INs, **Figure 5E-F**). Some of the CCK^+^ VIP INs may belong to the more general CCK IN population, shown above to be recruited by LEC inputs (**Figure 4**). Within both VIP IN subpopulations, only the neurons found around the SR/SLM border were driven to spike by glutamatergic LEC inputs (**Figure S8**). Interestingly, the probability of driving light-evoked spikes in the CCK^+^ VIP IN subpopulation was higher than in the CR^+^ VIP IN subpopulation.

**Figure 5.**
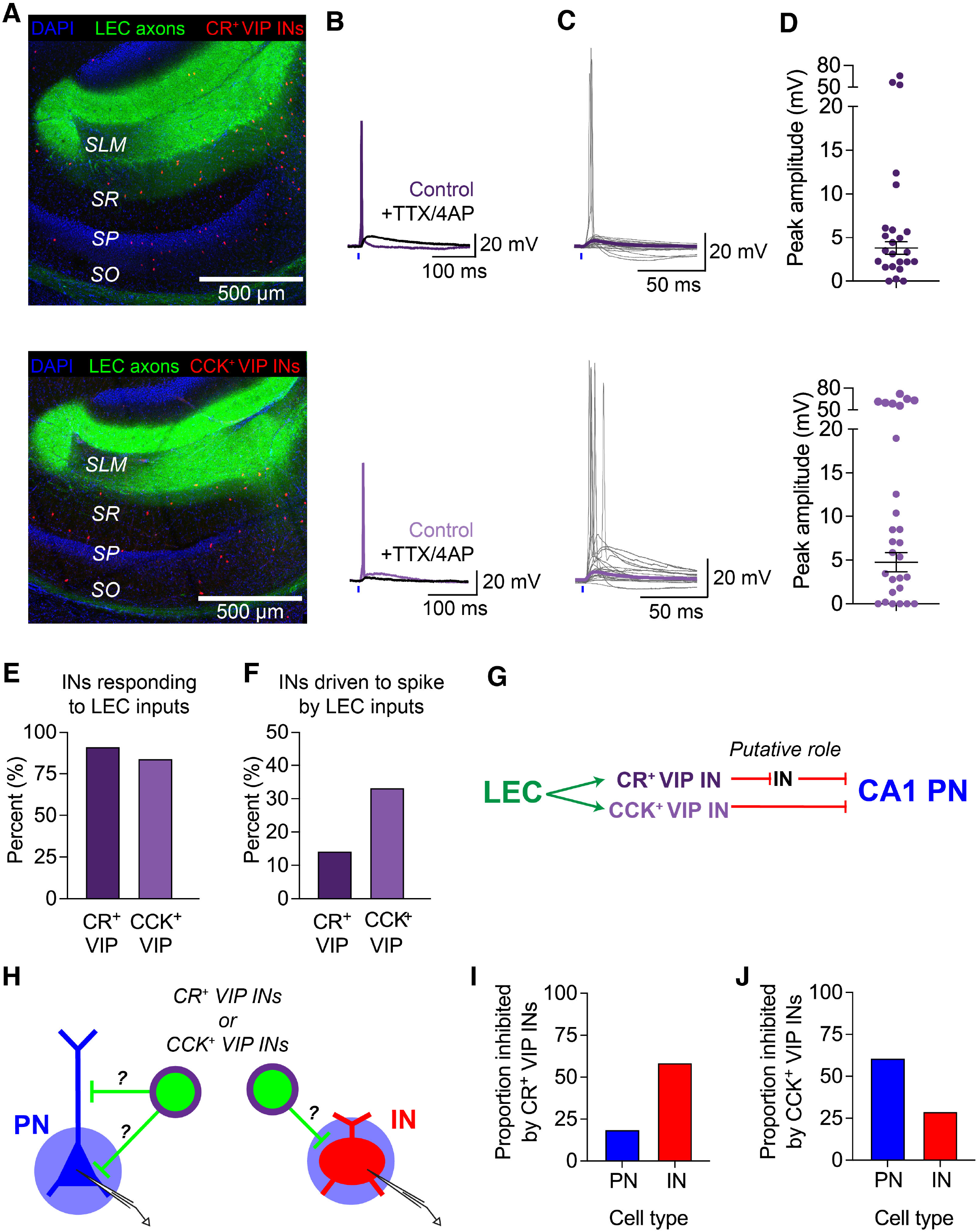
LEC inputs monosynaptically recruit both the CR^+^ VIP IN and CCK^+^ VIP IN subpopulations in area CA1. (A-D) Top row: CR^+^ VIP INs (dark purple), bottom row: CCK^+^ VIP INs (light purple). (A) Confocal images of hippocampal area CA1 with ChR2^+^ LEC axons (green) and tdTomato-labeled inhibitory neurons (red). Top: VIP-Flp/CR-Cre/Ai65 mouse, bottom: VIP-Flp/CCK-Cre/Ai65 mouse. (B) Example traces of LEC-driven activity in the VIP IN subtype before (purple) and after (black) application of tetrodotoxin (TTX) and 4-aminopyridine (4-AP). (C) LEC-driven post-synaptic responses recorded in current clamp mode in response to a 2 ms photostimulation of LEC axons with 470 nm light over SLM. Traces from individual somata shown in gray. Average PSP trace shown in purple. n = 23 CR^+^ VIP INs and 25 CCK^+^ VIP INs. (D) Scatterplots of all LEC-driven post-synaptic response amplitudes from CR^+^ VIP INs and CCK^+^ VIP INs recorded in area CA1. LEC-driven PSP amplitude mean ± SEM shown in black. (E) Summary graph demonstrating the percent of recorded INs that responded to photostimulation of glutamatergic LEC axons. 21/23 CR^+^ VIP INs and 21/25 CCK^+^ VIP INs responded to photostimulation of LEC axons. (F) Summary graph demonstrating the percent of responding INs that exhibited LEC-driven action potentials. Of the INs that responded to photostimulation of LEC axons, 3/21 CR^+^ VIP INs and 7/21 CCK^+^ VIP INs spiked. (G) Proposed circuit diagram of information flow from glutamatergic LEC neurons onto local VIP INs in area CA1, based on the literature. (H) Schematic of the experimental strategy. Voltage clamp recordings from putative pyramidal neurons and interneurons in hippocampal area CA1 from VIP-Flp/CR-Cre/Ai80 or VIP-Flp/CCK- Cre/Ai80 mice upon photostimulation of CatCh-positive CR^+^ VIP INs or CCK^+^ VIP INs. Neurons held at -80 mV to record inhibitory post-synaptic currents (IPSCs). (I) Proportion of recorded CA1 PNs and general INs that were inhibited by CR^+^ VIP INs. (J) Proportion of recorded CA1 PNs and general INs that were inhibited by CCK^+^ VIP INs.

In addition, we probed the local postsynaptic targets of CR^+^ VIP INs and CCK^+^ VIP INs to confirm whether they contribute to disinhibitory versus inhibitory downstream microcircuits. We expressed excitatory opsin CatCh (Kleinlogel et al., 2011) in the CR^+^ VIP IN or CCK^+^ VIP IN subpopulation (see STAR Methods for details) and mapped their inhibitory drive onto PNs and INs throughout area CA1, using optogenetics and voltage clamp recordings to record the presence of inhibitory post-synaptic currents (IPSCs). While each VIP IN subpopulation was found to target both PNs and INs, the CR^+^ VIP IN subpopulation predominantly targeted INs (inhibiting 60% of recorded INs vs 20% of recorded PNs; n = 29 INs, 16 PNs), whereas the CCK^+^ VIP IN subpopulation predominantly targeted PNs (inhibiting 30% of recorded INs vs 60% of recorded PNs; n = 31 INs, 23 PNs) (**Figure 5, Figure S9**).

In summary, we found that glutamatergic LEC inputs recruit various GABAergic interneuron (sub)populations with opposing functional roles. By targeting CCK INs, including the CCK^+^ VIP IN subpopulation, LEC inputs drive feed-forward inhibition in area CA1. On the other hand, by targeting VIP INs, including the CR^+^ VIP IN subpopulation, LEC inputs drive a disinhibitory microcircuit in area CA1.

### Computational modeling predicts that LEC-driven VIP INs gate excitation in CA1 pyramidal neurons

How do these local GABAergic microcircuits influence LEC-driven activity in area CA1? Thus far, we have demonstrated that glutamatergic LEC inputs can drive dendritic spikes in CA1 PNs while inhibition remains intact in the circuit (**Figure 2**). We have also shown that LEC inputs recruit local VIP INs, including the major disinhibitory subpopulation (**Figures 4, 5**). The question that arises next is whether a VIP IN-mediated disinhibitory microcircuit gates the LEC-driven dSpikes in CA1 PN distal dendrites.

We initially tested this hypothesis using a computational model. We built a multi-compartmental model of a single LEC-driven CA1 pyramidal neuron interconnected with four model neurons from the experimentally-probed candidate IN populations and tested how deleting LEC-driven INs affects dendritic spike generation (see STAR Methods for details) (**Figure 6**). Model neurons and IN connectivity features of the canonical circuit were adapted from previous modeling studies (Shuman et al., 2020; Turi et al., 2019). Biophysical and connectivity properties of all model neurons were heavily constrained by and updated to match experimental data collected in this study (**Figures S2, S4-S5, S8-S12**). The model CA1 PN received excitatory LEC inputs onto its distal dendrites and general inhibitory inputs throughout its somato-dendritic axis to provide perisomatic, proximal dendritic, and distal dendritic inhibition (**Figure 6A**). The model accurately recapitulated the incidence of the local LEC-driven dendritic spikes in control conditions (**Figure 6B**, **Figure S10D-E**), as well as the effects of inhibition blockade on LEC-driven dendritic and somatic activity (**Figure 6B**, **Figure S11**). Simulations of compartment-specific inhibition blockade further revealed that distal dendritic inhibition significantly limits the generation of LEC-driven dendritic spikes (**Figure S11B**), while perisomatic and proximal dendritic inhibition do not have a significant effect. Our canonical circuit model specifically included one of each IN type: CR^+^ VIP IN, CCK^+^ VIP IN, CCK IN found at the SR/SLM border, and SST-expressing OLM IN (**Figure 6C**, **Figure S12**). Experimental data (**Figure 4, S8**) were used to inform the recruitment and spiking probability of these INs by LEC inputs, so the OLM IN was not driven to fire action potentials. Experimental and published data were used to inform the downstream connectivity of the candidate interneurons (**Figure 5, S9**) (Basu et al., 2016; Bezaire et al., 2016; Bezaire and Soltesz, 2013; Bloss et al., 2016; Chamberland and Topolnik, 2012; Tyan et al., 2014).

**Figure 6.**
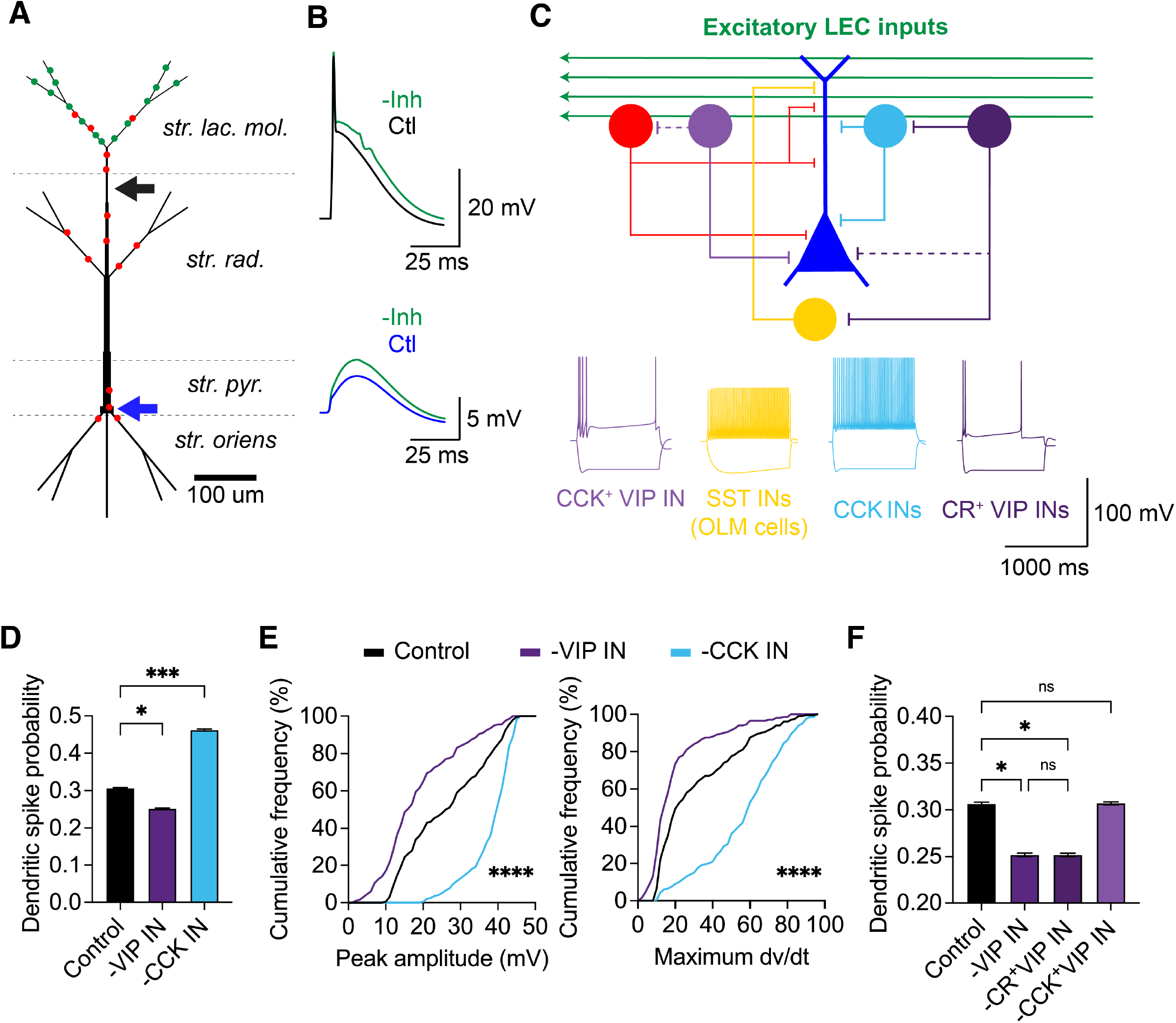
Computational modeling of silencing LEC-driven GABAergic microcircuits suggests that VIP INs disinhibit dendritic activity. (A) Schematic of the multi-compartment model of a CA1 pyramidal neuron. Excitatory synapses from glutamatergic LEC axons are found in SLM, shown in green. Inhibitory synapses from general local INs are found throughout the somato-dendritic axis, shown in red. Dendritic recording site marked by black arrow (300 µm from soma). Somatic recording site marked by blue arrow. (B) Simulated example traces of LEC-driven responses in the dendrite (black) and soma (blue) in control conditions and with inhibition removed (green), demonstrating that the model can recapitulate local dendritic spikes that do not propagate to the soma to cause action potentials. (C) Top: Schematic of model connectivity. LEC-driven CA1 pyramidal neuron surrounded by local canonical GABAergic microcircuitry: general feed-forward inhibition (red), CCK^+^ VIP INs (light purple), SST OLM INs (yellow), SR/SLM border CCK INs (light blue), and CR^+^ VIP Ins (dark purple). Recruitment of GABAergic INs by glutamatergic LEC inputs is modeled after data from Figures 3, 4, S8, S9. Weaker and/or less common connections shown as dotted lines. Bottom: Electrophysiological signatures of the canonical circuit INs generated by the model. CCK^+^ VIP IN: +250 and -200 pA, SST IN: +250 and -200 pA, CCK IN: +350 and -300 pA, CR^+^ VIP IN: +150 and -200 pA current injected into the IN soma. All IN membrane potentials held at -70 mV for measuring firing and sag properties. (D) Dendritic spike probability upon silencing LEC-driven VIP INs (purple) or CCK INs (light blue). Deletion of VIP INs (p = 0.0194), -CCK IN (p = 0.001), Dunn’s multiple comparisons test, n = 20 x 1000 trials run. (E) Cumulative frequency distributions of the peak amplitude (left) and maximum dV/dt (right) calculated for every LEC-driven dendritic spike. Peak amplitude (p < 0.0001), Maximum dV/dt (p < 0.0001), Friedman test, n = 200 trials. (F) Dendritic spike probability upon silencing LEC-driven VIP INs (purple), CR^+^ VIP INs (dark purple), or CCK^+^ VIP INs (light purple). Control versus -VIP IN (p = 0.0194), Control versus - CR^+^ VIP IN (p = 0.0194), Control versus -CCK^+^ VIP IN (p = 0.7186), -VIP IN versus -CR^+^ VIP IN (p = 0.99), Dunn’s multiple comparisons test, n = 20 x 1000 trials run.

First, we simulated the deletion of the entire VIP IN population, comprising of both CR^+^ and CCK^+^ VIP IN subpopulations. Compared to control conditions, VIP IN deletion led to a significant decrease in LEC-driven dSpike probability (p = 0.0194, Dunn’s multiple comparisons test, n = 20 trials, 1000 runs per trial, **Figure 6D**), revealing that LEC overall engages predominantly the disinhibitory function of VIP INs in area CA1 dendrites. The peak amplitude and maximum dV/dt values of dSpikes were similarly affected (peak amplitude: p < 0.0001, maximum dV/dt: p < 0.0001, Friedman test, n = 200 trials, **Figure 6E**, **Figure S13**). Meanwhile, CCK IN deletion had the opposite effect on dSpike probability (p = 0.001, **Figure 6D, Figure 6E**, **Figure S13**), amplitude, and maximum dV/dt values (peak amplitude: p < 0.0001, maximum dV/dt: p < 0.0001), consistent with the idea that CCK INs found in the SR/SLM border region mediate cortically-driven feed-forward inhibition onto CA1 PN distal dendrites (Basu et al., 2016).

Next, we simulated the selective deletion of CR^+^ VIP INs or CCK^+^ VIP IN subpopulations to parse the influence of the disinhibitory and inhibitory microcircuits on LEC-driven dendritic spike generation. CR^+^ VIP IN deletion led to a significant decrease in dSpike probability (p = 0.0194), whereas CCK^+^ VIP IN deletion produced no significant change (p = 0.7186, Dunn’s multiple comparisons test, n = 20 trials, 1000 runs per trial, **Figure 6F**). This result suggested that the effect of LEC-driven disinhibition stems from the activity of the CR^+^ VIP IN subpopulation in the model. CCK^+^ VIP INs have a basket cell axon morphology (Cope et al., 2002; Klausberger et al., 2005) and target the CA1 PN perisomatic region, which may limit the extent of their influence on LEC-driven suprathreshold dendritic activity. Indeed, deletion of the CCK^+^ VIP IN subpopulation resulted in a significant increase in the LEC-driven somatic PSP amplitude (**Figure S13E**), suggesting that the downstream effects of the two VIP IN subpopulations are localized to a particular neuronal compartment. Taken together, the computational modeling experiments predict that local VIP INs act as a disinhibitory gate to promote local, LEC-driven dendritic spikes in CA1 PNs, while CCK INs curb dendritic excitation by LEC through FFI.

### LEC-driven VIP INs provide net disinhibition, permitting dendritic spikes and shaping dendritic activity

Given that CCK INs have been previously shown to modulate CA1 PN dendritic and somatic activity (Basu et al., 2013; Basu et al., 2016; Del Pino et al., 2017; Whissell et al., 2019), we decided to focus on validating the disinhibitory gating role VIP INs play, as predicted by our model. We did so by removing this population from the LEC-CA1 circuit using temporally precise optogenetic silencing. First, we injected a Cre-dependent Jaws rAAV into hippocampal area CA1 of VIP-Cre or VIP-Cre/Ai14 mice to express the far-red-shifted inhibitory opsin Jaws (Chuong et al., 2014) in local CA1 VIP INs. We confirmed that Jaws could consistently and effectively silence VIP INs throughout the injected region by hyperpolarizing them to prevent action potential firing (**Figure S14A-E**). This viral strategy enabled us to indiscriminately silence Jaws^+^ VIP INs in CA1, regardless of CR or CCK co-expression.

Next, we co-injected the CaMKII-driven ChR2-eYFP virus into LEC and the Cre-dependent Jaws virus into CA1 of VIP-Cre mice. This allowed us to simultaneously activate glutamatergic LEC axons with 470 nm light and silence local CA1 VIP INs with 625 nm light (**Figure 7A-C**) in a circuit-specific, temporally-precise manner. Although the excitation spectra of ChR2 and Jaws show minimal overlap (Chuong et al., 2014; Lin, 2011), we reduced the strength and duration of the 470 nm blue light stimulus to decrease the chance of cross-talk with the Jaws expressed in CA1 VIP INs. We used this dual-color optogenetic strategy and dendritic patch-clamp recordings from CA1 PN distal dendrites to test our hypothesis that local VIP INs provide a disinhibitory gate to permit LEC-driven dendritic spikes. Each trial (30 seconds) contained two types of photostimulation conditions, separated by 15 seconds (**Figure 7D**). In the control condition, we activated the ChR2^+^ LEC axons only with 470 nm light (0.2 ms light pulse at 20-90% maximum photostimulation strength). This was followed by the VIP IN silencing condition, where we combined the activation of LEC axons with 470 nm light and silencing of Jaws^+^ VIP INs with 625 nm light (100 ms, 100% maximum photostimulation strength). The 625 nm light pulse began 50-60 ms prior to the start of the 470 nm photostimulation to ensure maximum hyperpolarization of Jaws^+^ VIP INs during the LEC-driven light-evoked response in the CA1 PN (**Figure 7D**, **Figure S14F-H**). The 625 nm light pulse ended with a 100 ms-long down ramp, to prevent rebound firing of the VIP INs (Chuong et al., 2014). Given the location of the glutamatergic LEC axons (**Figure 1**) and LEC-driven VIP INs (**Figure 4**, **S8**), we photostimulated with both wavelengths over SLM, as before (**Figure 7B-C**).

**Figure 7.**
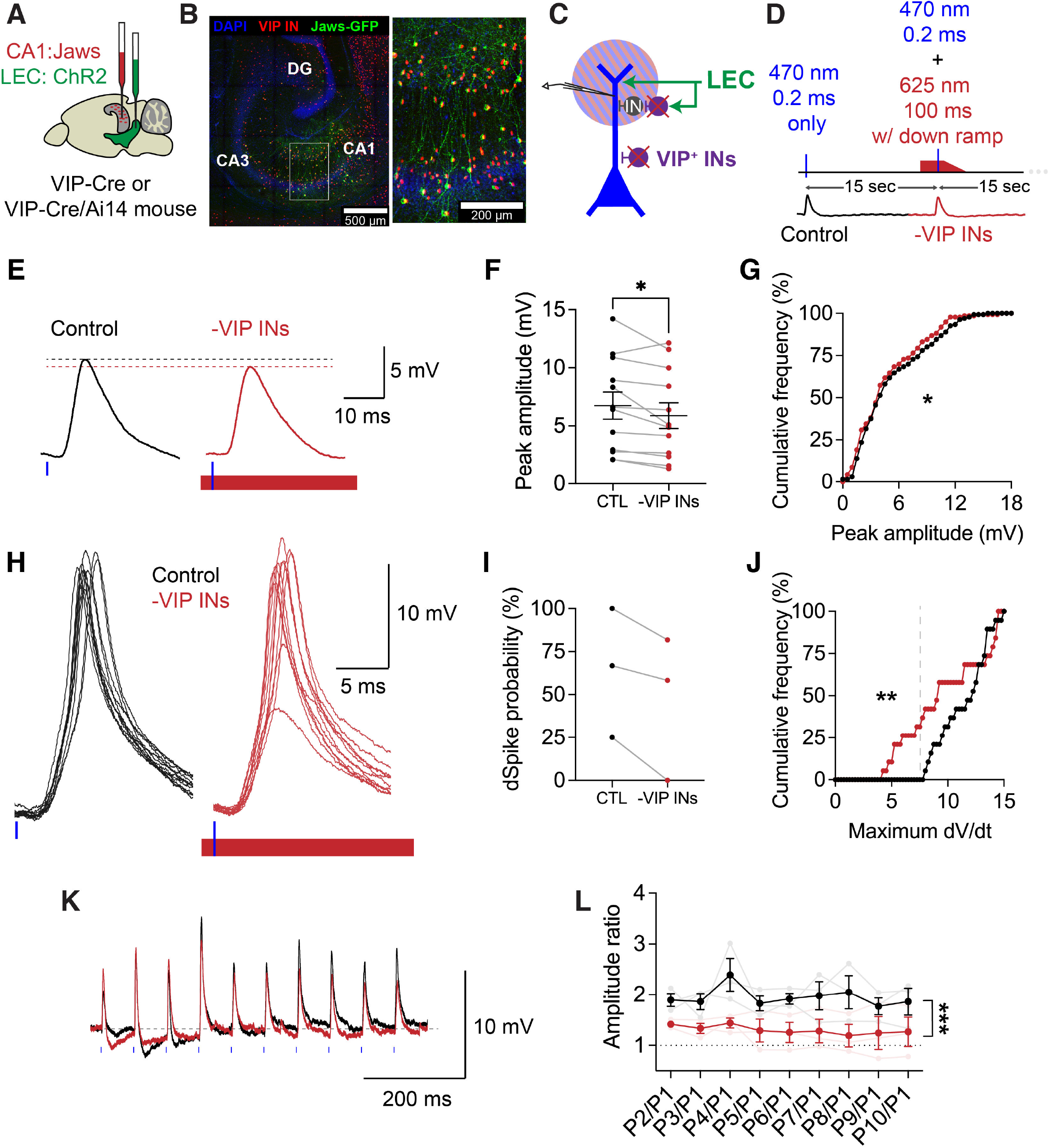
Optogenetic silencing of VIP INs confirms that LEC-driven VIP INs disinhibit dendritic spikes in CA1 PNs. (A) Schematic of the mouse brain, illustrating the co-injection of Jaws in CA1 (red) and ChR2 in LEC (green), injected into a VIP-Cre or VIP-Cre/Ai14 mouse. (B) Confocal image of the hippocampus of a VIP-Cre/Ai14 mouse, demonstrating the CA1 injection site (left) and subsequent co-expression of tdTomato^+^ (red) and Jaws-GFP (green) in VIP INs (right inset). (C) Schematic illustrating the dual color optogenetics strategy. Simultaneous photostimulation of ChR2^+^ glutamatergic LEC axons (green) with 470 nm light (blue stripes) and silencing of local Jaws^+^ VIP INs (purple neurons) in area CA1 with 625 nm light (red stripes), done during dendritic recordings from CA1 PN distal dendrites. (D) Schematic illustrating the experimental strategy. ChR2^+^ LEC axons were photostimulated (470 nm light, 0.2 ms, 20-100% maximum strength) alone or with simultaneous photostimulation of Jaws^+^ VIP INs (625 nm light, 100 ms beginning ∼50-60 ms before the 470 nm stimulus, 100% maximum strength, ending with a 100-ms-long down ramp) during the dendritic recordings. Control (black) and –VIP INs (red) stimulus conditions were separated by 15 seconds and repeated in pairs. (E) Example traces of the subthreshold LEC-driven dendritic responses (dPSPs) in a CA1 PN distal dendrite in Control (black) and -VIP INs (red) conditions. Black and red dotted lines indicate the peak amplitude of the dPSP in Control and –VIP INs conditions, respectively. (F) Average peak amplitude of the LEC-driven dPSPs in CA1 PN dendrites before and after VIP IN silencing (p = 0.0225). Paired t-test, n = 10 CA1 PN dendrites, 12 conditions, 10 slices, 8 mice. (G) Cumulative frequency distributions of the peak amplitude calculated for every subthreshold LEC-driven dendritic dPSP recorded in Control conditions (black) and the corresponding LEC-driven dendritic response recorded in the –VIP IN condition (red). Data were classified as subthreshold if the maximum dV/dt < 7.5 mV/ms in control conditions. p = 0.023, Wilcoxon matched-pairs signed-rank test. n = 10 dendrites (∼11 sweeps recorded per dendrite, on average), 12 conditions, 10 slices, 8 mice. (H) Example traces of suprathreshold LEC-driven dendritic responses (dSpikes) in a CA1 PN distal dendrite in Control (black) and -VIP IN (red) conditions. (I) Dendritic spike probability (number of dendritic spikes per number of traces recorded in the dendrite) before and after VIP IN silencing in neurons that exhibited LEC-driven dSpikes in control conditions. n = 3 dendrites, 3 slices, 2 mice. (J) Cumulative frequency distribution of the maximum dV/dt values calculated for every suprathreshold LEC-driven dSpike recorded in Control conditions (black) and the corresponding LEC-driven dendritic response recorded in the –VIP IN condition (red). Data were classified as suprathreshold if the maximum dV/dt > 7.5 mV/ms in control conditions. p = 0.0080. Wilcoxon matched-pairs signed-rank test. n = 3 dendrites (∼8 sweeps recorded per dendrite, on average), 3 slices, 2 mice. (K) Example traces demonstrating the LEC-driven responses recorded in CA1 PN dendrites in response to an 8 Hz train of light pulses at maximum (100%) photostimulation strength before (black) and after (red) VIP IN silencing. (L) Mean amplitude ratios (amplitudes of Pulse n/Pulse 1) before (black) and after (red) VIP IN silencing. p < 0.0001, Two-way ANOVA, n = 3 dendrites.

To determine the effect of VIP IN silencing on subthreshold versus suprathreshold dendritic activity, we categorized the LEC-driven light-evoked responses into dPSPs and dSpikes, based on their maximum dV/dt values (threshold dV/dt = 7.5 mV/ms, same as in **Figure 2**). Optogenetic silencing of the general VIP IN population in CA1 led to a small, yet significant decrease in dPSP amplitude, compared to the control conditions (p = 0.023, Paired t-test, n = 10 dendrites, **Figure 7E-G**, **Figure S15A-D**). There were no significant differences in dPSP kinetics (**Figure S15B**).

Notably, VIP IN silencing had a significant effect on suprathreshold activity (**Figure 7H-J**, **S15E-H**) in dendrites that exhibited LEC-driven dSpikes under control conditions. There was a consistent decrease in dendritic spike probability (**Figure 7I**), illustrated by the significant reduction in the maximum dV/dt value of LEC-driven dendritic responses after VIP IN silencing (p = 0.008, Wilcoxon matched-pairs signed-rank test, n = 3 dendrites, ∼8 sweeps recorded per dendrite, on average, **Figure 7J**, **S15E-H**). There was no significant difference in the dSpike amplitude (p = 0.495, Wilcoxon matched-pairs signed-rank test, n = 3 dendrites, **Figure S15G**). Finally, to test how VIP IN-mediated disinhibition impacts dendritic integration and summation of LEC inputs, we silenced VIP INs while photostimulating LEC inputs with 8 Hz trains. Optogenetic silencing of VIP INs led to a significant decrease in the summation of LEC-driven inputs in CA1 PN distal dendrites, compared to control conditions (p < 0.0001, Two-way ANOVA, n = 3, **Figure 7K-L**). Thus, local VIP INs in CA1 can dynamically boost the integration of repetitive dendritic inputs in CA1 PNs through disinhibition.

VIP IN silencing had no effect on LEC-driven somatic PSPs (**Figure S16**). Moreover, the level of Jaws expression correlated with the effect of VIP IN silencing seen in the dendrites but not somata (**Figure S17**), indicating that the LEC-driven VIP IN microcircuit acts in a compartment-specific manner. Thus, in line with the predictions of our model, our experiments validated our hypothesis that LEC-driven VIP INs indeed serve a disinhibitory function to promote dendritic spike generation by suppressing coincident feedforward inhibition in the distal dendrite. Our results reveal a circuit mechanism by which a dendrite-targeting, excitatory input from the lateral entorhinal cortex recruits a local disinhibitory microcircuit to drive supralinear computations in CA1 PNs, thus shaping the activity of hippocampal area CA1.

## DISCUSSION

In summary, our study demonstrated how a long-range cortical input activates a disinhibitory microcircuit in the hippocampus to modulate the generation of local dendritic spikes. Photostimulation of glutamatergic LEC axons in CA1 had asymmetric effects on the dendritic and somatic compartments of CA1 PNs. Even with inhibition intact, photostimulation of LEC inputs could drive local dendritic spikes in CA1 PNs without eliciting somatic action potentials. Additionally, we demonstrated that LEC inputs recruit strong feed-forward inhibition onto the dendrites and somata of CA1 PNs. We, therefore, hypothesized that LEC-driven circuit mechanisms gate the generation of local dendritic spikes by influencing compartment-specific excitation-inhibition balance. Targeted recordings from genetically-defined GABAergic interneurons revealed that glutamatergic LEC inputs recruit VIP INs (both CR^+^ VIP IN and CCK^+^ VIP IN subpopulations) and CCK INs. Thus, LEC inputs are poised to activate disinhibitory and inhibitory microcircuits in area CA1. A detailed computational model predicted that LEC-driven CCK INs mediate FFI to suppress dendritic spikes, whereas LEC-driven VIP INs mediate disinhibition (mostly driven by CR^+^ VIP INs) to counteract the dendritic inhibition and permit dendritic spike generation in CA1 PN distal dendrites. Optogenetic silencing of VIP INs supplied critical evidence that indeed VIP INs provide net disinhibition at the distal dendrites to gate dendritic spikes. Overall, this study uncovers a new circuit mechanism by which a single cortical input pathway can drive supralinear dendritic activity to shape single-neuron computations in the hippocampus.

### Stimulation of LEC input alone is sufficient for dendritic spike generation

A surprising but robust finding of our study is that even in the presence of inhibition, photostimulation of glutamatergic LEC inputs with a single, brief light pulse (at 100% maximum strength) elicited dendritic spikes in up to 30% of dendritic recordings (5/17 in **Figure 2**, 3/13 in **Figure 7**). A greatly-reduced photostimulation intensity (5% maximum) could still drive dendritic spikes, which suggests that LEC-driven dendritic spikes do not require maximum or repetitive activation of ChR2^+^ LEC axons within area CA1, perhaps because a VIP IN-mediated disinhibitory microcircuit, recruited in parallel, is sufficient to overcome the existing dendritic inhibition. Meanwhile, single light pulse photostimulation never elicited somatic action potentials in CA1 PNs, indicating that LEC-driven dendritic spikes are local dendritic events.

In previous *in vitro* studies, eliciting dendritic spikes in CA1 PNs typically required any combination of the following conditions: **(i)** strong electrical stimulation of the perforant path (Golding and Spruston, 1998), which recruits both excitatory and disinhibitory inputs from LEC (Basu et al., 2016) and MEC (Melzer et al., 2012), **(ii)** inhibition blockade (Golding et al., 2005; Golding et al., 2002; Lovett-Barron et al., 2012; Takahashi and Magee, 2009), **(iii)** pairing of distal and proximal dendritic inputs (Basu et al., 2013; Basu et al., 2016; Takahashi and Magee, 2009), **(iv)** high-frequency stimulation (Golding et al., 2002; Kim et al., 2015; Remy and Spruston, 2007; Takahashi and Magee, 2009), and **(v)** input stimulation combined with somatic or dendritic depolarization (Gasparini et al., 2004; Golding et al., 2002). These conditions have also been necessary to drive dendritic spikes in the cortex (Larkum et al., 2009; Stuart et al., 1997) and have been recapitulated in multiple CA1 PN modeling studies (Gomez Gonzalez et al., 2011; Pissadaki et al., 2010; Poirazi et al., 2003a, b). Thus, it was wholly unexpected that we could bypass these usual requirements for dendritic spike generation in our study. We observed, both experimentally and with modeling, that a single cortical input pathway can indeed drive dendritic spike generation while inhibition remained intact in the neural circuit. This was an exciting finding that contrasts the previously-accepted paradigms for dendritic spike generation *in vitro*, and highlights an important role for single input pathways in driving dendritic nonlinearities in principal neurons by recruiting disinhibitory microcircuits.

To speculate on their molecular composition, the majority of the LEC-driven dendritic spikes resembled simple sodium spikes (Gasparini et al., 2004; Golding and Spruston, 1998; Losonczy and Magee, 2006; Remy and Spruston, 2007), which were markedly different from the large-amplitude, long-duration plateau potentials and complex spikes with calcium and NMDA components (Basu et al., 2016; Bittner et al., 2015; Larkum et al., 2009). There was one instance of a complex, multi-peaked dendritic spike involving additional conductances, possibly calcium (d1, **Figure 2**). However, its small amplitude suggests a more distal dendritic origin and attenuation of the spike as it traveled toward our recording site. Future research, beyond the scope of this study, is necessary to determine the exact ionic nature of these LEC-driven local dendritic spikes and to verify whether they can be detected by higher-throughput imaging approaches using genetically-encoded calcium or voltage indicators that can be readily applied *in vivo*.

### A new GABAergic circuit mechanism for modulating dendritic activity

Our findings uncover new possibilities for the generation of dendritic nonlinearities. The experimental conditions used in previous studies all served to raise the dendritic membrane potential to suprathreshold levels and unleash dendritic spikes. The excitation-inhibition balance in area CA1 regulates the compartment-specific activity and neuronal computations in CA1 PNs (Glickfeld et al., 2009; Pouille and Scanziani, 2001). Dendritic inhibition limits dendritic spike generation (Larkum et al., 1999; Lovett-Barron et al., 2012; Miles et al., 1996; Murayama et al., 2009), so any mechanism that reduces this inhibition is uniquely positioned to dynamically influence nonlinear dendritic computations (Basu et al., 2016; Lovett-Barron et al., 2012). Our study demonstrates that input-driven GABAergic microcircuits can also serve this function. In our study, we recorded and modeled LEC-driven activity under more physiological conditions, with inhibition intact. These conditions allowed us to capture the downstream effects of the VIP IN-mediated disinhibitory microcircuit, which can sufficiently counteract the dendritic feed-forward inhibition to ultimately lead to dendritic spike generation. This local disinhibitory circuit motif is poised to serve as a powerful gain modulation mechanism to permit dendritic nonlinearities in a fast, dynamic, and pathway-specific manner. Local disinhibition may work in concert with long-range disinhibitory circuits such as the GABAergic projections from LEC (Basu et al., 2016) or MEC (Melzer et al., 2012) to further boost dendritic excitation of PNs.

We probed several IN populations as GABAergic microcircuit candidates for modulating glutamatergic LEC input-driven dendritic activity. We show that LEC inputs engage both feed-forward inhibitory and disinhibitory microcircuits. Among the feed-forward inhibitory microcircuits, LEC inputs excite CCK INs in the SR/SLM border region, which target CA1 pyramidal neuron dendrites (Basu et al., 2016) and the CCK^+^ VIP basket cells that target CA1 PN somata (**Figure S9E**) (Cope et al., 2002; Glickfeld and Scanziani, 2006). Thus, photoactivation of glutamatergic LEC input drives both dendritic and perisomatic inhibition. In contrast, LEC inputs also excite CCK-negative CR^+^ VIP INs that target local INs in CA1 and mediate disinhibition (**Figure S9C**) (Acsady et al., 1996b; Pelkey et al., 2017). Previous studies have identified that VIP INs inhibit SST-expressing OLM cells (Tyan et al., 2014) as well as various INs in the SR/SLM border region (Guet-McCreight et al., 2020), both of which synapse onto the distal dendritic compartment of CA1 PNs. Thus, LEC-driven VIP IN-mediated disinhibition will be most effective in counteracting dendritic inhibition. There is a third VIP IN subpopulation found within SLM, referred to as the interneuron specific type II (IS2) cells, which lack CR and CCK expression and generally target CA1 *stratum radiatum* interneurons, including CCK^+^ VIP IN basket cells (Acsady et al., 1996a; Acsady et al., 1996b; Chamberland and Topolnik, 2012; Freund and Buzsaki, 1996). Because we used the general VIP-Cre mouse line to target VIP INs, both our optogenetic mapping (**Figure 4; Figure S8** “SR/SLM VIP INs”) and silencing experiments (**Figure 7**) included the IS2 cell subpopulation. However, since IS2 cell properties are still poorly-understood, our computational model did not include them as a cell type. Therefore, our modeling results may have underestimated the net disinhibitory effect of VIP INs. On the other hand, optogenetic silencing of multiple, counteracting (disinhibitory and inhibitory) VIP IN subpopulations in area CA1 may have masked the observed disinhibitory effect upon dendritic and somatic activity. Nevertheless, our experimental and computational modeling results together suggest that while glutamatergic LEC inputs recruit a delicate balance of inhibition and disinhibition onto the CA1 PN dendrite, the VIP IN-mediated disinhibitory microcircuit aids in dendritic spike generation by boosting LEC-driven dendritic excitation.

Beyond the LEC-driven GABAergic microcircuits we uncovered, it is possible that LEC inputs excite additional interneuron populations that are trickier to access genetically, including neurogliaform cells (Price et al., 2005; Price et al., 2008) and perforant path-associated interneurons (Klausberger, 2009) found in the SLM layer (Kajiwara et al., 2008; Klausberger, 2009; Lacaille and Schwartzkroin, 1988; Pelkey et al., 2017; Price et al., 2005). Neurogliaform cells, including those expressing NPY^+^ and nNOS^+^ (Armstrong et al., 2012; Tricoire et al., 2010), mediate volume GABA transmission, and produce slower GABA_B_ receptor-mediated responses (Price et al., 2005). They are therefore more likely to influence dendritic plateau potential generation and dendritic integration at a longer time scale (Chittajallu et al., 2017; Milstein et al., 2015), rather than mediate the fast dendritic spikes we observed. Meanwhile, perforant path-associated cells are not a clearly-defined genetic or functional interneuron population. Given that they express CCK, calbindin/calretinin, and/or CB1 (Klausberger et al., 2005; Pawelzik et al., 2002), it is likely some were included in our recordings of LEC-driven CCK INs and CR^+^ VIP INs at the SR/SLM border (Klausberger et al., 2005; Pawelzik et al., 2002). Advances in our understanding of GABAergic interneuron diversity, as well as the generation of intersectional transgenic mice and perturbation tools, will allow for further inspection of other LEC-driven GABAergic microcircuits in the future.

### A novel computational tool that links dendritic nonlinearities with excitatory and inhibitory input pathways in CA1 PNs

Our experimental findings benefited greatly from the model’s predictions about the role of INs on LEC-driven activity in area CA1. Unlike studies using detailed (Bezaire et al., 2016) or highly simplified (Cutsuridis and Hasselmo, 2012; Ferguson et al., 2017; Udakis et al., 2020) CA1 microcircuit models, we opted for an intermediate level of model complexity, whereby detailed biophysical profiles were inserted into reduced neuronal morphologies that accounted for the key dendritic features of CA1 PNs. This approach allowed us to maximize tractability and computational efficiency while also enabling a close fit to our own subcellular and cellular data. This minimized the non-biological variability and ensured a very high model accuracy. Moreover, while most CA1 PN models account for the effects of MEC inputs (Hsu et al., 2018; Mergenthal et al., 2020; Migliore et al., 2015; Poirazi et al., 2003a; Schulz et al., 2018; Tomko et al., 2021), ours is the first to account for LEC mediated effects and readily reproduces the LEC-driven dendritic spikes and LEC-driven interneuron activity observed in our experiments. Moreover, the model allowed us to implement *in silico* compartment-specific removal of inhibition (**Figure S11**) and deletion experiments for interneuron (sub)populations and their combinations (**Figures 6, S13, S18**). Despite continuous advancement in genetic tools, many of these manipulations remain extremely challenging, if not impossible, to perform experimentally. Our computational model enabled an extensive exploration into the role of specific circuit elements, which led us to the IN type most likely to gate dendritic spikes, namely the CR^+^ VIP IN, and guided our subsequent optogenetic silencing experiments. The above properties make the model a valuable tool for the community, as it can simulate difficult experiments across subcellular (e.g., ionic/synaptic conductances in dendrites), neuronal (e.g., somatic input-output patterns) and circuit levels (e.g., interactions between excitatory and inhibitory pathways), thus saving considerable time, effort, and animal lives. The mutually-beneficial experiment-modeling loop implemented in our study demonstrates how computational models can help further our understanding of dendritic nonlinearities and their control by circuit mechanisms. One limitation of our model is that we simulate only a small fraction of the neurons within a local CA1 microcircuit. However, by running hundreds of simulations with different initial conditions, we ensured the robustness of the modeling results.

### Local dendritic spikes: causes and consequences

Our distal dendritic recordings captured LEC-driven dendritic spike activity in the neuronal compartment found closest to the location of glutamatergic LEC synapses. The same photostimulation paradigm that could lead to LEC-driven dendritic spikes failed to generate action potentials in our somatic recordings (**Figure 1**). From this, we deduced that LEC-driven dendritic spikes are local and do not actively drive somatic spikes.

What causes the LEC-driven dendritic spikes to remain local? This phenomenon is likely due to attenuation along the dendritic arbor (Golding et al., 2005; Spruston et al., 1994), but the role of compartment-specific inhibitory modulation cannot be ruled out (**Figure 3J**)(Basu et al., 2016; Bloss et al., 2016; Jadi et al., 2012; Kamondi et al., 1998b; Lovett-Barron et al., 2012; Milstein et al., 2015). For example, during hippocampal theta oscillations, entorhinal cortex-driven depolarization of CA1 PN distal dendrites coincides with somatic hyperpolarization, thereby failing to generate somatic firing (Kamondi et al., 1998b).

Notably, the local nature of LEC-driven dendritic spikes does not make them functionally inconsequential. Local dendritic spikes have been reported *in vivo* both in the hippocampus and cortex (Jia et al., 2010; Kamondi et al., 1998b; Kerlin et al., 2019; Lavzin et al., 2012; Moore et al., 2017; Smith et al., 2013). In hippocampal area CA1, large-amplitude fast dendritic spikes, resembling our dendritic spike recordings, occur *in vivo* during coordinated sharp wave and theta oscillatory activity from CA3 and EC, respectively. These dendritic spikes remain locally in the dendrite, often unaccompanied by somatic action potentials (Kamondi et al., 1998a). This suggests that local dendritic spikes may serve a distinct function from somatic spike signals. Local dendritic spikes can enhance the electrical and biochemical coupling between the dendrite and soma. Dendritic nonlinearities broaden temporal windows for integration by NMDAR activation and trigger Ca^2+^ influx and intracellular signaling cascades that are important for inducing long-term plasticity at the soma (Basu et al., 2013; Dudman et al., 2007; Golding et al., 2002; Kim et al., 2015; Li et al., 2020). Beyond serving as an important cellular correlate for long-term plasticity (Gambino et al., 2014; Golding et al., 2002; Kim et al., 2015), local dendrite activity is linked to feature-selective tuning (Cichon and Gan, 2015; Jia et al., 2010; Moore et al., 2021; Smith et al., 2013). Branch-specific and soma-independent activity within dendrites can function to greatly expand the computational capacity of a single neuron (Augusto and Gambino, 2019; Branco and Hausser, 2010, 2011; Chavlis and Poirazi, 2021; Hausser and Mel, 2003; Losonczy et al., 2008; Moore et al., 2021; Poirazi et al., 2003b; Poirazi and Mel, 2001; Sinha and Narayanan, 2021; Sjostrom et al., 2008; Tzilivaki et al., 2019).

### Integration with other synaptic inputs may enable LEC inputs to drive somatic output and induce plasticity

Under what conditions could excitatory LEC inputs drive somatic spike output? High frequency, burst-like activity and/or integration of coincident inputs could elicit somatic spikes by sufficiently depolarizing the somatic compartment. We expanded our computational model, which recapitulated our experimentally-recorded local LEC-driven dendritic spikes (**Figure S10**), and demonstrated that simulating high frequency, *in vivo*-like patterns of activity from excitatory LEC inputs, together with somatic depolarization, leads to action potential firing in the CA1 PN model neuron (**Figure S18**). Notably, at higher rates of LEC input firing (e.g., 100 Hz, burst-like firing at theta peak), the somatic spike output is differentially modulated by LEC-driven candidate INs. Disinhibitory VIP INs gate action potential firing, while SR/SLM CCK INs and CCK^+^ VIP INs limit action potential firing at specific LEC input activity frequencies.

The integration of co-active inputs within CA1 PN dendrites may provide additional boosts in depolarization, aiding the forward propagation of LEC-driven dendritic spikes to drive somatic action potentials. Glutamatergic LEC projections may integrate with other inputs arriving at various compartments of the CA1 PN. These include **(i)** LEC long-range inhibitory projections that disinhibit CA1 PN dendrites by inhibiting the dendrite-targeting CCK INs at the SR/SLM border (Basu et al., 2016), **(ii)** MEC inputs (Bittner et al., 2015) arriving at the distal dendritic compartment, or **(iii)** CA3 inputs arriving at the proximal dendritic compartment (Basu et al., 2013; Basu et al., 2016; Bittner et al., 2017; Dudman et al., 2007; Golding et al., 2002; Jarsky et al., 2005; Takahashi and Magee, 2009). Due to our viral strategy and *in vitro* preparation, it is highly unlikely that these additional inputs were recruited in our experiments. The CaMKII-driven ChR2 virus was specifically injected into LEC; data collected from animals that exhibited retrograde viral spread or leaking into MEC or the hippocampus were discarded. Moreover, axons synapsing onto the distal and proximal dendrites of CA1 PNs are cut off from their respective cell bodies in EC and CA3, so our focal photostimulation paradigm would have activated only the ChR2-expressing glutamatergic LEC axon terminals found in SLM. Nevertheless, future experiments should explore the dendritic integration of glutamatergic LEC inputs with other converging inputs in area CA1 and investigate whether their spatiotemporal coordination can drive learning rules such as input-timing-dependent plasticity (ITDP) (Basu et al., 2013; Dudman et al., 2007), spike-timing-dependent plasticity (STDP) (Bi and Poo, 1998; Markram et al., 1997), or behavioral time scale dependent plasticity (BTSP) (Bittner et al., 2017; Grienberger et al., 2017). The integration of various long-range and local inputs may differentially activate local circuit elements to further boost dendritic spike generation and propagation to drive context-dependent somatic output and coding in the hippocampus.

### LEC inputs may support hippocampal memory representations *in vivo* by gating contextually-relevant sensory activity

There is sufficient evidence to speculate that LEC may provide a powerful signal to shape context-dependent coding in the hippocampus. LEC neurons code for multisensory contextual information in the environment (Basu et al., 2016; Deshmukh and Knierim, 2011; Doan et al., 2019; Hargreaves et al., 2005; Igarashi et al., 2014; Kuruvilla et al., 2020; Lee et al., 2021; Leitner et al., 2016; Li et al., 2017; Lovett-Barron et al., 2014; Tsao et al., 2013; Tsao et al., 2018; Wang et al., 2018; Wilson et al., 2013; Witter et al., 2017; Xu and Wilson, 2012) and send dense projections to the hippocampus (Basu et al., 2016; Cui et al., 2013; Li et al., 2017; Luna et al., 2019; Masurkar et al., 2017; Nilssen et al., 2018; Ruth et al., 1988; van Groen et al., 2003; Witter et al., 2017; Witter et al., 1989; Woods et al., 2018). Studies have shown that the activity of LEC and CA1 strongly synchronize during the odor sampling phase of context-place association tasks (Igarashi et al., 2014). Moreover, disinhibitory GABAergic LEC-CA1 projections are activated by behaviorally-salient stimuli and are required for precise contextual discrimination (Basu et al., 2016). *In vivo*, LEC-driven dendritic spikes may therefore serve to assign contextual salience to sensory cues in the environment. This could be especially effective if dendritic excitability and signal propagation is boosted by the co-activation of long-range glutamatergic and GABAergic LEC inputs or other brain state-dependent neuromodulatory signals (Ito and Schuman, 2007; Lee et al., 2021; Lovett-Barron et al., 2014; Palacios-Filardo et al., 2021).

Finally, our findings pose an interesting possibility that LEC inputs may influence spatial episodic memory and adaptation during navigation by driving the context-dependent formation and remapping of place cells in the hippocampus (Buzsaki and Moser, 2013; Robinson et al., 2020). Dendritic spikes have been implicated as a predictive signal for the formation, remapping, and accuracy of place fields (Bittner et al., 2015; Bittner et al., 2017; Sheffield and Dombeck, 2015) in CA1 pyramidal neuron somata. Silencing MEC, the area predominantly associated with spatial information, only moderately affects dendritic spike output (Bittner et al., 2015), place cell properties (Hales et al., 2014; Schlesiger et al., 2018), and spatial memory behavior (Kitamura et al., 2014; Suh et al., 2011), suggesting that additional circuits may be involved in spatial coding. And although LEC has been classically associated with providing non-spatial sensory information to the hippocampus (Basu et al., 2016; Deshmukh and Knierim, 2011; Hargreaves et al., 2005; Lovett-Barron et al., 2014), recent studies suggest that LEC neurons additionally display spatially-modulated activity in the form of trace cells (Tsao et al., 2013), object location (Wang et al., 2018) and reward goals (see (Save and Sargolini, 2017). In fact, LEC lesions can impair place cell remapping in hippocampal area CA3 (Lu et al., 2013). Taken together, our and others’ findings warrant further investigation to link cellular-level dendritic computations to behaviorally-relevant network dynamics supported by cortico-hippocampal interactions.

## Supplemental Figures

**Figure S1. LEC infection confirmed by confocal imaging and local recordings**

(A) Cartoon illustrating the extent of viral infection in LEC with AAV-CaMKII-ChR2-eYFP, generated from 25 representative high-resolution confocal images of patched brain slices containing LEC.

(B) Example confocal image of LEC with viral infection.

(C) Positive control: ChR2 function in a representative infected LEC neuron. Top: Time-locked photocurrents recorded in an infected LEC neuron in cell-attached mode in response to a 10 Hz train light pulses (100% LED power). Bottom: Large, excitatory photocurrent recorded in a representative infected LEC neuron in whole-cell configuration in response to a 500 ms light pulse. Enlarged photocurrent shown to the left. Photostimulation done over LEC.

(D) Negative control: Lack of viral infection in a representative non-infected CA1 pyramidal neuron. Same protocols used as in C. Photostimulation done over CA1 PN soma.

(E) Cartoon illustrating the distribution of CA1 PN somatic and dendritic recordings along the proximal-distal and radial axes of hippocampal area CA1.

**Figure S2. Intrinsic properties of CA1 PN dendrites and soma and monosynaptic drive from glutamatergic LEC inputs**

(A) Additional example traces of the electrophysiological properties of CA1 PN dendrites. Left example: +800 pA, +500 pA, and -350 pA dendritic current injection. Right example: +800 pA, +200 pA, and -275 pA dendritic current injection. The dendrites were held at -70 mV.

(B) Summary firing (left) and sag (right) properties of CA1 PN dendrites. Individual data are shown in gray, whereas means ± SEM are shown in black. Firing and sag data n = 14 dendrites.

(C) Scatterplot of the resting membrane potential of patched CA1 PN distal dendrites. Mean ± SEM: -62.73 ± 0.92 mV. n = 17.

(D) Scatterplot of the input resistances of patched CA1 PN distal dendrites. Mean ± SEM: 287.8± 39.1 MΩ. n = 17.

(E) Additional example traces of the electrophysiological properties of CA1 PN somata. Left example: +275 pA, +50 pA, and -250 pA somatic current injection. Right example: +225 pA, +125 pA, and -300 pA somatic current injection. The somata were held at -70 mV.

(F) Summary firing (left) and sag (right) properties of CA1 PN somata. Individual data are shown in light blue, whereas means ± SEM are shown in blue. Firing data n = 52, sag data n = 43.

(G) Scatterplot of the resting membrane potential of patched CA1 PN somata. Mean ± SEM: - 68.16 ± 0.32 mV, n = 51.

(H) Scatterplot of the input resistances of patched CA1 PN somata. Mean ± SEM: 142.4 ± 6.82 MΩ, n = 54.

(I) Example traces of LEC-driven activity in a CA1 PN soma before (blue) and after (dark blue) perfusion of the brain slice with Na^+^ and K^+^ channel blockers, tetrodotoxin (TTX) and 4-aminopyridine (4-AP), respectively, used to isolate monosynaptic connections between glutamatergic LEC inputs and downstream neurons.

**Figure S3. Derivative and phase plots of LEC-driven responses in CA1 PN distal dendrites**

(A) Example LEC-driven post-synaptic responses, as shown in Figure 2A.

(B) Top to bottom rows: Light-evoked response traces, Derivative traces, and Phase plots of five example dendrites, demonstrating the visible differences between LEC-driven dPSPs (shades of yellow, left) and LEC-driven dSpikes (shades of blue, right).

**Figure S4. LEC-driven responses in CA1 PN dendrites, comparison between dPSPs and dSpikes**

Comparison between LEC-driven dendritic PSPs (n = 14) and dendritic spikes (n = 5).

(A) Resting membrane potential (p = 0.1806).

(B) Recording location (p = 0.8798).

(C) Response amplitude (p = 0.0264).

(D) Time of peak (p = 0.1695).

(E) Rise time (p = 0.2112).

(F) Half-width (p = 0.0011).

(G) Peak maximum dV/dt (p = 0.0097). Unpaired t test with Welch’s correction.

(H) Relationship between the recording location, response amplitude, and maximum dV/dt value.

**Figure S5. Comparison of LEC-driven PSPs in CA1 PN dendrites versus somata**

Comparison between LEC-driven dendritic post-synaptic potentials (n = 14) and somatic post-synaptic potentials (n = 52).

(A) Light-evoked post-synaptic potentials recorded in CA1 PN dendrites (left) and somata (middle). Individual cells’ data shown in gray. Average traces (right) shown in bold black or blue. (B-G) Compartment-specific differences in LEC-driven PSPs in the dendrite (D) versus soma (S).

(B) Peak amplitude (p < 0.0001).

(C) Time of peak (p < 0.0001).

(D) Rise time (p < 0.0001).

(E) Half-width (p < 0.0001).

(F) Hyperpolarization amplitude (p = 0.0927).

(G) Input resistance (p = 0.0001). Mann Whitney test.

**Figure S6. Short-term plasticity dynamics of LEC inputs onto CA1 PN soma**

(A) Schematic of experimental strategy. Trains of low stimulation strength photostimulation (2-3% maximum photostimulation strength), given at various physiological frequencies (1-10 Hz), were applied over ChR2^+^ LEC axons in SLM during whole-cell recordings from CA1 PN somata.

(B) Example traces of LEC-driven responses in CA1 PN somata.

(C) Average paired pulse ratios (PPR) at every frequency. PPR calculated as the ratio of Pulse n amplitude (Pn) to Pulse 1 amplitude (P1), as measured from the membrane potential immediately preceding each pulse. Sample sizes: 1 Hz n = 5, 2 Hz n = 5, 4 Hz n = 9, 8 Hz n = 9, and 10 Hz n = 15 CA1 PN somata.

(D) P2/P1 PPR. 1 Hz: p = 0.176, 2 Hz: p = 0.165, 4 Hz: p = 0.036, 8 Hz: p = 0.018, 10 Hz: p = 0.011, One sample t test comparing mean PPR to hypothetical mean = 1.

(E) P5/P1 PPR. 1 Hz: p = 0.094, 2 Hz: p = 0.034, 4 Hz: p = 0.047, 8 Hz: p = 0.037, 10 Hz: p = 0.001, One sample t test, comparing mean PPR to hypothetical mean = 1.

**Figure S7. Summation of LEC inputs in CA1 PN somata and dendrites**

(A) Schematic of experimental strategy. Trains of maximum strength photostimulation (100%), given at various physiological frequencies (2-10 Hz), were applied over ChR2^+^ LEC axons in SLM during whole-cell recordings from CA1 PN somata.

(B) Example traces of LEC-driven responses in CA1 PN somata.

(C) Average amplitude ratios at every frequency. Amplitude ratios are calculated as the ratio of Pulse n amplitude (Pn) to Pulse 1 amplitude (P1) as measured from the membrane potential immediately preceding Pulse 1. Sample sizes: 2 Hz n = 13, 4 Hz n = 14, 8 Hz n = 14, and 10 Hz n = 13 CA1 PN somata.

(D) P2/P1 amplitude ratios. 2 Hz: p = 0.0873; 4 Hz: p = 0.3329; 8 Hz: p = 0.0363; 10 Hz: p = 0.1397. One sample t test, comparing mean amplitude ratio to hypothetical mean = 1.

(E) P10/P1 amplitude ratios. 2 Hz: p = 0.0002; 4 Hz: p = 0.0583; 8 Hz: p = 0.0006; 10 Hz: p = 0.0002. One sample t test, comparing mean amplitude ratio to hypothetical mean = 1.

(F) Peak amplitudes at every frequency, illustrating the lack of LEC-driven action potentials in CA1 PN somata, even after activation with trains of photostimulation at maximum photostimulation strength. Individual data shown in faded blues. Average data shown in full opacity blues. Sample sizes: 2 Hz n = 13, 4 Hz n = 14, 8 Hz n = 14, and 10 Hz n = 13 CA1 PN somata.

(G) Schematic of experimental strategy. Trains of maximum strength photostimulation (100%), given at 8 Hz, were applied over ChR2^+^ LEC axons in SLM during whole-cell recordings from CA1 PN dendrites.

(H) Example trace of LEC-driven responses in a CA1 PN distal dendrite. Squares indicate dendritic spikes.

(I) Amplitude ratios. P2/P1: p = 0.1217; P10/P1: p = 0.0460. One sample t test, comparing mean amplitude ratio to hypothetical mean = 1, n = 5 dendrites.

(J) Summary graph demonstrating the LEC-driven spike probability in somata (blue) and dendrites (black) during a single 2 ms (circle) and 8 Hz photostimulation train (diamonds), when protocols were given at maximum (100%) photostimulation strength. Action potential probability was calculated for soma, whereas dendritic spike probability was calculated for dendrites. Dendrites, single stimulation: n = 17; Dendrites, 8 Hz train: n = 5; Somata, single stimulation: n = 50, Somata, 8 Hz train: n = 19.

**Figure S8. Interneuron subgroups in hippocampal area CA1, as defined by soma location**

(A) Left: Cartoon illustrating the distribution of recorded CA1 INs along the proximal-distal and radial axes of hippocampal area CA1. Right: Distribution of molecularly-defined GABAergic interneurons (VIP INs, CCK INs, SST INs; CR^+^ VIP INs, CCK^+^ VIP INs) in hippocampal area CA1, based on the quantification of tdTomato-labeled cells in SLM, SR, SP, and SO, scaled by imaging volume.

(B) Summary graph demonstrating the percent of INs that responded to photostimulation of glutamatergic LEC axons. Data categorized by the location of the interneuron soma in the CA1 strata. 18/20 SP VIP INs, 15/15 SLM VIP INs, 5/9 SP CCK INs, 14/14 SR/SLM border CCK INs, 1/10 SO SST INs, 3/4 SP SST INs; 6/7 SP CR^+^ VIP INs, 15/16 SR/SLM CR^+^ VIP INs, 6/9 SP CCK^+^ VIP INs, 15/16 SR/SLM CCK^+^ VIP INs responded to photostimulation of LEC axons.

(C) Summary graph demonstrating the percent of responding INs that exhibited LEC-driven action potentials in area CA1. Data categorized by the location of the interneuron soma in the CA1 strata. Of the INs that responded to photostimulation of LEC axons, 1/18 SP VIP INs, 8/15 SLM VIP INs, 0/5 SP CCK INs, 4/14 SR/SLM border CCK INs, 0/1 SO SST IN, 0/3 SP SST INs; 0/6 SP CR^+^ VIP INs, 3/15 SR/SLM CR^+^ VIP INs, 0/6 SP CCK^+^ VIP INs, 7/15 SR/SLM CCK^+^ VIP INs spiked.

**Figure S9. Local VIP IN subtypes exhibit disinhibitory and inhibitory functions**

(A) Schematic of the experimental strategy. Voltage clamp recordings from putative pyramidal neurons and interneurons in hippocampal area CA1 upon photostimulation of CatCh-positive CR^+^ VIP INs or CCK^+^ VIP INs. Neurons held at -80 mV to record inhibitory post-synaptic currents (IPSCs).

(B) Proportion of patched CA1 PNs and layer-specific INs that were inhibited by CR^+^ VIP INs.

(C) CR^+^ VIP IN-driven IPSCs from recorded CA1 PNs (left) and interneurons found in SLM, SR, and SO (right). Traces from individual neurons shown in gray. Average IPSC trace shown in blue (PN) or red (INs) with the fraction of responding cells noted on the top right.

(D) Proportion of patched CA1 PNs and layer-specific INs that were inhibited by CCK^+^ VIP INs.

(E) CCK^+^ VIP IN-driven IPSCs from recorded CA1 PNs (left) and interneurons found in SLM, SR, and SO (right). Traces from individual neurons shown in gray. Average IPSC trace shown in blue (PN) or red (INs) with the fraction of responding cells noted on the top right.

**Figure S10. Model validation: CA1 PN intrinsic properties; dPSP versus dSpike delineation**

Top row: dendritic data. Bottom row: somatic data.

(A) Example representative electrophysiological signatures of the CA1 PN distal dendrite (300 μm from the soma) or soma, generated by the multi-compartment computational model of a CA1 PN. Compartment membrane potential held at -70 mV measuring of firing and sag properties.

(B) Example dendritic (top, black) or somatic (bottom, blue) firing, in response to depolarizing current steps.

(C) Example dendritic (top, black) or somatic (bottom, blue) sag, in response to hyperpolarizing current steps.

(D) Example traces of dendritic spikes generated by the computational model.

(E) Box and whisker plots (median ± interquartile range) comparing LEC-driven dendritic post-synaptic potentials (n = 200) and dendritic spikes (n = 200) generated by the computational model. From left to right: Peak amplitude (p < 0.0001), Time of peak (p < 0.0001), Half-width (p = 0.0377), Maximum dV/dt (p < 0.0001). Mann Whitney test. Plus sign indicates mean values.

**Figure S11. Simulation: compartment-specific inhibition blockade**

(A) Simulated examples of LEC-driven responses in the dendrite (black) and soma (blue) in control conditions and with compartment-specific inhibition removed (complete removal in green, somatic removal in yellow, proximal dendritic removal in orange, distal dendritic removal in red).

(B) Dendritic spike probability upon removal of compartment-specific inhibition. Control versus -Inh (p < 0.0001), Control versus -Soma Inh (p = 0.99), Control versus -Prox Dend Inh (p = 0.99), Control versus -Dist Dend Inh (p < 0.0001), Tukey’s multiple comparisons test, n = 50 x 1000 trials.

(C) Box and whisker plots (median ± interquartile range) of the peak amplitude (left) and half-width (right) of LEC-driven dendritic spikes recorded in control conditions and upon removal of compartment-specific inhibition. Peak amplitude: Control versus -Inh (p < 0.0001), Control versus -Soma Inh (p = 0.0003), Control versus -Prox Dend Inh (p < 0.0001), Control versus -Dist Dend Inh (p < 0.0001), Dunn’s multiple comparisons test, n = 100 trials. Half-width: Control versus - Inh (p < 0.0001), Control versus -Soma Inh (p = 0.0012), Control versus -Prox Dend Inh (p < 0.0001), Control versus -Dist Dend Inh (p = 0.0025), Dunn’s multiple comparisons test, n = 100 trials.

(D) Box and whisker plots (median ± interquartile range) of the peak amplitude (left) and half-width (right) of LEC-driven dendritic PSPs recorded in control conditions and upon removal of compartment-specific inhibition. Peak amplitude: Control versus -Inh (p < 0.0001), Control versus -Soma Inh (p < 0.0001), Control versus -Prox Dend Inh (p < 0.0001), Control versus -Dist Dend Inh (p < 0.0001), Dunn’s multiple comparisons test, n = 100 trials. Half-width: Control versus - Inh (p < 0.0001), Control versus -Soma Inh (p < 0.0001), Control versus -Prox Dend Inh (p < 0.0001), Control versus -Dist Dend Inh (p < 0.0001), Dunn’s multiple comparisons test, n = 100 trials.

(E) Box and whisker plots (median ± interquartile range) of peak amplitude (left) and half-width (right) of LEC-driven somatic PSPs recorded in control conditions and upon removal of compartment-specific inhibition. Peak amplitude: Control versus -Inh (p < 0.0001), Control versus -Soma Inh (p < 0.0001), Control versus -Prox Dend Inh (p = 0.0007), Control versus -Dist Dend Inh (p < 0.0001), Dunn’s multiple comparisons test, n = 100 trials. Half-width: Control versus - Inh (p < 0.0001), Control versus -Soma Inh (p < 0.0001), Control versus -Prox Dend Inh (p = 0.0203), Control versus -Dist Dend Inh (p < 0.0001), Dunn’s multiple comparisons test, n = 100 trials.

**Figure S12. Model validation: LEC-driven GABAergic interneurons**

(A) Electrophysiological signatures of the canonical circuit INs generated by the model. CCK IN: +350, +100, and -300, SST IN: +250, +100, and -200, CR^+^ VIP IN: +150, +75, and -200, and CCK^+^ VIP IN: +250, +100, and -200 pA current injected into the IN soma. Insets show the action potentials generated in the VIP IN subpopulations in response to the highest depolarizing current injection. Trace generated from the middle current injection is shown in a darker color. All IN membrane potentials held at -70 mV for measuring firing and sag properties.

(B) Summary graph demonstrating the percent of INs that responded to glutamatergic LEC inputs in the model.

(C) Summary graph demonstrating the percent of responding INs that spiked in the model. B and C were calculated from 10 runs of the model, with 200 trials per run.

**Figure S13. Deletions of VIP IN and CCK IN populations and VIP IN subpopulations**

(A) Schematic of model connectivity, same as Figure 7A, illustrating the deletion of CR^+^ VIP IN and/or CCK^+^ VIP INs, as well as the deletion of CCK INs (black x’s).

(B) Dendritic spike probability upon silencing LEC-driven interneurons. Control versus -VIP IN (p = 0.0194), Control versus -CR^+^ VIP IN (p = 0.0194), Control versus -CCK^+^ VIP IN (p = 0.7186), Control versus -CCK IN (p = 0.001), -VIP IN versus -CR^+^ VIP IN (p = 0.99), -VIP IN versus - CCK^+^ VIP IN (p < 0.0001), Dunn’s multiple comparisons test, n = 20 x 1000 trials.

(C) Box and whisker plots (median ± interquartile range) plots of the peak amplitude (left) and maximum dV/dt (right) calculated for every LEC-driven dendritic spike recorded in control conditions (black) and the corresponding LEC-driven dendritic responses after interneuron deletion. Data were classified as suprathreshold if the maximum dV/dt > 10 mV/ms in control conditions. Peak amplitude: Control versus -VIP IN (p = 0.99), Control versus -CR^+^ VIP IN (p < 0.0001), Control versus -CCK^+^ VIP IN (p < 0.0001), Control versus -CCK IN (p < 0.0001); Maximum dV/dt: Control versus -VIP IN (p = 0.0017), Control versus -CR^+^ VIP IN (p < 0.0001), Control versus -CCK^+^ VIP IN (p = 0.0126), Control versus -CCK IN (p < 0.0001), -VIP IN versus -CR^+^ VIP IN (p = 0.7147) Dunn’s multiple comparison’s test, n = 200 trials.

(D) Box and whisker plots (median ± interquartile range) plots of the peak amplitude (left) and maximum dV/dt (right) calculated for every LEC-driven dendritic PSP recorded in control conditions (black) and the corresponding LEC-driven dendritic responses upon interneuron deletion. Data were classified as subthreshold if the maximum dV/dt < 10 mV/ms in control conditions. Peak amplitude: Control versus -VIP IN (p < 0.0001), Control versus -CR^+^ VIP IN (p = 0.99), Control versus -CCK^+^ VIP IN (p < 0.0001), Control versus -CCK IN (p < 0.0001), -VIP IN versus -CR^+^ VIP IN (p < 0.0001), -VIP IN versus -CCK^+^ VIP IN (p = 0.99); Maximum dV/dt: Control versus -VIP IN (p = 0.2376), Control versus -CR^+^ VIP IN (p = 0.99), Control versus - CCK^+^ VIP IN (p = 0.0009), Control versus -CCK IN (p < 0.0001) , -VIP IN versus -CR^+^ VIP IN (p = 0.0295), -VIP IN versus -CCK^+^ VIP IN (p = 0.9688), Dunn’s multiple comparison’s test, n = 200 trials.

(E) Box and whisker plots (median ± interquartile range) of the peak amplitude calculated for every LEC-driven somatic response in control conditions (black) and the corresponding LEC-driven somatic responses recorded upon interneuron deletion. Control versus -VIP IN (p < 0.0001), Control versus -CR^+^ VIP IN (p = 0.99), Control versus -CCK^+^ VIP IN (p < 0.0001), Control versus -CCK IN (p < 0.0001), -VIP IN versus -CR^+^ VIP IN (p < 0.0001), -VIP IN versus -CCK^+^ VIP IN (p = 0.99), Dunn’s multiple comparison’s test, n = 200 trials.

**Figure S14. Validation of optogenetic silencing of local VIP INs with Jaws**

(A) Top: Schematic of the mouse brain, illustrating the CA1 injection site (red). A flexed Jaws AAV was injected into a VIP-Cre or VIP-Cre/Ai14 mouse. Bottom: Schematic of experimental strategy. Optogenetic silencing of local VIP INs in area CA1 (yellow-filled purple neurons) to validate Jaws as a functional inhibitory opsin. Jaws-positive VIP INs were photostimulated with 625 nm light (100% maximum stimulation strength) for 10-1000 ms durations.

(B) Confocal images of hippocampal area CA1 of a VIP-Cre/Ai14 mouse, demonstrating the infection of tdTomato^+^ VIP INs (red) with Jaws-GFP (green). White insets: Jaws infection spanned various CA1 layers.

(C) Scatterplot demonstrating the degree of hyperpolarization recorded upon photostimulation of Jaws-positive VIP INs in area CA1 with 625 nm light.

(D) Positive control: 625 nm light can effectively silence action potential firing in Jaws-positive VIP INs.

(E) Spike frequency in Jaws^+^ VIP INs during positive current injection before (purple) and after (red) photostimulation with 625 nm light. Individual data shown in faded purple and faded red. Averages shown in purple and red. n = 19 VIP INs.

(F) Example traces of Jaws-mediated hyperpolarization in an example VIP IN in response to photostimulation with different durations of 625 nm light (10 ms increments, up to 100 ms).

(G) Peak hyperpolarization amplitude upon photostimulation with different durations of 625 nm light. Gray lines showing data from individual VIP INs. Average ± SEM hyperpolarization amplitude per stimulation duration shown in red.

(H) Peak hyperpolarization amplitude, normalized to the maximum hyperpolarization recorded within the individual VIP IN in G. Average hyperpolarization amplitude per stimulation duration shown in black. To maximally hyperpolarize the Jaws-positive VIP INs, the 625 nm stimulus will begin 50-60 ms before the 470 nm stimulus to activate glutamatergic LEC axons.

**Figure S15. Effect of VIP silencing on LEC-driven subthreshold and suprathreshold dendritic responses**

(A) Example traces of the subthreshold LEC-driven dPSPs in a CA1 PN distal dendrite in Control (black) and -VIP INs (red) conditions. Black and red dotted lines indicate the peak amplitude of the dPSP in Control and –VIP INs conditions, respectively.

(B) Subthreshold LEC-driven responses in CA1 PN dendrites before and after VIP IN silencing, average values. Resting membrane potential (p = 0.845), Peak amplitude (p = 0.023), Time of peak (p = 0.317), Half-width (p = 0.687), Area under the curve (p = 0.014). Paired t test, n = 10 dendrites, 12 conditions, 10 slices, 8 mice.

(C) Cumulative frequency distributions of the peak amplitude (left) and maximum dV/dt (right) calculated for every subthreshold LEC-driven dendritic response recorded in Control conditions (black) and the corresponding LEC-driven dendritic response recorded in the –VIP IN condition (red). Data were classified as subthreshold if the maximum dV/dt < 7.5 mV/ms in control conditions. Peak amplitude (p = 0.023), Maximum dV/dt (p = 0.013). Wilcoxon matched-pairs signed-rank test. n = 10 dendrites (∼11 sweeps recorded per dendrite, on average), 12 conditions, 10 slices, 8 mice.

(D) Scatterplot of the peak amplitude (left) and maximum dV/dt (right) of individual LEC-driven dPSPs in Control versus –VIP INs conditions. Diagonal represents the identity line.

(E) Example traces of suprathreshold LEC-driven dSpikes in a CA1 PN distal dendrite in Control (black) and -VIP IN (red) conditions.

(F) Suprathreshold LEC-driven responses in CA1 PN dendrites before and after VIP IN silencing, average values. n = 3 dendrites, 3 slices, 2 mice.

(G) Cumulative frequency distributions of the peak amplitude (left) and maximum dV/dt (right) calculated for every suprathreshold LEC-driven dendritic response recorded in Control conditions (black) and the corresponding LEC-driven dendritic response recorded in the –VIP IN condition (red). Data were classified as suprathreshold if the maximum dV/dt > 7.5 mV/ms in control conditions. Peak amplitude (p = 0.495), Maximum dV/dt (p = 0.008). Wilcoxon matched-pairs signed-rank test. n = 3 dendrites (∼8 sweeps recorded per dendrite, on average), 3 conditions, 3 slices, 2 mice.

(H) Scatterplot of the peak amplitude (left) and maximum dV/dt (right) of individual LEC-driven dSpikes in Control versus –VIP INs conditions. Diagonal represents the identity line.

**Figure S16. Effect of VIP silencing on LEC-driven somatic responses**

(A) Schematic of the mouse brain, illustrating the co-injection of Jaws in CA1 (red) and ChR2 in LEC (green), injected into a VIP-Cre or VIP-Cre/Ai14 mouse.

(B) Schematic illustrating the dual color optogenetics strategy. Same as Figure 5C, but representing whole-cell recordings from CA1 PN somata.

(C) Schematic illustrating the experimental strategy. Same as Figure 5D, but done during whole-cell recordings from CA1 PN somata.

(D) Example traces of the LEC-driven somatic responses in the CA1 PN somata in Control (blue) and – VIP INs (red) conditions. Blue and red dotted lines indicate the peak amplitude of the dPSP in Control and -VIP INs conditions, respectively.

(E) LEC-driven responses in CA1 PN somata before and after VIP IN silencing. From left to right: Resting membrane potential (p = 0.468), Peak amplitude (p = 0.730), Time of peak (p = 0.698), Half-width (p = 0.754), Area under the curve (p = 0.503). Paired t test, n = 7 CA1 PN somata, 7 conditions, 5 slices, 3 mice.

(F) Cumulative frequency distributions of the peak amplitude (left) and maximum dV/dt (right) calculated for every LEC-driven somatic response. Peak amplitude (p = 0.752), Maximum dV/dt (p = 0.295). Wilcoxon matched-pairs signed rank test, n = 7 CA1 PN somata (∼7 sweeps recorded per soma, on average), 7 conditions, 5 slices, 3 mice

(G) Scatterplot of the peak amplitude (left) and maximum dV/dt (right) of individual LEC-driven somatic PSPs in Control versus –VIP INs conditions. Diagonal represents the identity line.

**Figure S17. Jaws expression in CA1**

(A) Representative confocal images demonstrating adequate (left) or poor (right) Jaws expression in VIP INs in hippocampal area CA1. Data collected from brain slices with poor or no Jaws expression were excluded from further analyses in this study.

(B) Graphs illustrating the relationship between Jaws expression and the effect of VIP silencing on LEC-driven dendritic (top) and somatic (bottom) responses. Linear regression lines and R-squared values indicated. The slope of the linear regression line was determined to be significantly non-zero for dendritic data (p = 0.0314) but not statistically different than zero for somatic data (p= 0.6088).

**Figure S18. Effect of VIP silencing on LEC-driven somatic output**

(A) Schematic of model connectivity and activity patterns, calibrated to produce LEC-driven action potentials in CA1 PN somata. As in Figure S13, the schematic also illustrates the deletion of CR^+^ VIP IN and/or CCK^+^ VIP INs, as well as the deletion of CCK INs (black x’s).

(B) Action potential probability of the model CA1 PN, resulting from a combination of somatic depolarization and various frequencies of theta-filtered Poisson-like activity of excitatory LEC inputs (10-100 Hz), in control conditions and upon deletion of various LEC-driven interneurons. **p < 0.005, ****p < 0.0001. Tukey’s multiple comparison test, n = 10 x 1000 trials.

## STAR Methods

- Key Resources Table
- Resource Availability
- Experimental Model and Subject Details

Animals
- Method Details

Viruses
Viral Injections
Slice preparation and electrophysiology
Immunohistochemistry
Confocal imaging
- Quantification and statistical analysis

Data analysis
Statistical analysis
Computational modeling

### Key Resources Table

| Reagent or Resource | Source | Identifier |
| --- | --- | --- |
| **Mouse lines** |  |  |
| CaMKII-Cre | Jackson Laboratory | JAX: 005359; RRID: IMSR_JAX: 005359<br>(Tsien et al., 1996) |
| VIP-IRES-Cre | Jackson Laboratory | JAX: 010908 and 031628; RRID: IMSR_JAX:010908 and IMSR_JAX:031628<br>(Taniguchi et al., 2011) |
| SST-IRES-Cre | Jackson Laboratory | JAX: 013044; RRID: IMSR_JAX:013044<br>(Taniguchi et al., 2011) |
| CCK-IRES-Cre | Jackson Laboratory | JAX: 012706; RRID: IMSR_JAX: 012706<br>(Taniguchi et al., 2011) |
| CR-IRES-Cre | Jackson Laboratory | JAX: 010774; RRID: IMSR_JAX: 010774<br>(Taniguchi et al., 2011) |
| VIP-IRES-Flp | Jackson Laboratory | JAX: 028578; RRID: IMSR_JAX: 028578<br>(He et al., 2016) |
| Dlx-Flp | Obtained from Gord Fishell lab (NYU/Harvard) | (Miyoshi et al., 2010) |
| Ai14(RCL_tdT)-D | Jackson Laboratory | JAX: 007914; RRID: IMSR_JAX: 007914<br>(Madisen et al., 2010) |
| Ai65D(RCFL-tdT)-D | Jackson Laboratory | JAX: 021875; RRID: IMSR_JAX: 021875<br>(Madisen et al., 2015) |
| Ai80D(RCFL-CatCh)-D | Jackson Laboratory | JAX: 025109; RRID: IMSR_JAX: 025109<br>(Daigle et al., 2018) |
| **Viruses** |  |  |
| pAAV-CaMKIIa-hChR2(H143R)-eYFP | Addgene | Catalog Number: 26969-AAV9<br>(Lee et al., 2010) |
| rAAV-CAG-FLEX-rc [Jaws-KGC-GFP-ER2] | Addgene | Catalog Number: 84445-AAV5<br>(Chuong et al., 2014) |
| **Electrophysiology chemicals** |  |  |
| Tetrodotoxin | Hello Bio | Catalog Number: HB1035 |
| 4-aminopyridine | Sigma | Catalog Number: A0152 |
| SR 95531 hydrobromide | Hello Bio | Catalog Number: HB0901 |
| CGP 55845 hydrochloride | Tocris | Catalog Number: 1248 |
| **Immunostaining reagents** |  |  |
| Neurotrace 435/455 Blue Fluorescent Nissl Stain | Fisher Scientific | Catalog Number: N21479 |
| Rabbit anti-GFP (polyclonal) | Invitrogen | Catalog Number: A6455; RRID: AB_221570 |
| Goat anti-rabbit 488 | Invitrogen | Catalog Number: A11008; RRID: AB_143165 |
| Biocytin | Invitrogen | Catalog Number: B1592 |
| Alexa Fluor 647-conjugated Streptavidin | Thermo Fisher Scientific | Catalog Number: S32357 |
| *CUBIC tissue clearing reagents* |  |  |
| Triton X-100 | Sigma-Aldrich | Catalog Number: T8787 |
| Urea | Sigma-Aldrich | Catalog Number: 33247 |
| Triethanolamine | Sigma-Aldrich | Catalog Number: T58300 |
| N,N,N',N'-Tetrakis(2-Hydroxypropyl)ethylenediamine (Quadrol) | Sigma-Aldrich | Catalog Number: 122262 |
| **Software** |  |  |
| pClamp 9 | Molecular Devices | N/A |
| AxoGraph (v1.7.0) | Axograph | N/A |
| MATLAB R2019b | Mathworks | N/A |
| GraphPad Prism 9 | GraphPad | N/A |
| NEURON (v7.8.2) | Carnevale and Hines, 2006 | N/A |
| Python (v3.8.5) | https://www.python.org/ | N/A |
| ImageJ (FIJI) | NIH | N/A |
| Illustrator | Adobe | N/A |

### Experimental Model and Subject Details

#### Animals

All experiments were conducted in accordance with the National Institutes of Health guidelines and with the approval of the *New York University Grossman School of Medicine and NYU Langone Health Institutional Animal Care and Use Committee (IACUC).* Mouse lines were obtained from the Jackson Laboratory (JAX) and crossed to reporter lines in house. Slice electrophysiology experiments were done on acute brain slices from CaMKII-Cre (Tsien et al., 1996) or interneuron-specific mouse lines. To fluorescently label VIP- or SST-expressing interneurons, homozygous VIP-Cre and SST-Cre mice (Taniguchi et al., 2011) were crossed with homozygous Cre-dependent Ai14-tdTomato reporter mice (Madisen et al., 2010), respectively. To fluorescently label CCK-expressing interneurons, CCK-Cre drive mice (Taniguchi et al., 2011) were first crossed with the Cre-/Flp-dependent Ai65-tdTomato reporter strain (Madisen et al., 2015), and the resulting progeny were crossed with the Dlx-Flp driver mice (gift from Gord Fishell, NYU/Harvard, (Miyoshi et al., 2010)), producing a triple transgenic CCK-Cre/Dlx-Flp/Ai65 mouse line with tdTomato expression restricted to GABAergic CCK-expressing interneurons (Basu et al., 2013). To fluorescently label VIP IN subpopulations, double homozygous VIP-Flp/CR-Cre or VIP-Flp/CCK-Cre mice were crossed with homozygous Cre/Flp-dependent Ai65-tdTomato reporter mice. To express excitatory opsin CatCh (Kleinlogel et al., 2011) in VIP IN subpopulations, double homozygous VIP-Flp/CR-Cre or VIP-Flp/CCK-Cre mice were crossed with homozygous Cre- /Flp-dependent Ai80-CatCH-expressing reporter mice (Daigle et al., 2018). Litter mates were co-housed up to 5 mice per cage and were provided with food and water *ad libitum*. Male and female mice were used, ages 7-12 weeks (PN somatic data: ∼40% males/60% females; PN dendritic date: ∼60% males/40% females). No significant differences were found between data collected from male versus female mice (data not shown).

### Method Details

#### Viruses

Commercially-generated recombinant adeno-associated viruses (rAAVs) were injected into the lateral entorhinal cortex (LEC) or hippocampal area CA1 (dorsal and intermediate) region under stereotactic control. Viruses were stored at -80°C in 2-4 µL aliquots and were thawed prior to use. Co-injected viruses were combined <16 hours before viral injections. For optogenetic activation experiments, AAV2.9-CaMKIIa-hChR2(H143R)-eYFP (Lee et al., 2010) was injected into LEC. For targeted interneuron recordings, the virus was injected into LEC of interneuron-specific transgenic mice. Slice electrophysiology experiments were conducted 7-10 days after viral injection. For optogenetic silencing experiments, AAV2.5-CAG-flex-Jaws-GFP (Chuong et al., 2014) was injected into hippocampal area CA1 of VIP-Cre or VIP-Cre/Ai14 mice, targeting multiple CA1 layers. The Jaws virus incubated for 15-17 days before slice electrophysiology experiments. Meanwhile, the mice were additionally injected into the ipsilateral LEC with AAV2.9-CaMKIIa-hChR2(H143R)-eYFP, as described above, and slice electrophysiology experiments were conducted 7-10 days after injecting the ChR2 virus.

#### Viral injections

Viral injection pipettes were prepared using thin glass pipette (Drummond Scientific), pulled by a micropipette puller (Sutter P-1000, Sutter Instrument Company) and fire polished using a microforge to have a long taper and 7-10 μm tip diameter. Pipettes were back-filled with mineral oil, then front-filled with the virus using a Nanoject II injector (Drummond Scientific). Adult mice (6-11 weeks old) were injected using stereotactic surgery and sterile technique. The animal was deeply anesthetized with inhaled isoflurane (5% for induction, 1.5-3% for maintenance during surgery, Matrx VIP 3000 calibrated vaporizer), and was given an IP injection of 0.05-0.1 mg/kg Buprenorphine before surgery. After head-fixation of the animal in a stereotactic frame (Stoelting), the surgical area was prepared with aseptic technique. The hair on the mouse’s head was shaved with a razor, and the exposed skin was locally disinfected with ethanol and betadine. Before incision, Ropivacaine (1-3 drops) was applied to the skin as a local analgesic. An incision was made to expose the skull, and hydrogen peroxide (0.1%) was applied to the surface of the skull to clean it. The level of the skull was adjusted such that Bregma and Lambda were aligned in the z-axis.

A unilateral craniotomy was made by thinning the skull with a dental drill. After gaining access to the surface of the brain, the injection pipette was lowered into either the lateral entorhinal cortex (LEC) or into the dorsal and intermediate portions of hippocampal area CA1. LEC: A/P - 3.2 ± 0.2, M/L 4.5 ± 0.2, D/V -2.5 ± 0.2 mm from Bregma (6 sites); 46 nL injected per site, left hemisphere, specifically targeting superficial layers (L2/3). Dorsal and intermediate CA1, targeting local VIP INs in various CA1 strata: A/P -2.3 ± 0.3, M/L 1.5 ± 0.2, D/V -1.3 ± 0.2 mm from Bregma and A/P -3.2 ± 0.2, M/L 3.7, D/V -2.5 ± 0.2 mm from Bregma (11 sites total); 46 nL injected per site. The pipette was lowered into the brain, and the virus was injected into each coordinate (2 x 23 nL for a total of 46 nL injected per site). After the final coordinate in the injection site, the pipette was left in place for 10-15 minutes to allow for adequate dispersion of the virus. After slowly withdrawing the injection pipette from the brain, the incision was sutured (Henry Schein) and antibiotic ointment (Neosporin) was applied. Animals were injected with 0.5-1.0 mL saline solution to rehydrate and were allowed to recover for 1-3 weeks to allow for adequate viral expression.

#### Slice preparation and electrophysiology

##### Slice preparation

Adult mice were deeply anesthetized with 5% isoflurane and transcardially perfused with ice-cold sucrose-enriched dissection artificial cerebrospinal fluid (dACSF). After the dissected brain was cut down the midline, the two hemispheres were tilted ventro-medially at a 10° angle, and a vibratome (Leica VT1200S) was used to cut 400 μm-thick horizontal brain slices. The slices were incubated in a 1:1 mixture of sucrose-enriched dACSF and standard ACSF solution for 15 minutes at 34° C and 45 minutes at room temperature before being transferred to the recording chamber containing standard ACSF.

##### Solutions

Sucrose-enriched dissection ACSF (dACSF) consisted of (in mM): Sucrose 195.0, NaCl 10.0, Glucose 10.0, NaHCO_3_ 25.0, KCl 2.5, NaH_2_PO_4_ 1.25, Sodium pyruvate 2.0, CaCl_2_ 0.5, MgCl_2_ 7.0, saturated with 95% O_2_ and 5% CO_2_. Standard artificial cerebrospinal fluid (ACSF) consisted of (in mM): NaCl 125.0, Glucose 22.5, NaHCO_3_ 25.0, KCl 2.5, NaH_2_PO_4_ 1.25, Sodium pyruvate 3.0, Ascorbic Acid 1.0, CaCl_2_ 2.0, MgCl_2_ 1.0, saturated with 95% O_2_ and 5% CO_2_. The intracellular solution contained the following (in mM): KMeSO_4_ 135 (for current clamp recordings) or CsMeSO_4_ 135 (for voltage clamp recordings), KCl 5, EGTA 0.1, HEPES 10, NaCl 2, Mg-ATP 5, Na_2_-GTP 0.4, Na_2_Phosphocreatine 10, and Neurobiotin or Biocytin (0.2%). The pH of the intracellular solution was adjusted to 7.2 with KOH as needed. Osmolarity ranged from 270-290, and was adjusted with sterifiltered MilliQ water. In a subset of experiments, the following pharmacological agents were used via bath application in the ACSF: TTX (1 μM), 4-AP (100 μM), SR95531 (2 μM), and CGP55845 (1 μM).

##### Electrophysiology setu

Slices were visualized with a LNscope microscope from Luigs & Neumann, equipped with Infracontrast Dodt Gradient Contrast optics (Luigs & Neumann), 5x air and 60x/0.8 nA water immersion objective (Olympus), 1-4x zoom optics (Luigs & Neumann), and a Hamamatsu ORCA-R2 CCD camera using ImageJ Micromanager imaging acquisition software. A MultiClamp 700B amplifier (Axon Instruments), Digidata 9 Digitizer (Axon Instruments), and two constant current stimulators (Digitimer Ltd.) using pClamp 9 software (Molecular Devices) were used for data acquisition. Optogenetic stimulation and fluorescence-guided targeted patch-clamp recordings were performed using a Thorlabs LED system (405/470/565/635 nm wavelengths). Patch-clamp recordings were performed using junior micromanipulators (Luigs & Neumann) on movable motorized shifting tables (Luigs & Neumann). Recordings were performed at 34°C in standard ACSF, maintained by an inline heater (Warner Instruments) and saturated with 95% O_2_ and 5% CO_2_. Fire-polished borosilicate glass pipettes (Sutter) were used with tip resistances of 3.0-5.5 MΩ for somatic recordings (up to 6.5 MΩ for VIP IN somata) and 13-17 MΩ for dendritic recordings, pulled with a micropipette puller (Sutter P-1000). Current clamp recordings were obtained with access resistances up to 20 MΩ for somata and up to 40 MΩ for dendrites, compensated in bridge balance mode. The series resistance was not compensated for voltage-clamp recordings.

##### Electrophysiology recordings

To confirm adequate expression and function of ChR2 in infected LEC neurons, slices containing LEC were photostimulated over LEC using two protocols. First, a brief train of 470 nm light was given in cell-attached mode (10 Hz, 2 ms, 100% LED power), to test for time-locked action potentials. Next, in whole-cell configuration, a 500 ms-long light pulse was delivered, to check for a large, excitatory photocurrent. These protocols were also used with local CA1 photostimulation at the beginning of every recording to confirm the lack of off-target viral expression in area CA1. If CA1 photostimulation produced a large photocurrent in the recorded CA1 neuron, then all data from that animal were discarded.

CA1 neurons were targeted for recording throughout the proximal-distal of area CA1 with a focus on the middle and distal (close to subiculum) thirds (**Figures S1E** and **S8A**), where glutamatergic LEC inputs are most dense. Both deep and superficial CA1 pyramidal neurons were targeted for recordings.

To measure the intrinsic membrane and firing properties of CA1 neurons, the first current was injected into the neuron to maintain a membrane potential of -70 mV for the following protocols. Depolarizing or hyperpolarizing current steps (1 second) were then injected into the neuron to measure the firing and sag properties, respectively. The firing was measured as the number of spikes during the 1 second depolarizing current step. The sag ratio was calculated as ((V_Peak_–V_SteadyState_)/V_Peak_)*100%, where V_Peak_ and V_SteadyState_ represent the change in membrane potential from baseline (-70 mV) to the peak sag and steady state response, respectively.

To measure light-evoked post-synaptic responses, CA1 neurons were kept at their resting membrane potential (RMP). Light-evoked responses were recorded every 15 seconds. Unless otherwise stated, a 2 ms light pulse (470 nm) was delivered at 100% maximum photostimulation strength (56 mW, 50 µm diameter beam spot). To measure the input-output transformation of light-evoked responses, the stimulation strength was incrementally adjusted, keeping pulse duration constant. To measure the short-term plasticity dynamics of the LEC inputs, 5 light pulses (2 ms each) were given at 1-10 Hz at low photostimulation strength (2-3% maximum). To measure the summation of LEC-driven post-synaptic responses, 10 light pulses (2 ms each) were given at 2-10 Hz at maximum stimulation strength. For experiments involving pharmacology, light-evoked responses were recorded in control conditions (at least 5 minutes), during drug infusion (at least 10 minutes, while monitoring cell health), and in the experimental condition (at least 5 minutes). The amount of current injected was continuously monitored and adjusted to keep the neuron’s membrane potential close to the resting membrane potential in control conditions.

For the optogenetic silencing experiments, light-evoked post-synaptic responses were recorded every 15 seconds, alternating between control and experimental conditions. In the control condition, 470 nm light was used to photostimulate glutamatergic LEC axons only. In the experimental condition, 470 nm and 625 nm light were used simultaneously to optogenetically activate glutamatergic LEC axons and silence local CA1 VIP INs, respectively. The strength and duration of the 470 nm light pulse were adjusted (30-80% maximum photostimulation strength, 0.2 ms) to avoid potential crosstalk with Jaws expressed in the VIP INs in area CA1. To ensure maximum silencing of VIP INs during photostimulation of LEC inputs, the 625 nm light pulse (100% maximum photostimulation, 100 ms) began 50-60 ms before the 470 nm light pulse. The 625 nm light pulses ended with a down ramp (100 ms) to decrease the probability of rebound firing in the VIP INs. When LEC axons were photostimulated with an 8 Hz train of light pulses (0.2 ms each, 40-60% maximum photostimulation strength), VIP INs were silenced with 625 nm light for 1 second. Data from slices with insufficient Jaws expression (**Figure S18**) were discarded.

#### Immunohistochemistry

Animals were deeply anesthetized with 5% isoflurane for 5 minutes, followed by an injection of a mixture of ketamine (150 mg/kg) and xylazine (10 mg/kg). Animals were then perfused with 1X PBS, followed by 4% paraformaldehyde in PBS. The brain was removed and fixed overnight in 4% paraformaldehyde in 4°C and then sectioned into 100 μm thick slices using a microtome (Leica VT1000S) after washing. Alternatively, after slice electrophysiology experiments, brain slices were drop-fixed in 4% paraformaldehyde in PBS and left overnight at 4°C. The slices were washed once with 0.3 M PBS-Glycine for 15-20 minutes and three times in PBS. Slices were left in PBS at 4°C until immunostaining.

Slices that underwent CUBIC tissue clearing (Susaki et al., 2015) were thoroughly washed with 0.1M PB and then moved to CUBIC#1 tissue clearing solution for two days. Next, slices were thoroughly washed in 0.1M PB before undergoing the standard immunostaining protocol. Slices were permeabilized in 0.5% PBS-Triton X (PBST), blocked in 3% Normal Goat Serum (2-4 hours), incubated with primary antibodies (overnight, 4°C), washed, and incubated with secondary antibodies (1-2 days, 4°C). The antibodies were diluted in blocking solution (1X PBS, 0.5% Triton, 3% NGS). After the final antibody incubation step, the slices were thoroughly washed in 0.1M PB and moved to CUBIC#2 tissue clearing solution for 1 hour, after which the slices were mounted on glass slides in CUBIC#2 solution. Slices that did not undergo tissue clearing immediately started with the standard immunostaining protocol, beginning with the permeabilization in 0.5% PBST and ending with the secondary antibody incubation step. These slices were then thoroughly washed in 1X PBS and mounted in Vectashield Hard Set Mounting Medium with DAPI (Vector Laboratories).

Slices containing ChR2-eYFP LEC axons and/or Jaws-GFP-labeled VIP INs were stained using a rabbit polyclonal anti-GFP primary antibody (1:1000; Invitrogen) and a goat anti-rabbit Alexa Fluor 488 dye-conjugated IgG antibody secondary antibody (1:1000; Invitrogen). Alexa Fluor 593 or 647 streptavidin (1:500; Thermo Fischer Scientific) was added with the secondary antibody to counterstain the cells filled with biocytin during intracellular recordings. For slices that underwent the combined tissue clearing and immunostaining steps, Neurotrace 435/455 was added with the secondary antibody to label somata in the brain slice (1:200; Thermo Fischer Scientific). Data from animals with inadequate, mistargeted, or retrograde viral infection (of ChR2 and/or Jaws) were discarded.

#### Confocal Imaging

Slices were imaged using an upright Zeiss 520 Meta Confocal Microscope, using (magnification/NA) 10x/0.3 air, 20x/0.8 air, and/or 40x/1.30 oil immersion objectives (Zeiss) and 405, 488, 594, and 647 nm lasers for excitation. 512 x 512 pixel images were acquired every 5-10 microns throughout the depth of the tissue. Tiled images were stitched using Zen Microscopy Software (Zeiss), and the tiled z-stacks were subsequently processed using Fiji (ImageJ).

### Quantification and statistical analysis

#### Data analysis

*In vitro* electrophysiology data were analyzed using Axograph X (Version 1.7.0). After measuring the baseline membrane potential, all traces were zeroed at the start of photostimulation (x = 0) and baseline membrane potential (y = 0). Subthreshold light-evoked responses were averaged in Axograph before measuring the amplitude and kinetics of the average response. Suprathreshold light-evoked responses were measured first and then the values averaged.

For all light-evoked post-synaptic responses, peak amplitude was measured as the maximum depolarization from the baseline. The time of peak was measured as the duration from the start of photostimulation (x = 0) to peak amplitude. The rise time was measured as the duration from 10% to 90% of peak amplitude. The half-width was measured as the duration between the 50% peak amplitude during the rise and decay of depolarization. The IPSP amplitude was measured as the maximum negative amplitude of the hyperpolarization component of the post-synaptic response from the baseline.

For dendritic recordings, derivative voltage traces and phase plots were generated from dendritic post-synaptic response traces using MATLAB R2019b (MathWorks). The peak derivative was measured per trace and averaged per dendrite to produce the maximum dV/dt value, used to categorize the dendritic response. Within our experimental dataset, light-evoked dendritic responses with dV/dt < 7.5 mV/ms were categorized as dendritic post-synaptic potentials (dPSPs), whereas those with dV/dt > 7.5 mV/ms were categorized as dendritic spikes (dSpikes). Dendritic recording distance was measured in ImageJ using a 5x image taken after completing the dendritic recording.

To measure the paired pulse ratio (PPR), the peak amplitude of each pulse was measured from the baseline membrane potential immediately preceding that peak (P1 through P5), then divided by the amplitude of the first peak (P1). PPR > 1 indicates short-term facilitation of inputs. PPR < 1 indicates short-term depression of inputs. To measure the summation, or amplitude ratio, the peak amplitude of each pulse (P1 through P10) was measured from the baseline membrane potential immediately preceding P1, then divided by the amplitude of the first peak (P1). Photostimulation protocols were run three times, and the resulting PPR or amplitude ratio values were averaged per cell. During experiments that combined trains of photostimulation with somatic depolarization, the Expected number of spikes was calculated as the linear sum of (the number of action potentials recorded in the first second of the 8 Hz photostimulation condition) plus (the number of action potentials recorded during 1 second of depolarization with 100 pA somatic current injection).

During pharmacological inhibition blockade experiments, the Schaffer collaterals (CA3-CA1 connection) were first manually cut with a scalpel before the start of recording to prevent runaway epileptiform activity during the experiment. Data that suggested that the recorded cell was experiencing the effects of runaway epileptiform activity were discarded. Inferred IPSP traces were generated post hoc in Axograph as the post-synaptic potential recorded in control conditions (PSP) minus the post-synaptic potential recorded after inhibition blockade (EPSP).

During targeted recordings from CA1 interneurons, the location of the patched soma was noted as: SLM, SR/SLM border, SR, SP, or SO (**Figure S8**). Interneurons were counted as responding to LEC inputs if they exhibited a reliable light-evoked post-synaptic response and were counted as driven to spike by LEC inputs if they responded with a light-evoked action potential.

During the voltage clamp experiments investigating the downstream targets of VIP IN subpopulations, neurons were counted as targeted by a VIP IN subpopulation if they exhibited a reliable light-evoked inhibitory post-synaptic current (IPSC). Because IPSC amplitudes were often less than 20 pA, connectivity was determined while blinded to the identity of the recorded neuron. For optogenetic silencing experiments, the efficacy of Jaws was first confirmed by measuring the hyperpolarization amplitude and ability to reduce firing in VIP INs in response to 625 nm light at 100% maximum photostimulation strength. Next, data from the dual color optogenetic stimulation were analyzed by comparing pairs of control and experimental post-synaptic responses. Both individual and average traces were analyzed. Individual dendritic responses were categorized as subthreshold or suprathreshold, based on the maximum dV/dt value calculated for the post-synaptic response in the control condition. Cumulative frequency distribution plots were generated from individual traces, whereas scatterplots were generated from the amplitude or kinetics values of averaged traces per cell.

### Statistical analysis

No statistical methods were used to predetermine sample size. Dataset normality was determined using the Shapiro-Wilk test. Data were analyzed using appropriate parametric or non-parametric tests, as stated in the text and figure legends. When necessary, analyses were corrected for multiple comparisons using post hoc tests, as indicated. Data are shown as mean ± SEM, unless otherwise stated. Significance level was set at p < 0.05. Prism (GraphPad) was used for plotting data and statistical analyses. Figures were generated using Adobe Illustrator.

### Computational modeling

The neuronal models were developed based on prior computational models of the hippocampus (Shuman et al., 2020; Turi et al., 2019). Specifically, we implemented the simplified morphology and ionic mechanisms for the individual neuronal models (i.e., pyramidal cells, OLM INs, non-basket cell CCK INs in the SR/SLM region, CCK^+^ VIP INs, and CR^+^ VIP INs) based on (Cutsuridis and Poirazi, 2015). Individual cells were re-validated to match experimental data (experimental data from **Figures 1**-**5, S2, 4-5, 8-9;** validation in **Figures S10-12**), specifically the active and passive membrane properties such as input resistance, resting potential, and rheobase current. LEC provided input to the network as a single spike or as a Poisson distributed spike train (see sections below). For all simulations, we used the NEURON (v7.8.2) simulation environment (Carnevale and Hines, 2006) while analysis was performed in python (v3.8.5) using custom-based software. The simulations were performed on a server with Intel(R) Xeon(R) Silver 4210R CPU at 2.40GHz and 126 GB of RAM under CentOS (v 7.8) operating system. The code is available via the ModelDB webpage, under accession number 267221 (http://modeldb.yale.edu/267221).

#### Model neurons

Individual neuron types were modeled as simplified biophysical neurons with various active dendrites using the Hodgkin-Huxley formalism. Specifically, the pyramidal cells incorporated 27 apical and basal dendrites to capture their complex morphology, while most interneurons were simulated with a simplistic morphology, consisting of apical and basal dendrites in an ‘x’ shape. The OLM IN consisted of a soma and two sister dendrites (Cutsuridis et al., 2010). The morphological features of all neuronal types are listed in **Tables 1** and **2** for interneurons and pyramidal cells, respectively. The SR/SLM CCK IN model cell was generated *de novo* as described below.

**Table 1.** Morphological properties of the SR/SLM CCK IN, OLM IN, CCK^+^ VIP IN, and CR^+^ VIP IN. d and L denote the diameter and length of the compartments, respectively.

| Name of compartment | SR/SLM CCK IN |  | OLM IN |  | CCK<sup>+</sup> VIP IN |  | CR<sup>+</sup> VIP IN |  |
| --- | --- | --- | --- | --- | --- | --- | --- | --- |
|  | d x L (μm<sup>2</sup>) / # |  | d x L (μm<sup>2</sup>) / # |  | d x L (μm<sup>2</sup>) / # |  | d x L (μm<sup>2</sup>) / # |  |
|  | compartments |  | compartments |  | compartments |  | compartments |  |
| soma | 10 x 20 | 1 | 10 x 20 | 1 | 10 x 20 | 1 | 10 x 20 | 1 |
| radProx | 4.0 x 100 | 2 | - | - | 4.0 x 100 | 2 | 4.0 x 100 | 2 |
| radMed | 3.0 x 100 | 2 | - | - | 3.0 x 100 | 2 | 3.0 x 100 | 2 |
| radDist | 2.0 x 200 | 2 | - | - | 2.0 x 200 | 2 | 2.0 x 200 | 2 |
| lmM | 1.5 x 100 | 2 | - | - | 1.5 x 100 | 2 | 1.5 x 100 | 2 |
| Lmt | 1.0 x 100 | 2 | - | - | 1.0 x 100 | 2 | 1.0 x 100 | 2 |
| oriProx | 2.0 x 100 | 2 | - | - | 2.0 x 100 | 2 | 2.0 x 100 | 2 |
| oriMed | 1.5 x 100 | 2 | - | - | 1.5 x 100 | 2 | 1.5 x 100 | 2 |
| oriDist | 1.0 x 100 | 2 | - | - | 1.0 x 100 | 2 | 1.0 x 100 | 2 |
| dend | - | - | 3.0x250 | 2 | - | - | - | - |
| axon | - | - | 1.5x150 | 1 | - | - | - | - |
| **Total** |  | **17** |  | **4** |  | **17** |  | **17** |

#### Random distributions

To account for the variability observed in experiments, all synaptic numbers were drawn from probability distributions. We used the Poisson (Eq. 1), the Exponential (Eq. 2), and the Gaussian (Eq. 3) distributions. The synaptic location on the postsynaptic neuron was chosen based on a uniform distribution, i.e., all sites were equally probable to contain a synapse.

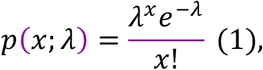

where *x* denotes the number of events, and *λ* denotes the mean of the distribution. ⋅ ! is the factorial of a number.

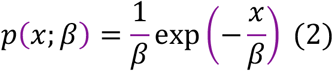

where *β* is the scale parameter which represents the mean of the distribution.

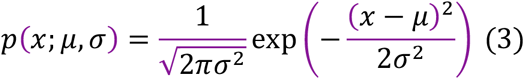

where *μ* denotes the mean and *σ* the standard deviation.

We returned the rounded to the closer integer value for all distributions, as the number of synapses cannot be a float number.

#### Pyramidal neuron (CA1 PN) model cell

The CA1 PN model cell was adapted from (Turi et al., 2019) and modified to increase its complexity. Specifically, we increased the number of compartments representing the apical trunk and adjusted all compartments’ lengths (**Table 2**). The diameters of all compartments were updated to follow the d3/2 rule. The model consists of five apical dendritic compartments simulating the apical trunk (radProx, radMed, radDist_i, *ϵ* ∈ {1,2,3}) and six dendritic compartments simulating the basal tree, each containing a Ca^2+^ pump and buffering mechanism, Ca^2+^ activated slow AHP and medium AHP potassium (K^+^) currents, an HVA L-type Ca^2+^ current, an HVA R-type Ca^2+^ current, an LVA T-type Ca^2+^ current, an h current, a fast Na^+^ and a delayed rectifier K^+^ current, a slowly inactivating K^+^ M-type current and a fast inactivating K^+^ A-type current (Poirazi et al., 2003a, b). The proximal and distal apical dendrites consisted of eight compartments, respectively, each containing a fast Na^+^, a delayed rectifier K^+^ and a fast-inactivating K^+^ A-type current. The PN model cell was topologically oriented: its soma was located in the simulated SP layer, its basal dendrites in the SO layer, and its proximal and distal apical dendrites in the SR and SLM layers, respectively. The type and distribution of ionic mechanisms in the PN model are described below, while each channel’s maximum conductance is given in **Tables 3** and **4**.

**Table 2.** Morphological properties of the pyramidal model cell. Prefix ‘radT’ in compartments correspond to the apical trunk, ‘rad’ to oblique dendrites, ‘lm’ to apical tuft, and prefix ‘ori’ to basal dendrites, respectively. d and L denote the diameter and length of the compartment.

| Name of compartment | CA1 PN |  |
| --- | --- | --- |
| | d x L ($\mu\text{m}^2$) / # compartments | |
| soma | 10.0 x 20 | 1 |
| axon | 1.0 x 150 | 1 |
| radTProx | 3.0 x 80 | 2 |
| radTMed | 2.5 x 70 | 2 |
| radTdist1 | 2.0 x 50 | 1 |
| radTdist2 | 1.75 x 50 | 1 |
| radTdist3 | 1.5 x 50 | 1 |
| lm_thick | 0.95 x 50 | 2 |
| lm_medium | 0.9 x 50 | 2 |
| lm_thin | 0.6 x 75 | 4 |
| rad_thick | 1.9 x 50 | 2 |
| rad_medium | 1.5 x 50 | 2 |
| rad_thin | 0.95 x 75 | 4 |
| oriprox | 1.6 x 75 | 2 |
| oridist | 1.1 x 75 | 4 |
| **Total** |  | **31** |

**Table 3.** Passive parameters and active ionic conductances for somatic, axonal, and apical dendritic trunk compartments of the CA1 PN model cell.

| Mechanism | Compartment |  |  |  |  |  |  |
| --- | --- | --- | --- | --- | --- | --- | --- |
|  | Soma | Axon | radTprox | radTmed | radTdist |  |  |
|  |  |  |  |  | 1 | 2 | 3 |
| Leak conductance [S/cm<sup>2</sup>] | 5e-8 | 5e-8 | 5.26e-8 | 6.67e-8 |  | 2.85e-7 |  |
| Na<sup>+</sup> conductance [S/cm<sup>2</sup>] | 3.5e-2 | 1.5e0 | 1.4e-2 | 1.4e-2 |  | 1.4e-2 |  |
| Delayed rectifier K<sup>+</sup> conductance [S/cm<sup>2</sup>] | 1.4e-3 | 1e-1 | 1.74e-3 | 1.73e-3 |  | 1.73e-3 |  |
| Proximal A-type K<sup>+</sup> conductance [S/cm<sup>2</sup>] | 7.5e-3 | x | 1.8e-3 | x |  | x |  |
| Distal A-type K<sup>+</sup> conductance [S/cm<sup>2</sup>] | x | x | x | 2.78e-2 | 3.75e-2 | 4.73e-2 | 4.86e-2 |
| M-type K<sup>+</sup> conductance [S/cm<sup>2</sup>] | 1.66e-2 | 3e-6 | 6e-4 | 6e-4 |  | 6e-4 |  |
| I<sub>h</sub> conductance [S/cm<sup>2</sup>] | 1.5e-5 | x | 1.8e-5 | 6e-5 |  | 9e-5 |  |
| L-type Ca<sup>2+</sup> conductance [S/cm<sup>2</sup>] | 7e-3 | x | 3.16e-6 | 6.32e-6 | 1.27e-5 | 1.9e-5 | 2.53e-5 |
| R-type Ca<sup>2+</sup> conductance [S/cm<sup>2</sup>] | 3e-3 | x | 3e-4 | 3e-4 |  | 3e-4 |  |
| T-type Ca<sup>2+</sup> conductance [S/cm<sup>2</sup>] | 5e-5 | x | 4e-4 | 4e-4 |  | 4e-4 |  |
| Ca<sup>2+</sup>-dependent mAHP K<sup>+</sup> conductance [S/cm<sup>2</sup>] | 9.075e-2 | x | 6.6e-3 | 6.6e-3 |  | 4.12e-3 |  |
| Ca<sup>2+</sup>-dependent sAHP K<sup>+</sup> conductance [S/cm<sup>2</sup>] | 5e-4 | x | 5e-4 | 5e-4 |  | 5e-5 |  |
| Membrane capacitance c<sub>m</sub> [μF/cm<sup>2</sup>] | 1 | 1 | 2.2 | 2.2 |  | 2.2 |  |
| Membrane resistance R<sub>m</sub> [Ohm cm<sup>2</sup>] | 200,000 | 200,000 | 190,000 | 150,000 |  | 35,000 |  |
| Axial resistance $R_a$<br>[Ohm cm] | 300 | 300 | 285 | 270 | 255 | 225 | 216 |
| $V_{half,h}$ [mV] | -90 | x | -90 | -90 | | -90 | |
| $E_L$ [mV] | -70 | -70 | -70 | -70 | | -70 | |
| $E_{Na}$ [mV] | 50 | 50 | 50 | 50 | | 50 | |
| $E_K$ [mV] | -80 | -80 | -80 | -80 | | -80 | |
| $E_h$ [mV] | -10 | x | -10 | -10 | | -10 | |
| $E_{Ca}$ [mV] | 140 | x | 140 | 140 | | 140 | |

#### SR/SLM CCK IN model cell

The non-basket cell SR/SLM CCK IN had 17 compartments, containing a leak conductance, a sodium current, a fast-delayed rectifier K^+^ current, an A-type K^+^ current, L- and N-type Ca^2+^ currents, a calcium-dependent K^+^ current, and calcium- and voltage-dependent K^+^ current (**Table 5, Figure S12**). The SR/SLM CCK IN received excitatory connections from LEC to their distal SLM dendrites and from the CR^+^ VIP IN to their soma. To reproduce the LEC-induced activity on SR/SLM CCK INs (see **Figure 4G-I, S12B-C**), we drew the number of excitatory synapses from an exponential distribution with mean equals 10. At approximately 17% of the trials (n = 10 x 200 trials), the SR/SLM CCK IN receives no LEC connections (**Figure 4H**). The CR^+^ VIP IN also targeted the SR/SLM CCK IN, with the synaptic number drawn from an exponential distribution with mean equals 60. Using this value, the SR/SLM CCK IN was silenced when the number of inhibitory synapses was chosen to be high enough, but in other cases, remained intact.

**Table 4.**
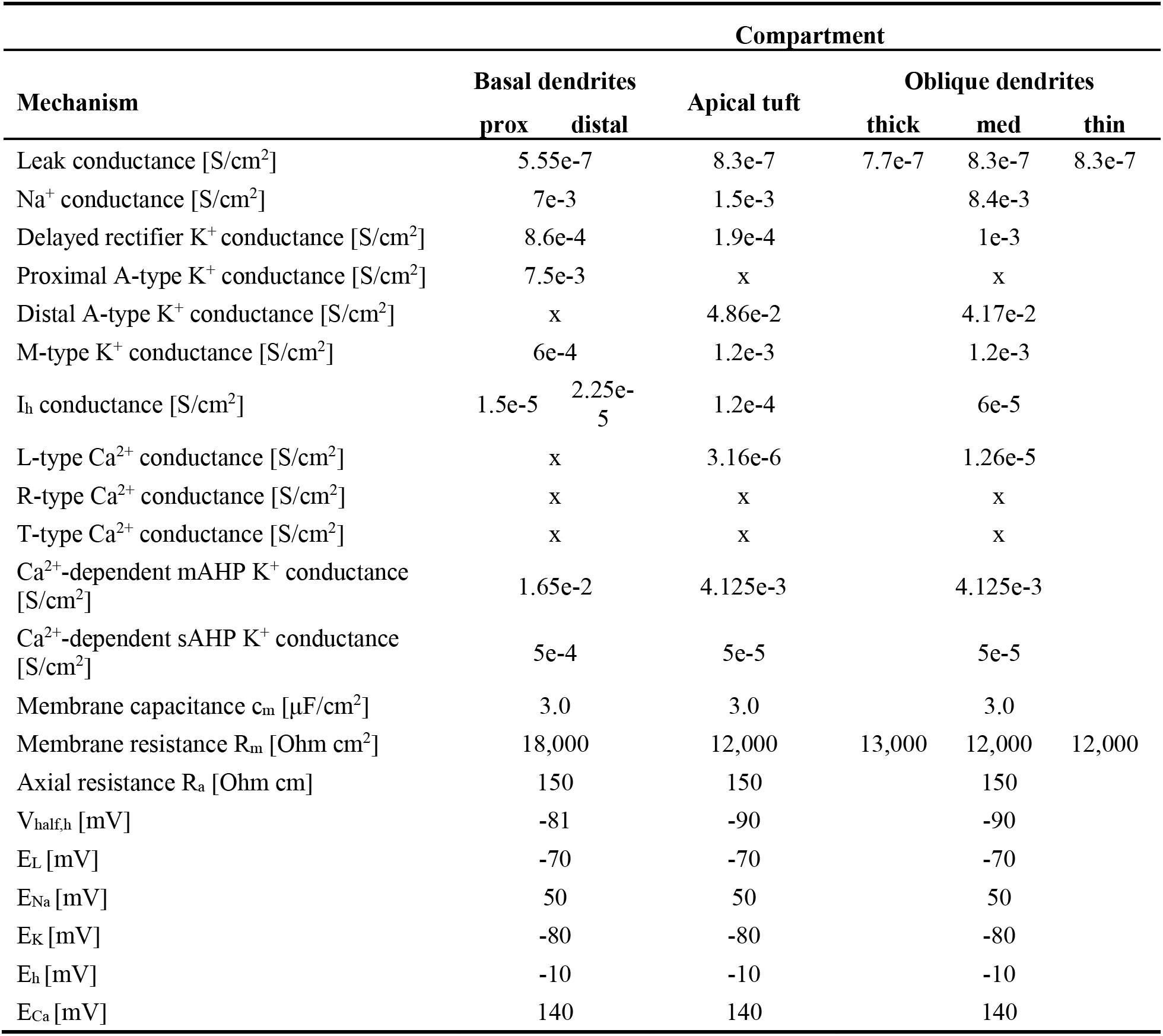
Passive parameters and active ionic conductances for basal (SO), apical tuft (SLM), and oblique (SR) dendrites of the CA1 PN model cell.

| Mechanism | Compartment |  |  |  |  |  |
| --- | --- | --- | --- | --- | --- | --- |
|  | Basal dendrites |  | Apical tuft | Oblique dendrites |  |  |
|  | prox | distal |  | thick | med | thin |
| Leak conductance [S/cm<sup>2</sup>] |  | 5.55e-7 | 8.3e-7 | 7.7e-7 | 8.3e-7 | 8.3e-7 |
| Na<sup>+</sup> conductance [S/cm<sup>2</sup>] |  | 7e-3 | 1.5e-3 |  | 8.4e-3 |  |
| Delayed rectifier K<sup>+</sup> conductance [S/cm<sup>2</sup>] |  | 8.6e-4 | 1.9e-4 |  | 1e-3 |  |
| Proximal A-type K<sup>+</sup> conductance [S/cm<sup>2</sup>] |  | 7.5e-3 | x |  | x |  |
| Distal A-type K<sup>+</sup> conductance [S/cm<sup>2</sup>] |  | x | 4.86e-2 |  | 4.17e-2 |  |
| M-type K<sup>+</sup> conductance [S/cm<sup>2</sup>] |  | 6e-4 | 1.2e-3 |  | 1.2e-3 |  |
| I<sub>h</sub> conductance [S/cm<sup>2</sup>] | 1.5e-5 | 2.25e-5 | 1.2e-4 |  | 6e-5 |  |
| L-type Ca<sup>2+</sup> conductance [S/cm<sup>2</sup>] |  | x | 3.16e-6 |  | 1.26e-5 |  |
| R-type Ca<sup>2+</sup> conductance [S/cm<sup>2</sup>] |  | x | x |  | x |  |
| T-type Ca<sup>2+</sup> conductance [S/cm<sup>2</sup>] |  | x | x |  | x |  |
| Ca<sup>2+</sup>-dependent mAHP K<sup>+</sup> conductance [S/cm<sup>2</sup>] |  | 1.65e-2 | 4.125e-3 |  | 4.125e-3 |  |
| Ca<sup>2+</sup>-dependent sAHP K<sup>+</sup> conductance [S/cm<sup>2</sup>] |  | 5e-4 | 5e-5 |  | 5e-5 |  |
| Membrane capacitance $c_m$ [μF/cm<sup>2</sup>] | | 3.0 | 3.0 | | 3.0 | |
| Membrane resistance $R_m$ [Ohm cm<sup>2</sup>] | | 18,000 | 12,000 | 13,000 | 12,000 | 12,000 |
| Axial resistance $R_a$ [Ohm cm] | | 150 | 150 | | 150 | |
| $V_{half,h}$ [mV] | | -81 | -90 | | -90 | |
| $E_L$ [mV] | | -70 | -70 | | -70 | |
| $E_{Na}$ [mV] | | 50 | 50 | | 50 | |
| $E_K$ [mV] | | -80 | -80 | | -80 | |
| $E_h$ [mV] | | -10 | -10 | | -10 | |
| $E_{Ca}$ [mV] | | 140 | 140 | | 140 | |

**Table 5.**
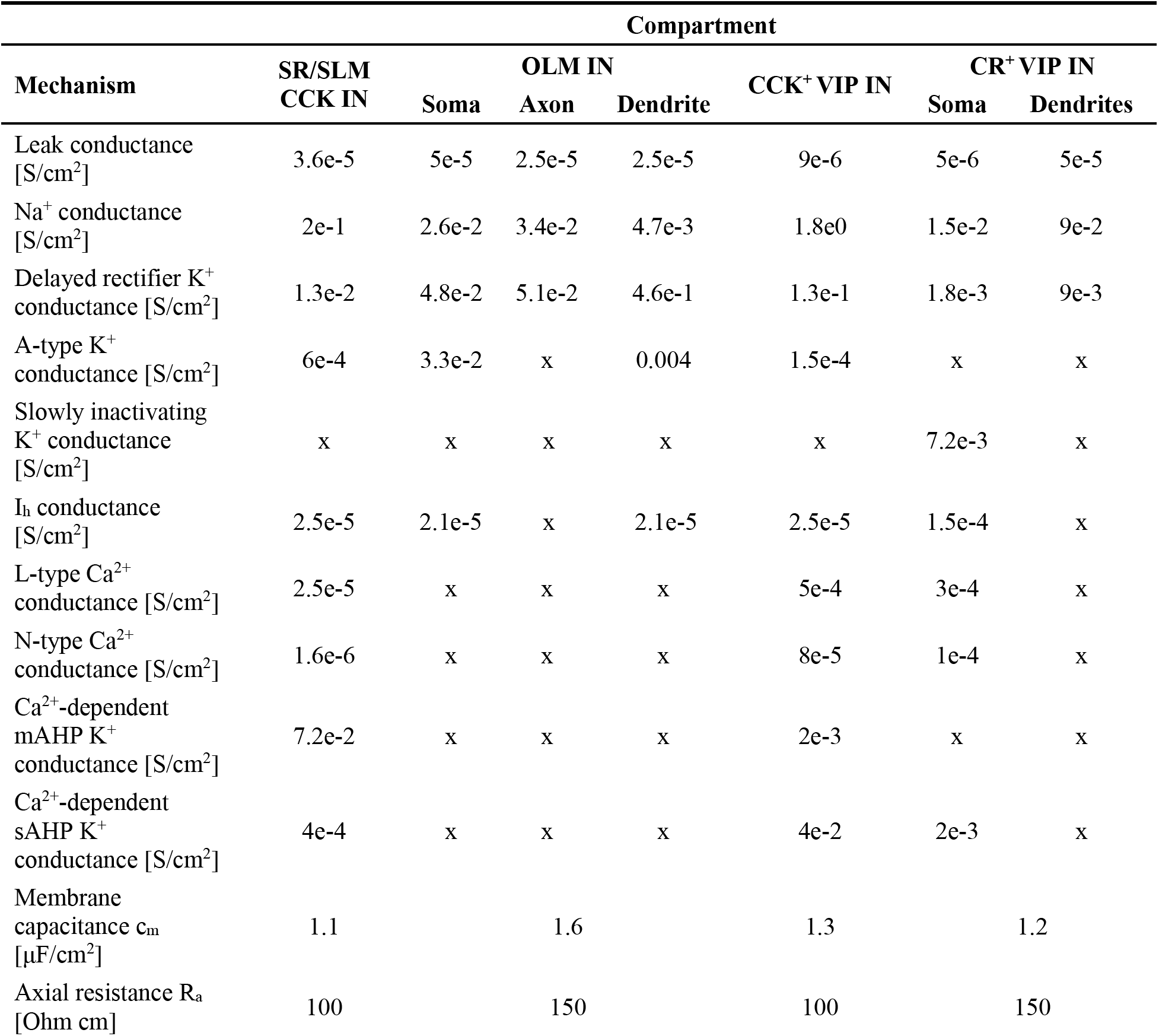
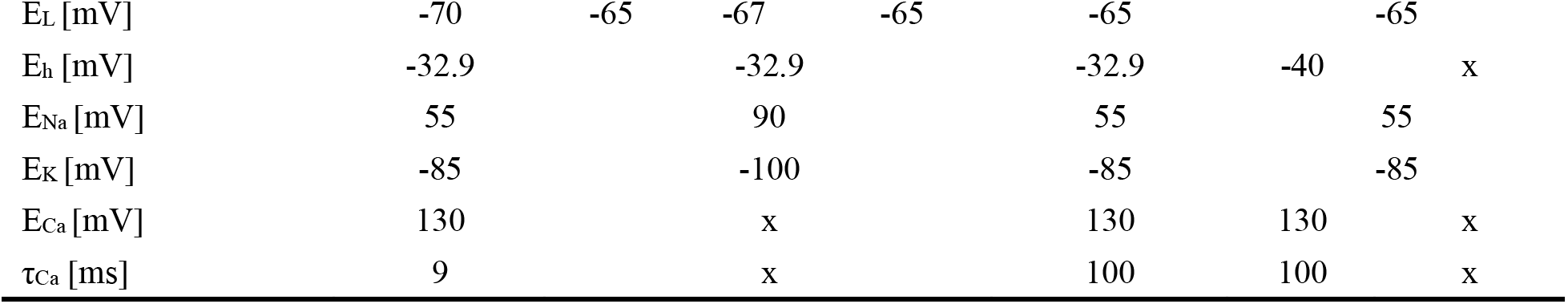
Passive parameters and active ionic conductances for all interneurons and their compartments when applicable.

#### OLM IN model cell

The OLM IN had four compartments, which included a Na^+^ current, a delayed rectifier K^+^ current, an A-type K^+^ current, and an h-current (**Table 5, Figure S12**). The OLM IN received excitatory connections from CA1 PN in their basal dendrites and inhibitory connections from the CR^+^ VIP IN in their soma. The number of excitatory synapses from LEC was drawn from a Poisson distribution, with lambda equals to 0.3, to explain the low activation and no induction of spike shown in experiments (**Figure 4G-I, S12B-C**). The number of excitatory connections from CA1 PN was drawn from an exponential distribution with mean equals to 10. Using this distribution, we generated activity on the OLM IN when using the Poisson protocol (see sections below). The OLM IN also received an inhibitory connection from the CR^+^ VIP IN, the number drawn from an exponential distribution with mean equals to 60, a value sufficient to induce a hyperpolarization to the OLM IN soma.

#### CCK^+^ VIP IN model cell

The CCK^+^ VIP IN had 17 compartments, containing a leak conductance, a sodium current, a fast- delayed rectifier K^+^ current, an A-type K^+^ current, L- and N-type Ca^2+^ currents, a Ca^2+^-dependent K^+^ current, and a Ca^2+^- and voltage-dependent K^+^ current (**Table 5, Figure S12**). To replicate the irregular firing which was experimentally observed, we tuned the calcium influx of this model, making the decay of Ca^2+^ current slower (**Figure S12A)**. The CCK^+^ VIP IN received excitatory connections from LEC to the distal SLM dendrites and made inhibitory synapses onto the CA1 PN soma. The number of excitatory connections was drawn from a Poisson distribution with mean equals 21, while in 16% of the trials (n = 10 x 200 trials), the CCK^+^ VIP IN received no connections from LEC (**Figure 5D-F, S12B-C**).

#### CR^+^ VIP IN model cell

The CR^+^ VIP IN consisted of 17 compartments, including mechanisms for slow K^+^ current, fast Ca^2+^-activated K^+^ current, and N-type Ca^2+^ current (**Table 5, Figure S12**). Each CR^+^ VIP IN received excitatory input from LEC. The number of excitatory connections from LEC was drawn from a Poisson distribution with a mean equal to 5, while in 9% of the trials (n = 10 x 200 trials), the CR^+^ VIP IN did not receive any connection from LEC (**Figure 5D-F, S12B-C**).

#### Validation

Passive (intrinsic) and active (spiking) properties of each neuronal type were validated against experimental data. For the validation, we used the frequency vs. injected current (f-I) curve and the sag-ratio as a function of injected currents (hyperpolarization) (**Figure S2**).

#### Modeling Synapses

AMPA, NMDA and GABA_A,_ and GABA_B_ synapses were included in the model. NMDA synapses were used only in CA1 PNs. The interneurons received excitatory afferents via AMPA receptors. AMPA, NMDA, GABA_A,_ and GABA_B_ synapses were simulated as a sum of two exponentials. All synaptic properties (i.e., conductance g, rise time τ_r_, decay time τ_d_, synaptic delay) were validated with experimental evidence (see **Figures 4, S4, S5**).

#### AMPA and GABA synapses

The AMPA, GABA_A,_ and GABA_B_ synapses were simulated as a two-state kinetic scheme, described by the rise and decay phases. The following equations give the synaptic current through these synapses:

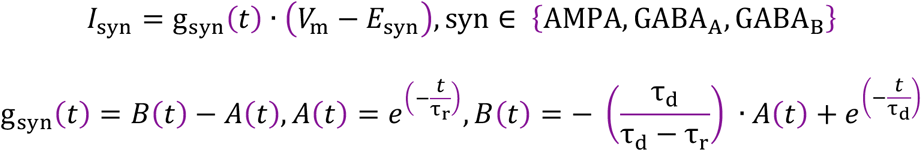

#### NMDA synapse

The NMDA synapse was simulated as a two-state kinetic scheme involving a factor that represents the voltage dependency. The corresponding equations were:

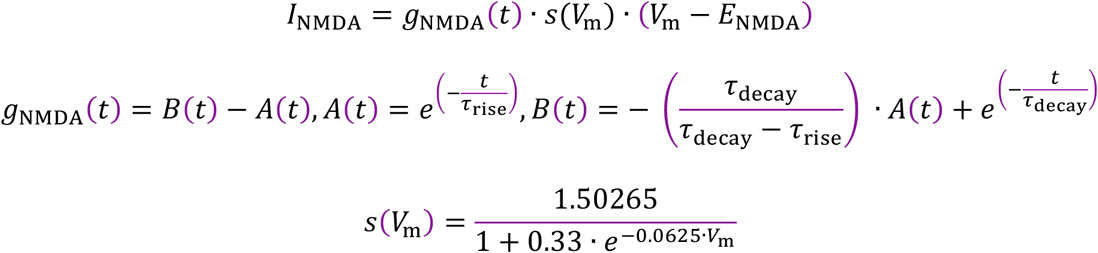

where *V*_m_ denotes the membrane voltage.

In order to validate the synaptic weights, we used the experimental data shown in **Figures 4**, **S4**, and **S5**. All synaptic properties are summarized in **Table 6**.

**Table 6.** Synaptic properties of the canonical circuitry.

| From/to | CA1 PN<sub>s</sub> |  | SR/SLM<br>CCK IN<sub>s</sub> | OLM IN<sub>s</sub> | CCK<sup>+</sup> VIP<br>IN<sub>s</sub> | CR<sup>+</sup> VIP<br>IN<sub>s</sub> |
| --- | --- | --- | --- | --- | --- | --- |
| **LEC LIII** | AMPA | NMDA | AMPA | AMPA | AMPA | AMPA |
| g (nS) | 2.0e-4 | 2.0e-4 | 1.5e-4 | x | 3.0e-4 | 3.0e-4 |
| $\tau_r$ (ms) | 0.5 | 2.3 | 2.0 | x | 2.0 | 2.0 |
| $\tau_d$ (ms) | 3.0 | 100 | 6.3 | x | 6.3 | 6.3 |
| **CA1 PN<sub>s</sub>** | AMPA |  | AMPA | AMPA | AMPA | AMPA |
| g (nS) | 4.9e-4 |  | 1.9e-3 | 2.0e-4 | 5.0e-4 | 5.0e-4 |
| $\tau_r$ (ms) | 0.1 | | 0.11 | 0.3 | 0.5 | 0.5 |
| $\tau_d$ (ms) | 9.5 | | 0.25 | 0.6 | 3.0 | 3.0 |
| **SR/SLM**<br>**CCK IN<sub>s</sub>** | GABA<sub>A</sub> | GABA<sub>B</sub> | GABA<sub>A</sub> | GABA<sub>A</sub> | GABA<sub>A</sub> | GABA<sub>A</sub> |
| g (nS) | 5.1e-4 | 1.1e-4 | 5.1e-4 | 2.0e-5 | 8.0e-4 | x |
| $\tau_r$ (ms) | 0.11 | 35 | 0.29 | 1.0 | 0.29 | x |
| $\tau_d$(ms) | 9.70 | 100 | 2.67 | 8.0 | 2.67 | x |
| **OLM IN<sub>s</sub>** | GABA<sub>A</sub> | GABA<sub>B</sub> | GABA<sub>A</sub> | GABA<sub>A</sub> | GABA<sub>A</sub> | GABA<sub>A</sub> |
| g (nS) | 3.0e-4 | 3.0e-4 | 1.1e-4 | 1.2e-3 | 1.2e-3 | x |
| $\tau_r$ (ms) | 0.13 | 35 | 0.6 | 0.25 | 0.73 | x |
| $\tau_d$ (ms) | 11.0 | 100 | 15 | 7.50 | 20.20 | x |
| **CCK<sup>+</sup> VIP**<br>**IN<sub>s</sub>** | GABA<sub>A</sub> |  | GABA<sub>A</sub> | GABA<sub>A</sub> | GABA<sub>A</sub> | GABA<sub>A</sub> |
| g (nS) | 5.2e-4 |  | 7.0e-4 | x | 4.5e-4 | x |
| $\tau_r$ (ms) | 0.3 | | 0.43 | x | 0.43 | x |
| $\tau_d$ (ms) | 4.2 | | 4.49 | x | 4.49 | x |
| **CR<sup>+</sup> VIP**<br>**IN<sub>s</sub>** | GABA<sub>A</sub> |  | GABA<sub>A</sub> | GABA<sub>A</sub> | GABA<sub>A</sub> | GABA<sub>A</sub> |
| g (nS) | x |  | x | 7.0e-4 | x | x |
| $\tau_r$ (ms) | x | | x | 1.0 | x | x |
| $\tau_d$ (ms) | x | | x | 8.0 | x | x |

#### Single-cell simulations

Various synapses, excitatory and inhibitory, were randomly placed in the CA1 PN model. The excitatory synapses representing the synapses from LEC were placed in the apical tuft at the SLM layer. In contrast, the inhibitory synapses were placed either in SLM, SR, or SP based on random distribution. The number of excitatory synapses was drawn from an exponential distribution with a mean of 12. Each excitatory synapse consisted of one AMPA and one NMDA component. The 60% of inhibitory synapses were located at SLM (distal dendritic inhibition), 20% at SR (proximal dendritic inhibition), and 20% were located at SP (somatic inhibition). The synapses located at SR and SLM were GABA_A_ and GABA_B_ based on uniform distribution. The total number of inhibitory synapses was selected from a random Gaussian distribution with a mean of 40 and std 4. The simulation duration was set at t=2sec, and the activation of the pre-synaptic axons was set at t=700ms. For the various deletion protocols, we set the respective weights to zero. All other parameters remained the same so that to compare the conditions pairwise.

#### Canonical microcircuit

The CA1 canonical circuitry consisted of the following cell types: 1 CA1 PN, 1 SR/SLM CCK IN, 1 OLM IN, 1 CCK^+^ VIP IN, and 1 CR^+^ VIP IN. The LEC input was given to all neuronal cells with several synapses drawn from distributions, as explained above. All neuronal models were held at their resting membrane potential. At t=700ms, we stimulated the circuitry with a single spike. The simulation duration was set to 2sec. For the various deletion protocols, we set the weights of the afferents of the deleted neurons to zero. The rest parameters remained the same across the deletion so that the results were paired. We repeated the simulations using a Poisson-like, theta-filtered spike train. The theta filter equation is:

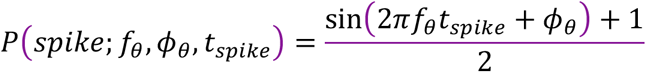

where *f*_θ_ is the theta cycle frequency set to 8Hz, *φ*_θ_ is the phase of the cycle set to zero, and *t_spike_* denotes the spike-time, which came from the Poisson distribution. If the probability is greater than 0.7, a spike is generated for this specific train. Also, 20% of out-of-theta-cycle spikes were included in the final spike train to have noise in the spike trains.

## Author contributions

OMB and JB designed the study and wrote the manuscript. OMB performed the viral injections, slice electrophysiology experiments, immunohistochemistry, confocal imaging, and analysis. SC and PP designed and implemented the computational model and ran the interneuron deletion simulations.

## Acknowledgements

This work was supported by the NIH BRAIN INITIATIVE 1R01NS109994, NIH 1R01NS109362-01, McKnight Scholar Award in Neuroscience, Klingenstein-Simons Fellowship Award in Neuroscience, Alfred P. Sloan Research Fellowship, Whitehall Research Grant, American Epilepsy Society Junior Investigator Award, Blas Frangione Young Investigator Research Grant, New York University Whitehead Fellowship for Junior Faculty in Biomedical and Biological Sciences, and Leon Levy Foundation Award to JB; NIH 5T32NS86750 to OB; FET OPEN NEUREKA (GA 863254) of the European Commission, NIH 1R01MH124867-01, and a Visiting Fellowship of the Einstein Foundation Berlin (EVF-2019-508) grant to PP.

We thank Drs. Tanvi Butola, Adam Carter, Dmitri Chklovskii, Melissa Hernandez Frausto, Robert Machold, Jason Moore, John Rinzel, Vincent Robert, and Richard Tsien for helpful discussion on earlier versions of the manuscript. We thank Drs. Bernardo Rudy and Chiung-Yin Chung for generously providing the triple transgenic VIP IN subtype mice.

## REFERENCES

Acsady, L., Arabadzisz, D., and Freund, T.F. (1996a). Correlated morphological and neurochemical features identify different subsets of vasoactive intestinal polypeptide-immunoreactive interneurons in rat hippocampus. Neuroscience 73, 299–315.

Acsady, L., Gorcs, T.J., and Freund, T.F. (1996b). Different populations of vasoactive intestinal polypeptide-immunoreactive interneurons are specialized to control pyramidal cells or interneurons in the hippocampus. Neuroscience 73, 317–334.

Armstrong, C., Krook-Magnuson, E., and Soltesz, I. (2012). Neurogliaform and Ivy Cells: A Major Family of nNOS Expressing GABAergic Neurons. Front Neural Circuits 6, 23.

Augusto, E., and Gambino, F. (2019). Can NMDA Spikes Dictate Computations of Local Networks and Behavior? Front Mol Neurosci 12, 238.

Basu, J., and Siegelbaum, S.A. (2015). The Corticohippocampal Circuit, Synaptic Plasticity, and Memory. Cold Spring Harb Perspect Biol 7.

Basu, J., Srinivas, K.V., Cheung, S.K., Taniguchi, H., Huang, Z.J., and Siegelbaum, S.A. (2013). A cortico-hippocampal learning rule shapes inhibitory microcircuit activity to enhance hippocampal information flow. Neuron 79, 1208–1221.

Basu, J., Zaremba, J.D., Cheung, S.K., Hitti, F.L., Zemelman, B.V., Losonczy, A., and Siegelbaum, S.A. (2016). Gating of hippocampal activity, plasticity, and memory by entorhinal cortex long-range inhibition. Science 351, aaa5694.

Beaulieu-Laroche, L., and Harnett, M.T. (2018). Dendritic Spines Prevent Synaptic Voltage Clamp. Neuron 97, 75–82 e73.

Bezaire, M.J., Raikov, I., Burk, K., Vyas, D., and Soltesz, I. (2016). Interneuronal mechanisms of hippocampal theta oscillations in a full-scale model of the rodent CA1 circuit. Elife 5.

Bezaire, M.J., and Soltesz, I. (2013). Quantitative assessment of CA1 local circuits: knowledge base for interneuron-pyramidal cell connectivity. Hippocampus 23, 751–785.

Bi, G.Q., and Poo, M.M. (1998). Synaptic modifications in cultured hippocampal neurons: dependence on spike timing, synaptic strength, and postsynaptic cell type. J Neurosci 18, 10464–10472.

Bittner, K.C., Grienberger, C., Vaidya, S.P., Milstein, A.D., Macklin, J.J., Suh, J., Tonegawa, S., and Magee, J.C. (2015). Conjunctive input processing drives feature selectivity in hippocampal CA1 neurons. Nat Neurosci 18, 1133–1142.

Bittner, K.C., Milstein, A.D., Grienberger, C., Romani, S., and Magee, J.C. (2017). Behavioral time scale synaptic plasticity underlies CA1 place fields. Science 357, 1033–1036.

Bloss, E.B., Cembrowski, M.S., Karsh, B., Colonell, J., Fetter, R.D., and Spruston, N. (2016). Structured Dendritic Inhibition Supports Branch-Selective Integration in CA1 Pyramidal Cells. Neuron 89, 1016–1030.

Branco, T., and Hausser, M. (2010). The single dendritic branch as a fundamental functional unit in the nervous system. Curr Opin Neurobiol 20, 494–502.

Branco, T., and Hausser, M. (2011). Synaptic integration gradients in single cortical pyramidal cell dendrites. Neuron 69, 885–892.

Brun, V.H., Leutgeb, S., Wu, H.Q., Schwarcz, R., Witter, M.P., Moser, E.I., and Moser, M.B. (2008). Impaired spatial representation in CA1 after lesion of direct input from entorhinal cortex. Neuron 57, 290–302.

Brun, V.H., Otnass, M.K., Molden, S., Steffenach, H.A., Witter, M.P., Moser, M.B., and Moser, E.I. (2002). Place cells and place recognition maintained by direct entorhinal-hippocampal circuitry. Science 296, 2243–2246.

Buzsaki, G., and Moser, E.I. (2013). Memory, navigation and theta rhythm in the hippocampal-entorhinal system. Nat Neurosci 16, 130–138.

Chamberland, S., and Topolnik, L. (2012). Inhibitory control of hippocampal inhibitory neurons. Front Neurosci 6, 165.

Chavlis, S., and Poirazi, P. (2021). Drawing inspiration from biological dendrites to empower artificial neural networks. Curr Opin Neurobiol 70, 1–10.

Chittajallu, R., Wester, J.C., Craig, M.T., Barksdale, E., Yuan, X.Q., Akgul, G., Fang, C., Collins, D., Hunt, S., Pelkey, K.A., et al. (2017). Afferent specific role of NMDA receptors for the circuit integration of hippocampal neurogliaform cells. Nat Commun 8, 152.

Chuong, A.S., Miri, M.L., Busskamp, V., Matthews, G.A., Acker, L.C., Sorensen, A.T., Young, A., Klapoetke, N.C., Henninger, M.A., Kodandaramaiah, S.B., et al. (2014). Noninvasive optical inhibition with a red-shifted microbial rhodopsin. Nat Neurosci 17, 1123–1129.

Cichon, J., and Gan, W.B. (2015). Branch-specific dendritic Ca(2+) spikes cause persistent synaptic plasticity. Nature 520, 180–185.

Cope, D.W., Maccaferri, G., Marton, L.F., Roberts, J.D., Cobden, P.M., and Somogyi, P. (2002). Cholecystokinin-immunopositive basket and Schaffer collateral-associated interneurones target different domains of pyramidal cells in the CA1 area of the rat hippocampus. Neuroscience 109, 63–80.

Cui, Z., Gerfen, C.R., and Young, W.S., 3rd (2013). Hypothalamic and other connections with dorsal CA2 area of the mouse hippocampus. J Comp Neurol 521, 1844–1866.

Cutsuridis, V., Cobb, S., and Graham, B.P. (2010). Encoding and retrieval in a model of the hippocampal CA1 microcircuit. Hippocampus 20, 423–446.

Cutsuridis, V., and Hasselmo, M. (2012). GABAergic contributions to gating, timing, and phase precession of hippocampal neuronal activity during theta oscillations. Hippocampus 22, 1597–1621.

Cutsuridis, V., and Poirazi, P. (2015). A computational study on how theta modulated inhibition can account for the long temporal windows in the entorhinal-hippocampal loop. Neurobiol Learn Mem 120, 69–83.

Daigle, T.L., Madisen, L., Hage, T.A., Valley, M.T., Knoblich, U., Larsen, R.S., Takeno, M.M., Huang, L., Gu, H., Larsen, R., et al. (2018). A Suite of Transgenic Driver and Reporter Mouse Lines with Enhanced Brain-Cell-Type Targeting and Functionality. Cell 174, 465–480 e422.

David, C., Schleicher, A., Zuschratter, W., and Staiger, J.F. (2007). The innervation of parvalbumin-containing interneurons by VIP-immunopositive interneurons in the primary somatosensory cortex of the adult rat. Eur J Neurosci 25, 2329–2340.

Del Pino, I., Brotons-Mas, J.R., Marques-Smith, A., Marighetto, A., Frick, A., Marin, O., and Rico, B. (2017). Abnormal wiring of CCK(+) basket cells disrupts spatial information coding. Nat Neurosci 20, 784–792.

Deshmukh, S.S., and Knierim, J.J. (2011). Representation of non-spatial and spatial information in the lateral entorhinal cortex. Front Behav Neurosci 5, 69.

Desmond, N.L., Scott, C.A., Jane, J.A., Jr., and Levy, W.B. (1994). Ultrastructural identification of entorhinal cortical synapses in CA1 stratum lacunosum-moleculare of the rat. Hippocampus 4, 594–600.

Doan, T.P., Lagartos-Donate, M.J., Nilssen, E.S., Ohara, S., and Witter, M.P. (2019). Convergent Projections from Perirhinal and Postrhinal Cortices Suggest a Multisensory Nature of Lateral, but Not Medial, Entorhinal Cortex. Cell Rep 29, 617–627 e617.

Dudman, J.T., Tsay, D., and Siegelbaum, S.A. (2007). A role for synaptic inputs at distal dendrites: instructive signals for hippocampal long-term plasticity. Neuron 56, 866–879.

Dudok, B., Klein, P.M., Hwaun, E., Lee, B.R., Yao, Z., Fong, O., Bowler, J.C., Terada, S., Sparks, F.T., Szabo, G.G., et al. (2021a). Alternating sources of perisomatic inhibition during behavior. Neuron 109, 997–1012 e1019.

Dudok, B., Szoboszlay, M., Paul, A., Klein, P.M., Liao, Z., Hwaun, E., Szabo, G.G., Geiller, T., Vancura, B., Wang, B.S., et al. (2021b). Recruitment and inhibitory action of hippocampal axo-axonic cells during behavior. Neuron 109, 3838–3850 e3838.

Dvorak-Carbone, H., and Schuman, E.M. (1999). Patterned activity in stratum lacunosum moleculare inhibits CA1 pyramidal neuron firing. J Neurophysiol 82, 3213–3222.

Ferguson, K.A., Chatzikalymniou, A.P., and Skinner, F.K. (2017). Combining Theory, Model, and Experiment to Explain How Intrinsic Theta Rhythms Are Generated in an In Vitro Whole Hippocampus Preparation without Oscillatory Inputs. eNeuro 4.

Francavilla, R., Luo, X., Magnin, E., Tyan, L., and Topolnik, L. (2015). Coordination of dendritic inhibition through local disinhibitory circuits. Front Synaptic Neurosci 7, 5.

Francavilla, R., Villette, V., Luo, X., Chamberland, S., Munoz-Pino, E., Camire, O., Wagner, K., Kis, V., Somogyi, P., and Topolnik, L. (2018). Connectivity and network state-dependent recruitment of long-range VIP-GABAergic neurons in the mouse hippocampus. Nat Commun 9, 5043.

Frank, L.M., Brown, E.N., and Wilson, M.A. (2001). A comparison of the firing properties of putative excitatory and inhibitory neurons from CA1 and the entorhinal cortex. J Neurophysiol 86, 2029–2040.

Freund, T.F., and Buzsaki, G. (1996). Interneurons of the hippocampus. Hippocampus 6, 347–470.

Gambino, F., Pages, S., Kehayas, V., Baptista, D., Tatti, R., Carleton, A., and Holtmaat, A. (2014). Sensory-evoked LTP driven by dendritic plateau potentials in vivo. Nature 515, 116–119.

Gasparini, S., Migliore, M., and Magee, J.C. (2004). On the initiation and propagation of dendritic spikes in CA1 pyramidal neurons. J Neurosci 24, 11046–11056.

Geiller, T., Vancura, B., Terada, S., Troullinou, E., Chavlis, S., Tsagkatakis, G., Tsakalides, P., Ocsai, K., Poirazi, P., Rozsa, B.J., et al. (2020). Large-Scale 3D Two-Photon Imaging of Molecularly Identified CA1 Interneuron Dynamics in Behaving Mice. Neuron 108, 968–983 e969.

Gidon, A., and Segev, I. (2012). Principles governing the operation of synaptic inhibition in dendrites. Neuron 75, 330–341.

Glickfeld, L.L., Roberts, J.D., Somogyi, P., and Scanziani, M. (2009). Interneurons hyperpolarize pyramidal cells along their entire somatodendritic axis. Nat Neurosci 12, 21–23.

Glickfeld, L.L., and Scanziani, M. (2006). Distinct timing in the activity of cannabinoid-sensitive and cannabinoid-insensitive basket cells. Nat Neurosci 9, 807–815.

Golding, N.L., Jung, H.Y., Mickus, T., and Spruston, N. (1999). Dendritic calcium spike initiation and repolarization are controlled by distinct potassium channel subtypes in CA1 pyramidal neurons. J Neurosci 19, 8789–8798.

Golding, N.L., Mickus, T.J., Katz, Y., Kath, W.L., and Spruston, N. (2005). Factors mediating powerful voltage attenuation along CA1 pyramidal neuron dendrites. J Physiol 568, 69–82.

Golding, N.L., and Spruston, N. (1998). Dendritic sodium spikes are variable triggers of axonal action potentials in hippocampal CA1 pyramidal neurons. Neuron 21, 1189–1200.

Golding, N.L., Staff, N.P., and Spruston, N. (2002). Dendritic spikes as a mechanism for cooperative long-term potentiation. Nature 418, 326–331.

Gomez Gonzalez, J.F., Mel, B.W., and Poirazi, P. (2011). Distinguishing Linear vs. Non-Linear Integration in CA1 Radial Oblique Dendrites: It’s about Time. Front Comput Neurosci 5, 44.

Grienberger, C., Milstein, A.D., Bittner, K.C., Romani, S., and Magee, J.C. (2017). Inhibitory suppression of heterogeneously tuned excitation enhances spatial coding in CA1 place cells. Nat Neurosci 20, 417–426.

Guet-McCreight, A., Skinner, F.K., and Topolnik, L. (2020). Common Principles in Functional Organization of VIP/Calretinin Cell-Driven Disinhibitory Circuits Across Cortical Areas. Front Neural Circuits 14, 32.

Hales, J.B., Schlesiger, M.I., Leutgeb, J.K., Squire, L.R., Leutgeb, S., and Clark, R.E. (2014). Medial entorhinal cortex lesions only partially disrupt hippocampal place cells and hippocampus-dependent place memory. Cell Rep 9, 893–901.

Hargreaves, E.L., Rao, G., Lee, I., and Knierim, J.J. (2005). Major dissociation between medial and lateral entorhinal input to dorsal hippocampus. Science 308, 1792–1794.

Hausser, M., and Mel, B. (2003). Dendrites: bug or feature? Curr Opin Neurobiol 13, 372–383.

Hausser, M., Spruston, N., and Stuart, G.J. (2000). Diversity and dynamics of dendritic signaling. Science 290, 739–744.

He, M., Tucciarone, J., Lee, S., Nigro, M.J., Kim, Y., Levine, J.M., Kelly, S.M., Krugikov, I., Wu, P., Chen, Y., et al. (2016). Strategies and Tools for Combinatorial Targeting of GABAergic Neurons in Mouse Cerebral Cortex. Neuron 91, 1228–1243.

Hsu, C.L., Zhao, X., Milstein, A.D., and Spruston, N. (2018). Persistent Sodium Current Mediates the Steep Voltage Dependence of Spatial Coding in Hippocampal Pyramidal Neurons. Neuron 99, 147–162 e148.

Igarashi, K.M., Lu, L., Colgin, L.L., Moser, M.B., and Moser, E.I. (2014). Coordination of entorhinal-hippocampal ensemble activity during associative learning. Nature 510, 143–147.

Ito, H.T., and Schuman, E.M. (2007). Frequency-dependent gating of synaptic transmission and plasticity by dopamine. Front Neural Circuits 1, 1.

Jadi, M., Polsky, A., Schiller, J., and Mel, B.W. (2012). Location-dependent effects of inhibition on local spiking in pyramidal neuron dendrites. PLoS Comput Biol 8, e1002550.

Jarsky, T., Roxin, A., Kath, W.L., and Spruston, N. (2005). Conditional dendritic spike propagation following distal synaptic activation of hippocampal CA1 pyramidal neurons. Nat Neurosci 8, 1667–1676.

Jia, H., Rochefort, N.L., Chen, X., and Konnerth, A. (2010). Dendritic organization of sensory input to cortical neurons in vivo. Nature 464, 1307–1312.

Jun, H., Bramian, A., Soma, S., Saito, T., Saido, T.C., and Igarashi, K.M. (2020). Disrupted Place Cell Remapping and Impaired Grid Cells in a Knockin Model of Alzheimer’s Disease. Neuron 107, 1095–1112 e1096.

Kajiwara, R., Wouterlood, F.G., Sah, A., Boekel, A.J., Baks-te Bulte, L.T., and Witter, M.P. (2008). Convergence of entorhinal and CA3 inputs onto pyramidal neurons and interneurons in hippocampal area CA1--an anatomical study in the rat. Hippocampus 18, 266–280.

Kamondi, A., Acsady, L., and Buzsaki, G. (1998a). Dendritic spikes are enhanced by cooperative network activity in the intact hippocampus. J Neurosci 18, 3919–3928.

Kamondi, A., Acsady, L., Wang, X.J., and Buzsaki, G. (1998b). Theta oscillations in somata and dendrites of hippocampal pyramidal cells in vivo: activity-dependent phase-precession of action potentials. Hippocampus 8, 244–261.

Kerlin, A., Mohar, B., Flickinger, D., MacLennan, B.J., Dean, M.B., Davis, C., Spruston, N., and Svoboda, K. (2019). Functional clustering of dendritic activity during decision-making. Elife 8.

Kerr, K.M., Agster, K.L., Furtak, S.C., and Burwell, R.D. (2007). Functional neuroanatomy of the parahippocampal region: the lateral and medial entorhinal areas. Hippocampus 17, 697–708.

Kim, Y., Hsu, C.L., Cembrowski, M.S., Mensh, B.D., and Spruston, N. (2015). Dendritic sodium spikes are required for long-term potentiation at distal synapses on hippocampal pyramidal neurons. Elife 4.

Kitamura, T., Pignatelli, M., Suh, J., Kohara, K., Yoshiki, A., Abe, K., and Tonegawa, S. (2014). Island cells control temporal association memory. Science 343, 896–901.

Kitamura, T., Sun, C., Martin, J., Kitch, L.J., Schnitzer, M.J., and Tonegawa, S. (2015). Entorhinal Cortical Ocean Cells Encode Specific Contexts and Drive Context-Specific Fear Memory. Neuron 87, 1317–1331.

Klausberger, T. (2009). GABAergic interneurons targeting dendrites of pyramidal cells in the CA1 area of the hippocampus. Eur J Neurosci 30, 947–957.

Klausberger, T., Marton, L.F., O’Neill, J., Huck, J.H., Dalezios, Y., Fuentealba, P., Suen, W.Y., Papp, E., Kaneko, T., Watanabe, M., et al. (2005). Complementary roles of cholecystokinin- and parvalbumin-expressing GABAergic neurons in hippocampal network oscillations. J Neurosci 25, 9782–9793.

Klausberger, T., and Somogyi, P. (2008). Neuronal diversity and temporal dynamics: the unity of hippocampal circuit operations. Science 321, 53–57.

Kleinlogel, S., Feldbauer, K., Dempski, R.E., Fotis, H., Wood, P.G., Bamann, C., and Bamberg, E. (2011). Ultra light-sensitive and fast neuronal activation with the Ca(2)+-permeable channelrhodopsin CatCh. Nat Neurosci 14, 513–518.

Kuruvilla, M.V., Wilson, D.I.G., and Ainge, J.A. (2020). Lateral entorhinal cortex lesions impair both egocentric and allocentric object-place associations. Brain Neurosci Adv 4, 2398212820939463.

Lacaille, J.C., and Schwartzkroin, P.A. (1988). Stratum lacunosum-moleculare interneurons of hippocampal CA1 region. II. Intrasomatic and intradendritic recordings of local circuit synaptic interactions. J Neurosci 8, 1411–1424.

Larkum, M.E., Nevian, T., Sandler, M., Polsky, A., and Schiller, J. (2009). Synaptic integration in tuft dendrites of layer 5 pyramidal neurons: a new unifying principle. Science 325, 756–760.

Larkum, M.E., Zhu, J.J., and Sakmann, B. (1999). A new cellular mechanism for coupling inputs arriving at different cortical layers. Nature 398, 338–341.

Lavzin, M., Rapoport, S., Polsky, A., Garion, L., and Schiller, J. (2012). Nonlinear dendritic processing determines angular tuning of barrel cortex neurons in vivo. Nature 490, 397–401.

Leao, R.N., Mikulovic, S., Leao, K.E., Munguba, H., Gezelius, H., Enjin, A., Patra, K., Eriksson, A., Loew, L.M., Tort, A.B., et al. (2012). OLM interneurons differentially modulate CA3 and entorhinal inputs to hippocampal CA1 neurons. Nat Neurosci 15, 1524–1530.

Lee, J.H., Durand, R., Gradinaru, V., Zhang, F., Goshen, I., Kim, D.S., Fenno, L.E., Ramakrishnan, C., and Deisseroth, K. (2010). Global and local fMRI signals driven by neurons defined optogenetically by type and wiring. Nature 465, 788–792.

Lee, J.Y., Jun, H., Soma, S., Nakazono, T., Shiraiwa, K., Dasgupta, A., Nakagawa, T., Xie, J.L., Chavez, J., Romo, R., et al. (2021). Dopamine facilitates associative memory encoding in the entorhinal cortex. Nature 598, 321–326.

Lee, S., Kruglikov, I., Huang, Z.J., Fishell, G., and Rudy, B. (2013). A disinhibitory circuit mediates motor integration in the somatosensory cortex. Nat Neurosci 16, 1662–1670.

Leitner, F.C., Melzer, S., Lutcke, H., Pinna, R., Seeburg, P.H., Helmchen, F., and Monyer, H. (2016). Spatially segregated feedforward and feedback neurons support differential odor processing in the lateral entorhinal cortex. Nat Neurosci 19, 935–944.

Li, B., Suutari, B.S., Sun, S.D., Luo, Z., Wei, C., Chenouard, N., Mandelberg, N.J., Zhang, G., Wamsley, B., Tian, G., et al. (2020). Neuronal Inactivity Co-opts LTP Machinery to Drive Potassium Channel Splicing and Homeostatic Spike Widening. Cell 181, 1547–1565 e1515.

Li, Y., Xu, J., Liu, Y., Zhu, J., Liu, N., Zeng, W., Huang, N., Rasch, M.J., Jiang, H., Gu, X., et al. (2017). A distinct entorhinal cortex to hippocampal CA1 direct circuit for olfactory associative learning. Nat Neurosci 20, 559–570.

Lin, J.Y. (2011). A user’s guide to channelrhodopsin variants: features, limitations and future developments. Exp Physiol 96, 19–25.

Losonczy, A., and Magee, J.C. (2006). Integrative properties of radial oblique dendrites in hippocampal CA1 pyramidal neurons. Neuron 50, 291–307.

Losonczy, A., Makara, J.K., and Magee, J.C. (2008). Compartmentalized dendritic plasticity and input feature storage in neurons. Nature 452, 436–441.

Lovett-Barron, M., Kaifosh, P., Kheirbek, M.A., Danielson, N., Zaremba, J.D., Reardon, T.R., Turi, G.F., Hen, R., Zemelman, B.V., and Losonczy, A. (2014). Dendritic inhibition in the hippocampus supports fear learning. Science 343, 857–863.

Lovett-Barron, M., Turi, G.F., Kaifosh, P., Lee, P.H., Bolze, F., Sun, X.H., Nicoud, J.F., Zemelman, B.V., Sternson, S.M., and Losonczy, A. (2012). Regulation of neuronal input transformations by tunable dendritic inhibition. Nat Neurosci 15, 423–430, S421-423.

Lu, L., Leutgeb, J.K., Tsao, A., Henriksen, E.J., Leutgeb, S., Barnes, C.A., Witter, M.P., Moser, M.B., and Moser, E.I. (2013). Impaired hippocampal rate coding after lesions of the lateral entorhinal cortex. Nat Neurosci 16, 1085–1093.

Luna, V.M., Anacker, C., Burghardt, N.S., Khandaker, H., Andreu, V., Millette, A., Leary, P., Ravenelle, R., Jimenez, J.C., Mastrodonato, A., et al. (2019). Adult-born hippocampal neurons bidirectionally modulate entorhinal inputs into the dentate gyrus. Science 364, 578–583.

Madisen, L., Garner, A.R., Shimaoka, D., Chuong, A.S., Klapoetke, N.C., Li, L., van der Bourg, A., Niino, Y., Egolf, L., Monetti, C., et al. (2015). Transgenic mice for intersectional targeting of neural sensors and effectors with high specificity and performance. Neuron 85, 942–958.

Madisen, L., Zwingman, T.A., Sunkin, S.M., Oh, S.W., Zariwala, H.A., Gu, H., Ng, L.L., Palmiter, R.D., Hawrylycz, M.J., Jones, A.R., et al. (2010). A robust and high-throughput Cre reporting and characterization system for the whole mouse brain. Nat Neurosci 13, 133–140.

Markram, H., Lubke, J., Frotscher, M., and Sakmann, B. (1997). Regulation of synaptic efficacy by coincidence of postsynaptic APs and EPSPs. Science 275, 213–215.

Masurkar, A.V., Srinivas, K.V., Brann, D.H., Warren, R., Lowes, D.C., and Siegelbaum, S.A. (2017). Medial and Lateral Entorhinal Cortex Differentially Excite Deep versus Superficial CA1 Pyramidal Neurons. Cell Rep 18, 148–160.

Megias, M., Emri, Z., Freund, T.F., and Gulyas, A.I. (2001). Total number and distribution of inhibitory and excitatory synapses on hippocampal CA1 pyramidal cells. Neuroscience 102, 527–540.

Melzer, S., Michael, M., Caputi, A., Eliava, M., Fuchs, E.C., Whittington, M.A., and Monyer, H. (2012). Long-range-projecting GABAergic neurons modulate inhibition in hippocampus and entorhinal cortex. Science 335, 1506–1510.

Mergenthal, A., Bouteiller, J.C., Yu, G.J., and Berger, T.W. (2020). A Computational Model of the Cholinergic Modulation of CA1 Pyramidal Cell Activity. Front Comput Neurosci 14, 75.

Migliore, M., De Simone, G., and Migliore, R. (2015). Effect of the initial synaptic state on the probability to induce long-term potentiation and depression. Biophys J 108, 1038–1046.

Miles, R., Toth, K., Gulyas, A.I., Hajos, N., and Freund, T.F. (1996). Differences between somatic and dendritic inhibition in the hippocampus. Neuron 16, 815–823.

Milstein, A.D., Bloss, E.B., Apostolides, P.F., Vaidya, S.P., Dilly, G.A., Zemelman, B.V., and Magee, J.C. (2015). Inhibitory Gating of Input Comparison in the CA1 Microcircuit. Neuron 87, 1274–1289.

Miyoshi, G., Hjerling-Leffler, J., Karayannis, T., Sousa, V.H., Butt, S.J., Battiste, J., Johnson, J.E., Machold, R.P., and Fishell, G. (2010). Genetic fate mapping reveals that the caudal ganglionic eminence produces a large and diverse population of superficial cortical interneurons. J Neurosci 30, 1582–1594.

Moore, J.J., Ravassard, P.M., Ho, D., Acharya, L., Kees, A.L., Vuong, C., and Mehta, M.R. (2017). Dynamics of cortical dendritic membrane potential and spikes in freely behaving rats. Science 355.

Moore, J.J., Robert, V., Rashid, S.K., and Basu, J. (2021). Assessing Local and Branch-specific Activity in Dendrites. Neuroscience.

Muller, C., Beck, H., Coulter, D., and Remy, S. (2012). Inhibitory control of linear and supralinear dendritic excitation in CA1 pyramidal neurons. Neuron 75, 851–864.

Murayama, M., Perez-Garci, E., Nevian, T., Bock, T., Senn, W., and Larkum, M.E. (2009). Dendritic encoding of sensory stimuli controlled by deep cortical interneurons. Nature 457, 1137–1141.

Nilssen, E.S., Jacobsen, B., Fjeld, G., Nair, R.R., Blankvoort, S., Kentros, C., and Witter, M.P. (2018). Inhibitory Connectivity Dominates the Fan Cell Network in Layer II of Lateral Entorhinal Cortex. J Neurosci 38, 9712–9727.

Palacios-Filardo, J., Udakis, M., Brown, G.A., Tehan, B.G., Congreve, M.S., Nathan, P.J., Brown, A.J.H., and Mellor, J.R. (2021). Acetylcholine prioritises direct synaptic inputs from entorhinal cortex to CA1 by differential modulation of feedforward inhibitory circuits. Nat Commun 12, 5475.

Palmer, L.M., Schulz, J.M., and Larkum, M.E. (2013). Layer-specific regulation of cortical neurons by interhemispheric inhibition. Commun Integr Biol 6, e23545.

Park, J., Papoutsi, A., Ash, R.T., Marin, M.A., Poirazi, P., and Smirnakis, S.M. (2019). Contribution of apical and basal dendrites to orientation encoding in mouse V1 L2/3 pyramidal neurons. Nat Commun 10, 5372.

Pawelzik, H., Hughes, D.I., and Thomson, A.M. (2002). Physiological and morphological diversity of immunocytochemically defined parvalbumin- and cholecystokinin-positive interneurones in CA1 of the adult rat hippocampus. J Comp Neurol 443, 346–367.

Pedrosa, V., and Clopath, C. (2020). The interplay between somatic and dendritic inhibition promotes the emergence and stabilization of place fields. PLoS Comput Biol 16, e1007955.

Pelkey, K.A., Chittajallu, R., Craig, M.T., Tricoire, L., Wester, J.C., and McBain, C.J. (2017). Hippocampal GABAergic Inhibitory Interneurons. Physiol Rev 97, 1619–1747.

Petreanu, L., Mao, T., Sternson, S.M., and Svoboda, K. (2009). The subcellular organization of neocortical excitatory connections. Nature 457, 1142–1145.

Pfeffer, C.K., Xue, M., He, M., Huang, Z.J., and Scanziani, M. (2013). Inhibition of inhibition in visual cortex: the logic of connections between molecularly distinct interneurons. Nat Neurosci 16, 1068–1076.

Pi, H.J., Hangya, B., Kvitsiani, D., Sanders, J.I., Huang, Z.J., and Kepecs, A. (2013). Cortical interneurons that specialize in disinhibitory control. Nature 503, 521–524.

Pilkiw, M., Insel, N., Cui, Y., Finney, C., Morrissey, M.D., and Takehara-Nishiuchi, K. (2017). Phasic and tonic neuron ensemble codes for stimulus-environment conjunctions in the lateral entorhinal cortex. Elife 6.

Pissadaki, E.K., Sidiropoulou, K., Reczko, M., and Poirazi, P. (2010). Encoding of spatio-temporal input characteristics by a CA1 pyramidal neuron model. PLoS Comput Biol 6, e1001038.

Poirazi, P., Brannon, T., and Mel, B.W. (2003a). Arithmetic of subthreshold synaptic summation in a model CA1 pyramidal cell. Neuron 37, 977–987.

Poirazi, P., Brannon, T., and Mel, B.W. (2003b). Pyramidal neuron as two-layer neural network. Neuron 37, 989–999.

Poirazi, P., and Mel, B.W. (2001). Impact of active dendrites and structural plasticity on the memory capacity of neural tissue. Neuron 29, 779–796.

Pouille, F., and Scanziani, M. (2001). Enforcement of temporal fidelity in pyramidal cells by somatic feed-forward inhibition. Science 293, 1159–1163.

Price, C.J., Cauli, B., Kovacs, E.R., Kulik, A., Lambolez, B., Shigemoto, R., and Capogna, M. (2005). Neurogliaform neurons form a novel inhibitory network in the hippocampal CA1 area. J Neurosci 25, 6775–6786.

Price, C.J., Scott, R., Rusakov, D.A., and Capogna, M. (2008). GABA(B) receptor modulation of feedforward inhibition through hippocampal neurogliaform cells. J Neurosci 28, 6974–6982.

Remondes, M., and Schuman, E.M. (2004). Role for a cortical input to hippocampal area CA1 in the consolidation of a long-term memory. Nature 431, 699–703.

Remy, S., Csicsvari, J., and Beck, H. (2009). Activity-dependent control of neuronal output by local and global dendritic spike attenuation. Neuron 61, 906–916.

Remy, S., and Spruston, N. (2007). Dendritic spikes induce single-burst long-term potentiation. Proc Natl Acad Sci U S A 104, 17192–17197.

Robinson, N.T.M., Descamps, L.A.L., Russell, L.E., Buchholz, M.O., Bicknell, B.A., Antonov, G.K., Lau, J.Y.N., Nutbrown, R., Schmidt-Hieber, C., and Hausser, M. (2020). Targeted Activation of Hippocampal Place Cells Drives Memory-Guided Spatial Behavior. Cell 183, 1586–1599 e1510.

Ruth, R.E., Collier, T.J., and Routtenberg, A. (1988). Topographical relationship between the entorhinal cortex and the septotemporal axis of the dentate gyrus in rats: II. Cells projecting from lateral entorhinal subdivisions. J Comp Neurol 270, 506–516.

Save, E., and Sargolini, F. (2017). Disentangling the Role of the MEC and LEC in the Processing of Spatial and Non-Spatial Information: Contribution of Lesion Studies. Front Syst Neurosci 11, 81.

Schlesiger, M.I., Boublil, B.L., Hales, J.B., Leutgeb, J.K., and Leutgeb, S. (2018). Hippocampal Global Remapping Can Occur without Input from the Medial Entorhinal Cortex. Cell Rep 22, 3152–3159.

Schulz, J.M., Knoflach, F., Hernandez, M.C., and Bischofberger, J. (2018). Dendrite-targeting interneurons control synaptic NMDA-receptor activation via nonlinear alpha5-GABAA receptors. Nat Commun 9, 3576.

Sheffield, M.E., and Dombeck, D.A. (2015). Calcium transient prevalence across the dendritic arbour predicts place field properties. Nature 517, 200–204.

Shuman, T., Aharoni, D., Cai, D.J., Lee, C.R., Chavlis, S., Page-Harley, L., Vetere, L.M., Feng, Y., Yang, C.Y., Mollinedo-Gajate, I., et al. (2020). Breakdown of spatial coding and interneuron synchronization in epileptic mice. Nat Neurosci 23, 229–238.

Sik, A., Penttonen, M., Ylinen, A., and Buzsaki, G. (1995). Hippocampal CA1 interneurons: an in vivo intracellular labeling study. J Neurosci 15, 6651–6665.

Sinha, M., and Narayanan, R. (2021). Active Dendrites and Local Field Potentials: Biophysical Mechanisms and Computational Explorations. Neuroscience.

Sjostrom, P.J., Rancz, E.A., Roth, A., and Hausser, M. (2008). Dendritic excitability and synaptic plasticity. Physiol Rev 88, 769–840.

Smith, S.L., Smith, I.T., Branco, T., and Hausser, M. (2013). Dendritic spikes enhance stimulus selectivity in cortical neurons in vivo. Nature 503, 115–120.

Spruston, N., Jaffe, D.B., and Johnston, D. (1994). Dendritic attenuation of synaptic potentials and currents: the role of passive membrane properties. Trends Neurosci 17, 161–166.

Stuart, G., Schiller, J., and Sakmann, B. (1997). Action potential initiation and propagation in rat neocortical pyramidal neurons. J Physiol 505 ( *Pt 3*), 617–632.

Stuart, G.J., and Spruston, N. (2015). Dendritic integration: 60 years of progress. Nat Neurosci 18, 1713–1721.

Suh, J., Rivest, A.J., Nakashiba, T., Tominaga, T., and Tonegawa, S. (2011). Entorhinal cortex layer III input to the hippocampus is crucial for temporal association memory. Science 334, 1415–1420.

Susaki, E.A., Tainaka, K., Perrin, D., Yukinaga, H., Kuno, A., and Ueda, H.R. (2015). Advanced CUBIC protocols for whole-brain and whole-body clearing and imaging. Nat Protoc 10, 1709–1727.

Takacs, V.T., Szonyi, A., Freund, T.F., Nyiri, G., and Gulyas, A.I. (2015). Quantitative ultrastructural analysis of basket and axo-axonic cell terminals in the mouse hippocampus. Brain Struct Funct 220, 919–940.

Takahashi, H., and Magee, J.C. (2009). Pathway interactions and synaptic plasticity in the dendritic tuft regions of CA1 pyramidal neurons. Neuron 62, 102–111.

Taniguchi, H., He, M., Wu, P., Kim, S., Paik, R., Sugino, K., Kvitsiani, D., Fu, Y., Lu, J., Lin, Y., et al. (2011). A resource of Cre driver lines for genetic targeting of GABAergic neurons in cerebral cortex. Neuron 71, 995–1013.

To, M.S., Honnuraiah, S., and Stuart, G.J. (2021). Voltage Clamp Errors During Estimation of Concurrent Excitatory and Inhibitory Synaptic Input to Neurons with Dendrites. Neuroscience.

Tomko, M., Benuskova, L., and Jedlicka, P. (2021). A new reduced-morphology model for CA1 pyramidal cells and its validation and comparison with other models using HippoUnit. Sci Rep 11, 7615.

Tricoire, L., Pelkey, K.A., Daw, M.I., Sousa, V.H., Miyoshi, G., Jeffries, B., Cauli, B., Fishell, G., and McBain, C.J. (2010). Common origins of hippocampal Ivy and nitric oxide synthase expressing neurogliaform cells. J Neurosci 30, 2165–2176.

Tsao, A., Moser, M.B., and Moser, E.I. (2013). Traces of experience in the lateral entorhinal cortex. Curr Biol 23, 399–405.

Tsao, A., Sugar, J., Lu, L., Wang, C., Knierim, J.J., Moser, M.B., and Moser, E.I. (2018). Integrating time from experience in the lateral entorhinal cortex. Nature 561, 57–62.

Tsien, J.Z., Chen, D.F., Gerber, D., Tom, C., Mercer, E.H., Anderson, D.J., Mayford, M., Kandel, E.R., and Tonegawa, S. (1996). Subregion- and cell type-restricted gene knockout in mouse brain. Cell 87, 1317–1326.

Turi, G.F., Li, W.K., Chavlis, S., Pandi, I., O’Hare, J., Priestley, J.B., Grosmark, A.D., Liao, Z., Ladow, M., Zhang, J.F., et al. (2019). Vasoactive Intestinal Polypeptide-Expressing Interneurons in the Hippocampus Support Goal-Oriented Spatial Learning. Neuron 101, 1150–1165 e1158.

Tyan, L., Chamberland, S., Magnin, E., Camire, O., Francavilla, R., David, L.S., Deisseroth, K., and Topolnik, L. (2014). Dendritic inhibition provided by interneuron-specific cells controls the firing rate and timing of the hippocampal feedback inhibitory circuitry. J Neurosci 34, 4534–4547.

Tzilivaki, A., Kastellakis, G., and Poirazi, P. (2019). Challenging the point neuron dogma: FS basket cells as 2-stage nonlinear integrators. Nat Commun 10, 3664.

Udakis, M., Pedrosa, V., Chamberlain, S.E.L., Clopath, C., and Mellor, J.R. (2020). Interneuron-specific plasticity at parvalbumin and somatostatin inhibitory synapses onto CA1 pyramidal neurons shapes hippocampal output. Nat Commun 11, 4395.

van Groen, T., Miettinen, P., and Kadish, I. (2003). The entorhinal cortex of the mouse: organization of the projection to the hippocampal formation. Hippocampus 13, 133–149.

Wang, C., Chen, X., Lee, H., Deshmukh, S.S., Yoganarasimha, D., Savelli, F., and Knierim, J.J. (2018). Egocentric coding of external items in the lateral entorhinal cortex. Science 362, 945–949.

Whissell, P.D., Bang, J.Y., Khan, I., Xie, Y.F., Parfitt, G.M., Grenon, M., Plummer, N.W., Jensen, P., Bonin, R.P., and Kim, J.C. (2019). Selective Activation of Cholecystokinin-Expressing GABA (CCK-GABA) Neurons Enhances Memory and Cognition. eNeuro 6.

Wilmes, K.A., Sprekeler, H., and Schreiber, S. (2016). Inhibition as a Binary Switch for Excitatory Plasticity in Pyramidal Neurons. PLoS Comput Biol 12, e1004768.

Wilson, D.I., Langston, R.F., Schlesiger, M.I., Wagner, M., Watanabe, S., and Ainge, J.A. (2013). Lateral entorhinal cortex is critical for novel object-context recognition. Hippocampus 23, 352–366.

Witter, M.P., Doan, T.P., Jacobsen, B., Nilssen, E.S., and Ohara, S. (2017). Architecture of the Entorhinal Cortex A Review of Entorhinal Anatomy in Rodents with Some Comparative Notes. Front Syst Neurosci 11, 46.

Witter, M.P., Groenewegen, H.J., Lopes da Silva, F.H., and Lohman, A.H. (1989). Functional organization of the extrinsic and intrinsic circuitry of the parahippocampal region. Prog Neurobiol 33, 161–253.

Woods, N.I., Vaaga, C.E., Chatzi, C., Adelson, J.D., Collie, M.F., Perederiy, J.V., Tovar, K.R., and Westbrook, G.L. (2018). Preferential Targeting of Lateral Entorhinal Inputs onto Newly Integrated Granule Cells. J Neurosci 38, 5843–5853.

Xu, W., and Wilson, D.A. (2012). Odor-evoked activity in the mouse lateral entorhinal cortex. Neuroscience 223, 12–20.

Yeckel, M.F., and Berger, T.W. (1995). Monosynaptic excitation of hippocampal CA1 pyramidal cells by afferents from the entorhinal cortex. Hippocampus 5, 108–114.

Zhang, S.J., Ye, J., Miao, C., Tsao, A., Cerniauskas, I., Ledergerber, D., Moser, M.B., and Moser, E.I. (2013). Optogenetic dissection of entorhinal-hippocampal functional connectivity. Science 340, 1232627.

