## Supplemental Figures for "Lateral entorhinal cortex inputs modulate hippocampal dendritic excitability by recruiting a local disinhibitory microcircuit"

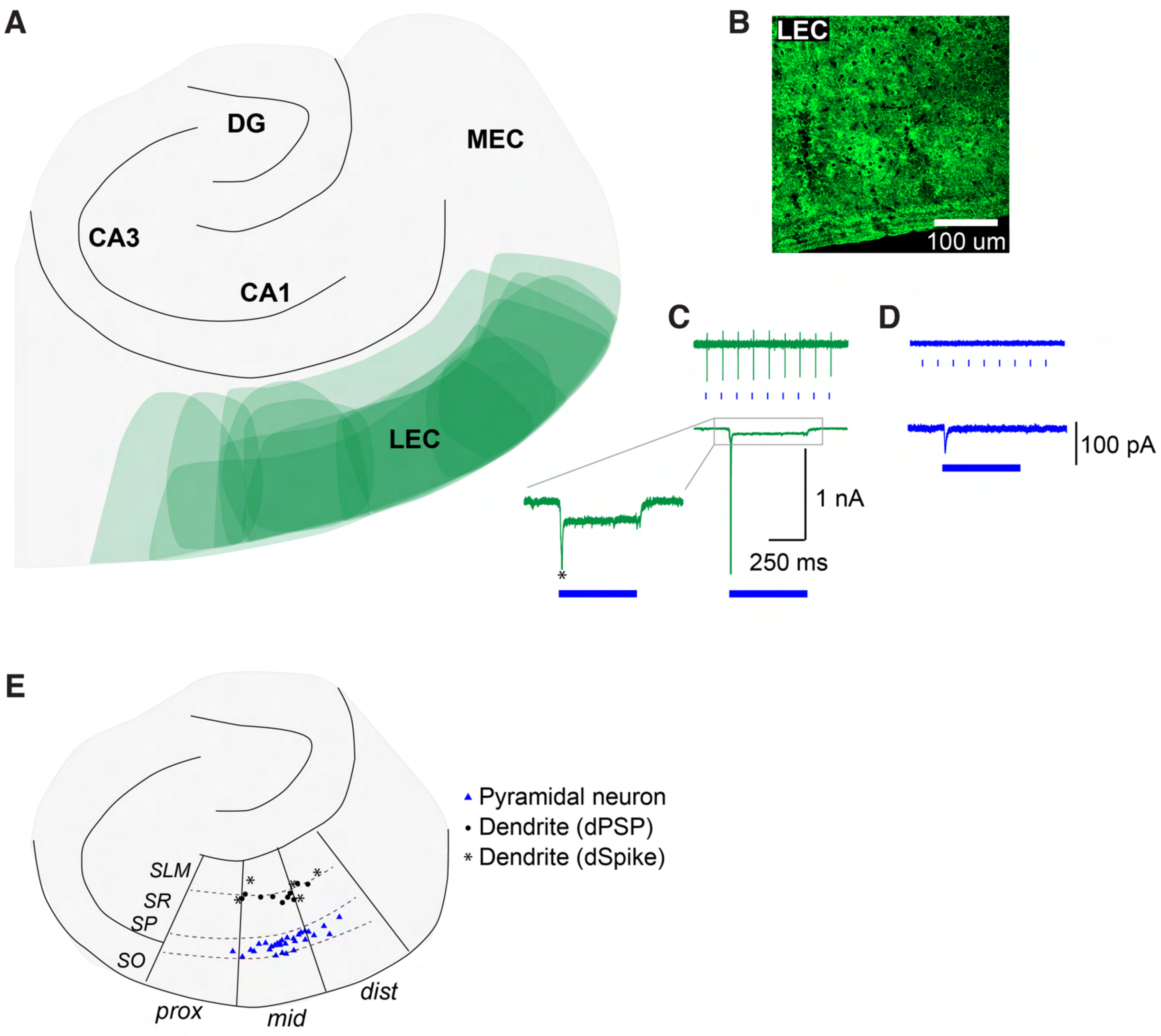

A
DG
MEC
CA3
CA1
LEC
B
LEC
100 um
C
1 nA
250 ms
D
100 pA
E
Pyramidal neuron
Dendrite (dPSP)
Dendrite (dSpike)
SLM
SR
SP
SO
prox
mid
dist


### Figure S1

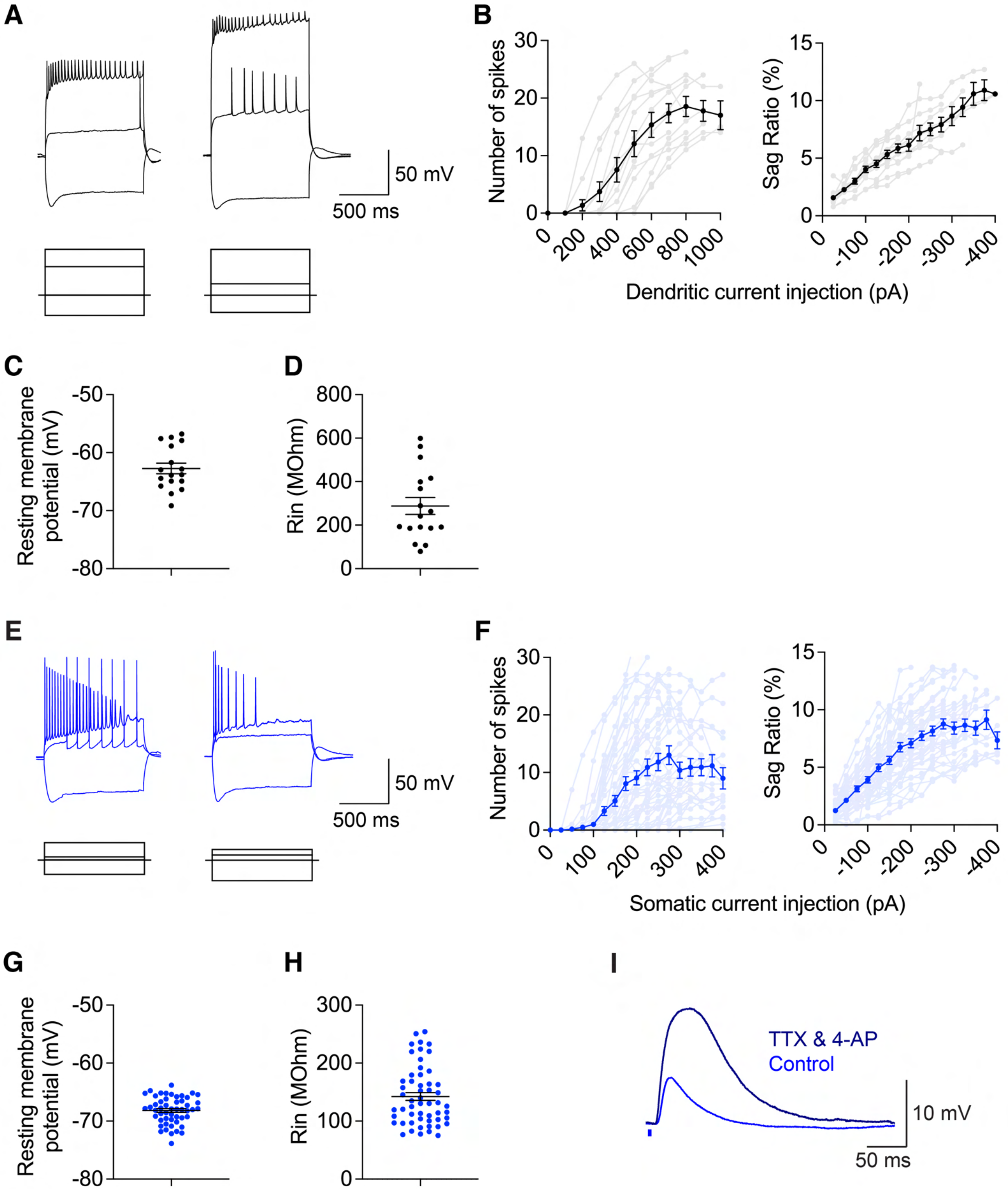
A
50 mV
500 ms
B
Number of spikes
Sag Ratio (%)
Dendritic current injection (pA)
C
Resting membrane potential (mV)
D
Rin (MOhm)
E
50 mV
500 ms
F
Number of spikes
Sag Ratio (%)
Somatic current injection (pA)
G
Resting membrane potential (mV)
H
Rin (MOhm)
I
TTX & 4-AP
Control
10 mV
50 ms

### Figure S2

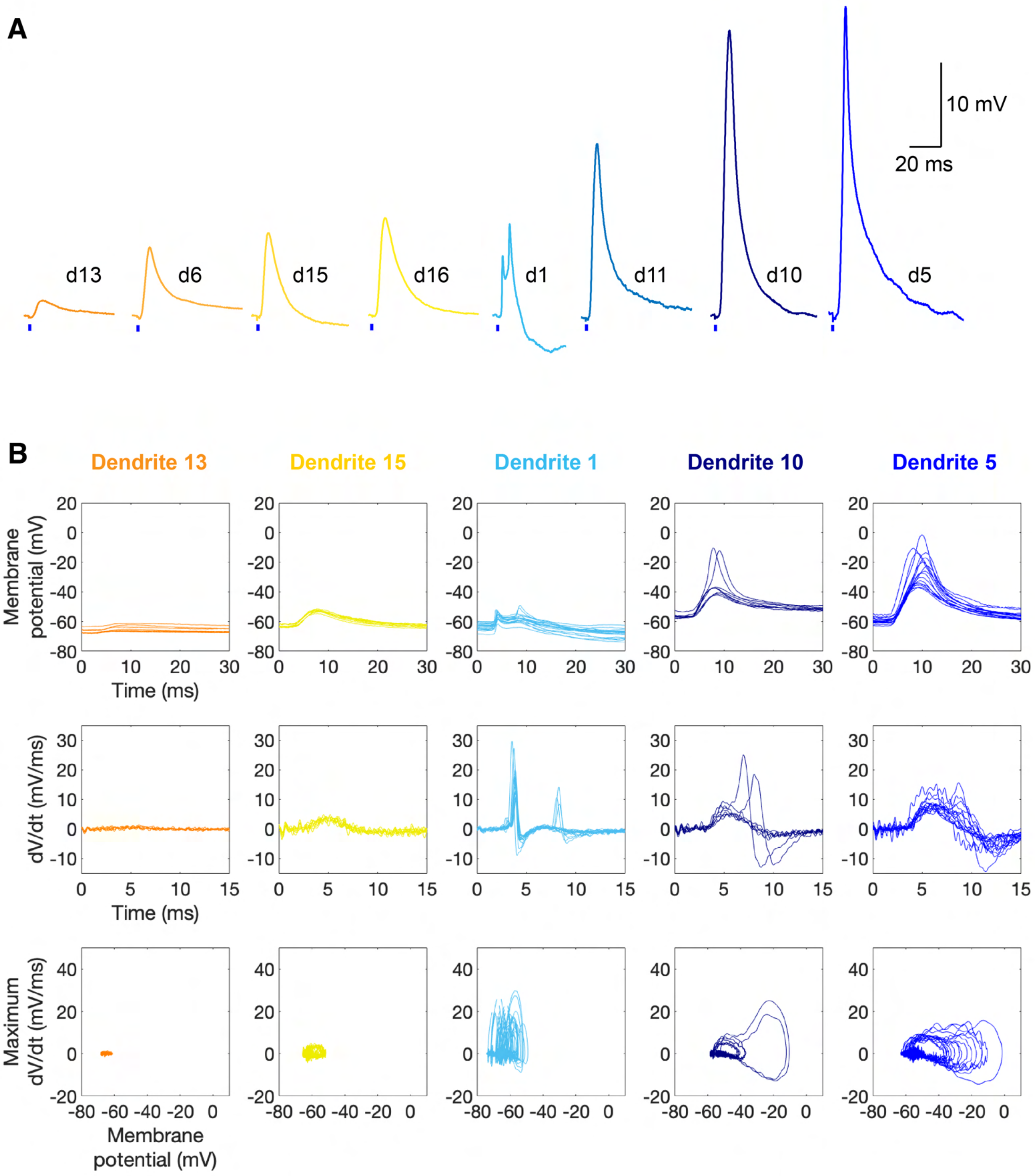
A
10 mV
20 ms
d13
d6
d15
d16
d1
d11
d10
d5
B
Dendrite 13
Dendrite 15
Dendrite 1
Dendrite 10
Dendrite 5
Membrane potential (mV)
Time (ms)
dV/dt (mV/ms)
Maximum dV/dt (mV/ms)
Membrane potential (mV)

### Figure S3

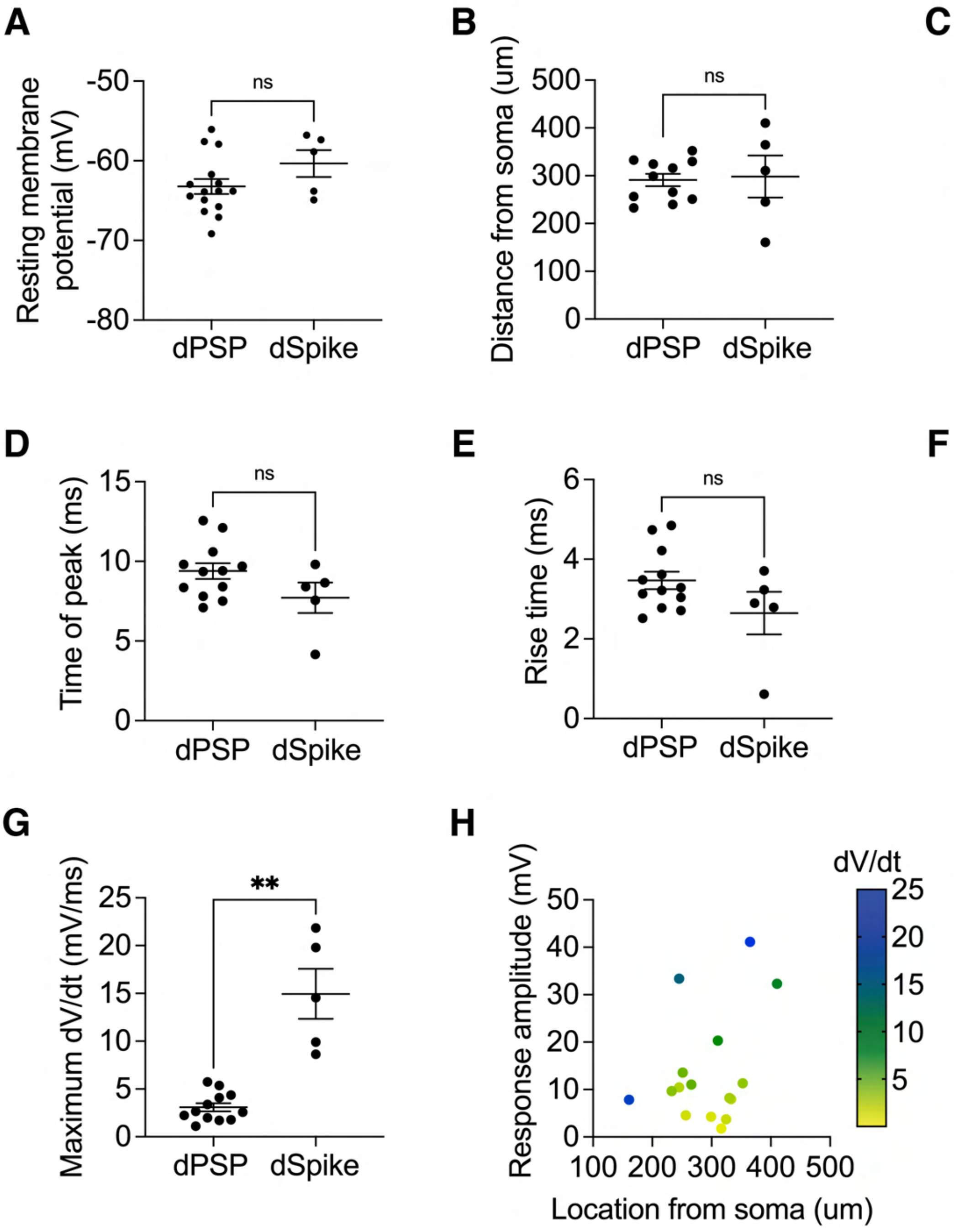

A
Resting membrane potential (mV)
-50
-60
-70
-80
ns
dPSP
dSpike
B
Distance from soma (um)
500
400
300
200
100
0
ns
dPSP
dSpike
C
Peak amplitude (mV)
50
40
30
20
10
0
*
dPSP
dSpike
D
Time of peak (ms)
15
10
5
0
ns
dPSP
dSpike
E
Rise time (ms)
6
4
2
0
ns
dPSP
dSpike
F
Half-width (ms)
40
30
20
10
0
**
dPSP
dSpike
G
Maximum dV/dt (mV/ms)
25
20
15
10
5
0
**
dPSP
dSpike
H
Response amplitude (mV)
50
40
30
20
10
0
100 200 300 400 500
Location from soma (um)
dV/dt
25
20
15
10
5


### Figure S4

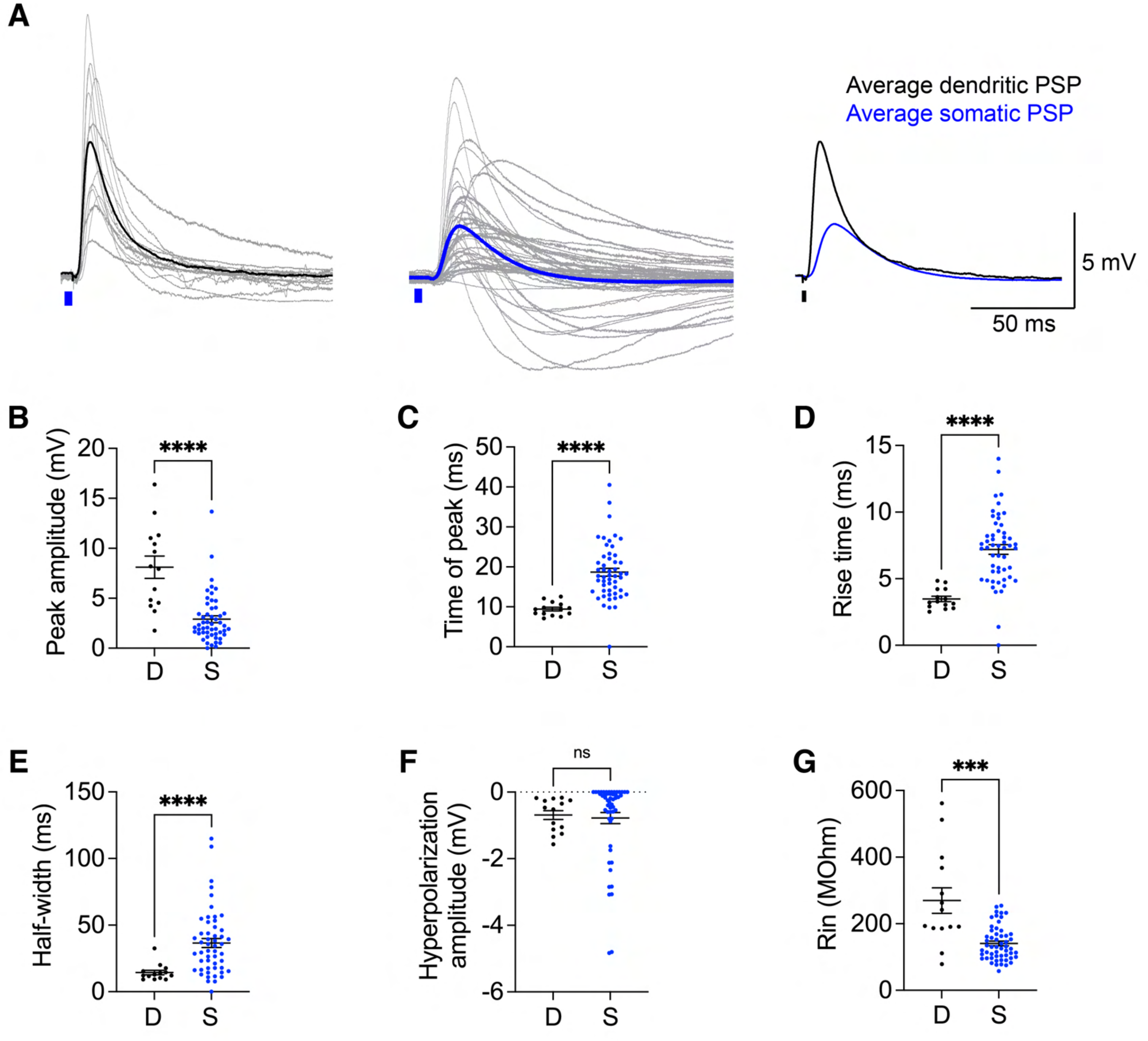
A
Average dendritic PSP
Average somatic PSP
5 mV
50 ms
B
Peak amplitude (mV)
****
D
S
C
Time of peak (ms)
****
D
Rise time (ms)
****
E
Half-width (ms)
****
F
Hyperpolarization amplitude (mV)
ns
G
Rin (MOhm)
***

### Figure S5

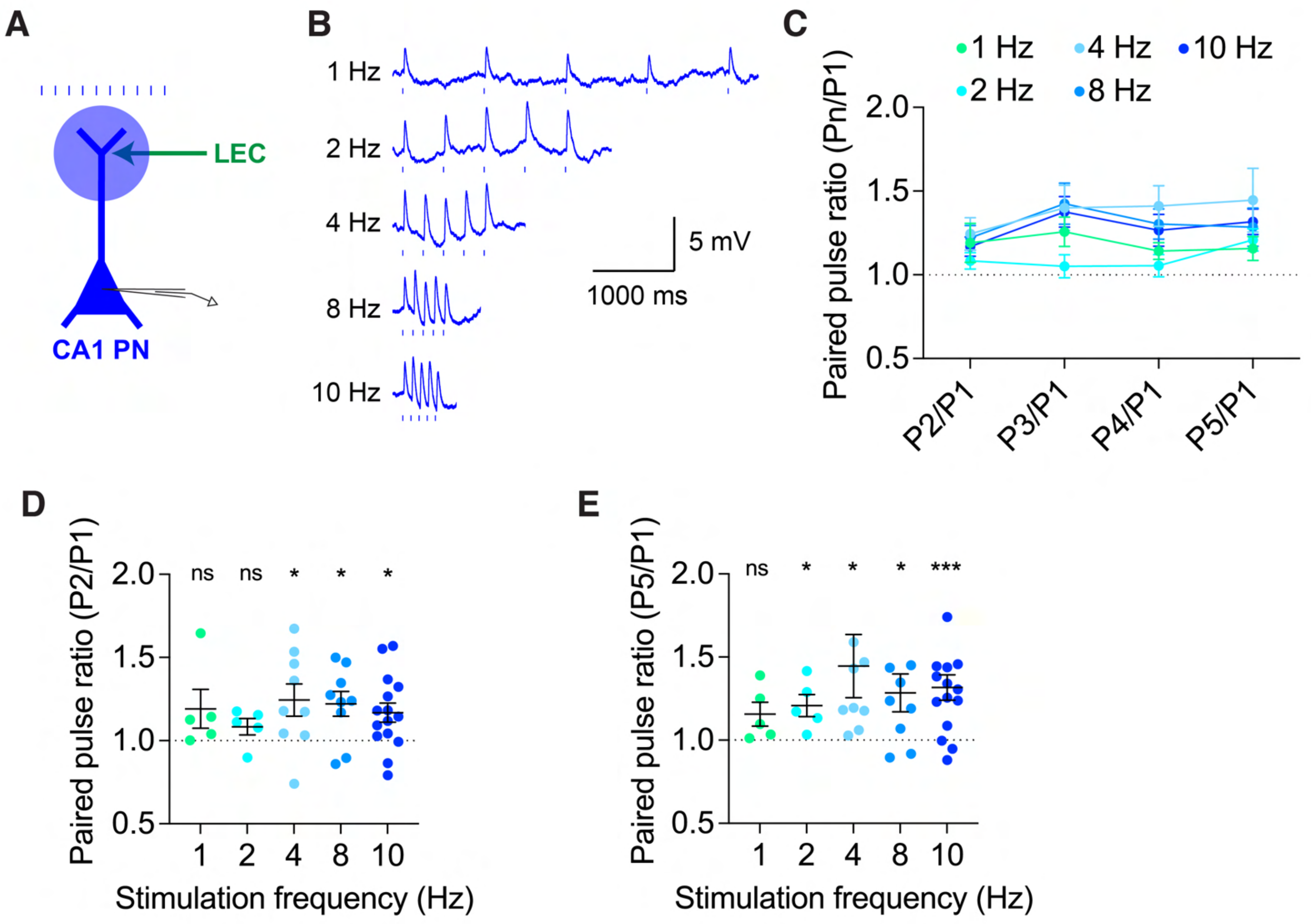
A
LEC
CA1 PN
B
1 Hz
2 Hz
4 Hz
8 Hz
10 Hz
5 mV
1000 ms
C
1 Hz
2 Hz
4 Hz
8 Hz
10 Hz
Paired pulse ratio (Pn/P1)
P2/P1
P3/P1
P4/P1
P5/P1
D
Paired pulse ratio (P2/P1)
ns ns * * *
Stimulation frequency (Hz)
E
Paired pulse ratio (P5/P1)
ns * * * ***
Stimulation frequency (Hz)

### Figure S6

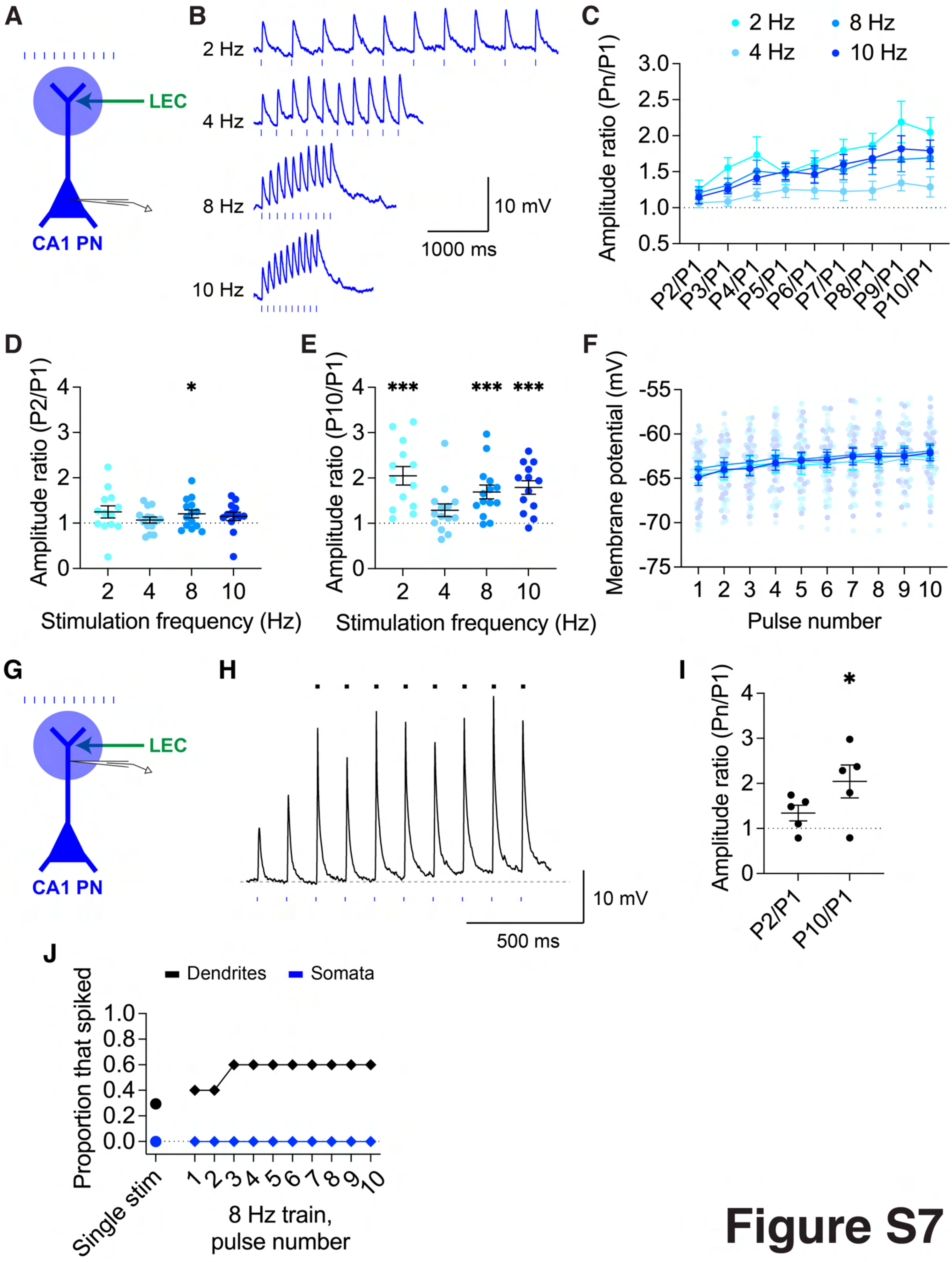
A
LEC
CA1 PN
B
2 Hz
4 Hz
8 Hz
10 Hz
10 mV
1000 ms
C
2 Hz
4 Hz
8 Hz
10 Hz
Amplitude ratio (Pn/P1)
P2/P1
P3/P1
P4/P1
P5/P1
P6/P1
P7/P1
P8/P1
P9/P1
P10/P1
D
Amplitude ratio (P2/P1)
Stimulation frequency (Hz)
E
Amplitude ratio (P10/P1)
Stimulation frequency (Hz)
F
Membrane potential (mV)
Pulse number
G
LEC
CA1 PN
H
10 mV
500 ms
I
Amplitude ratio (Pn/P1)
P2/P1
P10/P1
J
Dendrites
Somata
Proportion that spiked
Single stim
8 Hz train, pulse number

Figure S7

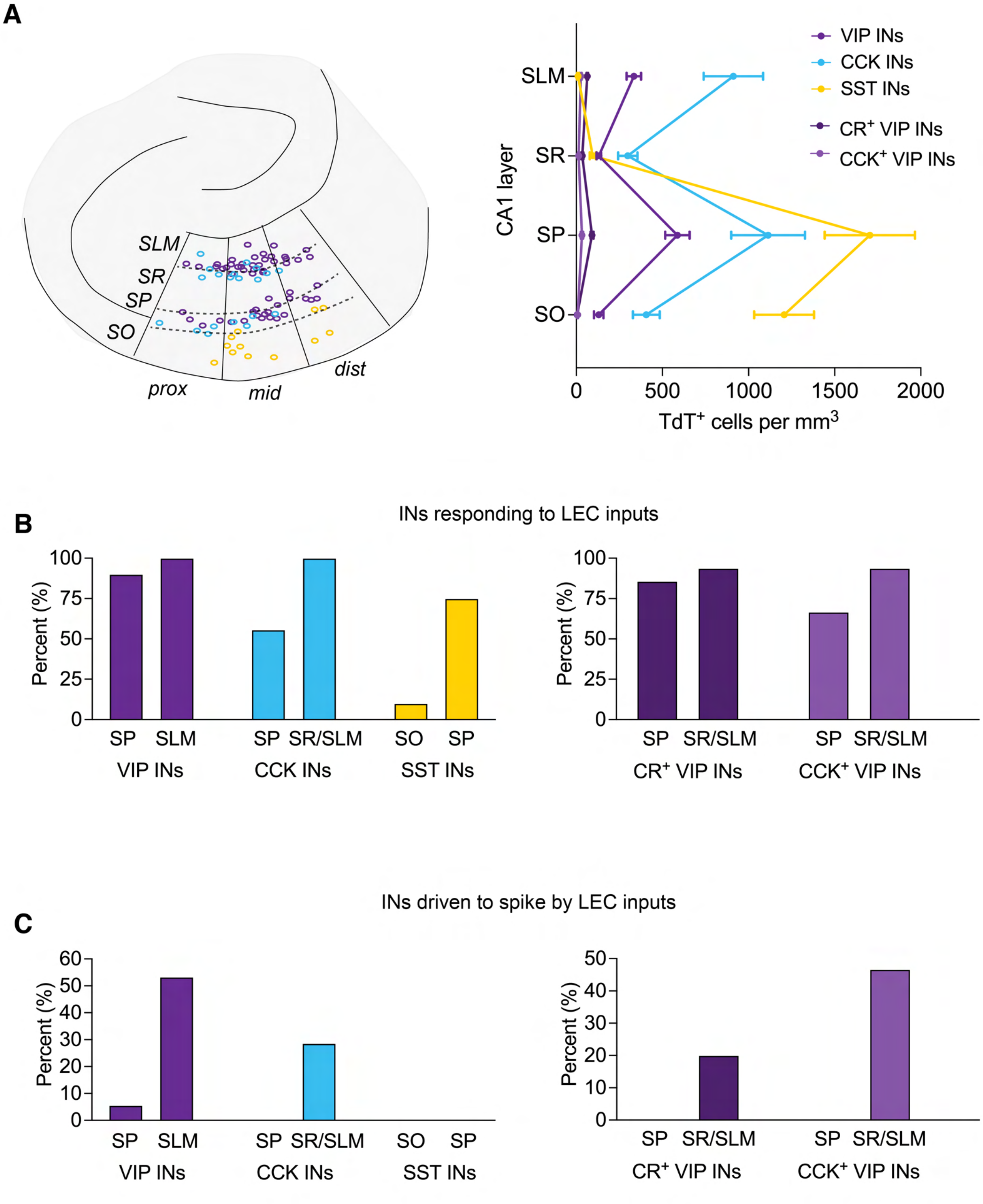
A
SLM
SR
SP
SO
prox
mid
dist
CA1 layer
SLM
SR
SP
SO
VIP INs
CCK INs
SST INs
CR+ VIP INs
CCK+ VIP INs
0
500
1000
1500
2000
TdT+ cells per mm3
INs responding to LEC inputs
B
Percent (%)
100
75
50
25
0
SP
SLM
VIP INs
SP
SR/SLM
CCK INs
SO
SP
SST INs
SP
SR/SLM
CR+ VIP INs
SP
SR/SLM
CCK+ VIP INs
INs driven to spike by LEC inputs
C
Percent (%)
60
50
40
30
20
10
0
SP
SLM
VIP INs
SP
SR/SLM
CCK INs
SO
SP
SST INs
50
40
30
20
10
0
SP
SR/SLM
CR+ VIP INs
SP
SR/SLM
CCK+ VIP INs

**Figure S8**

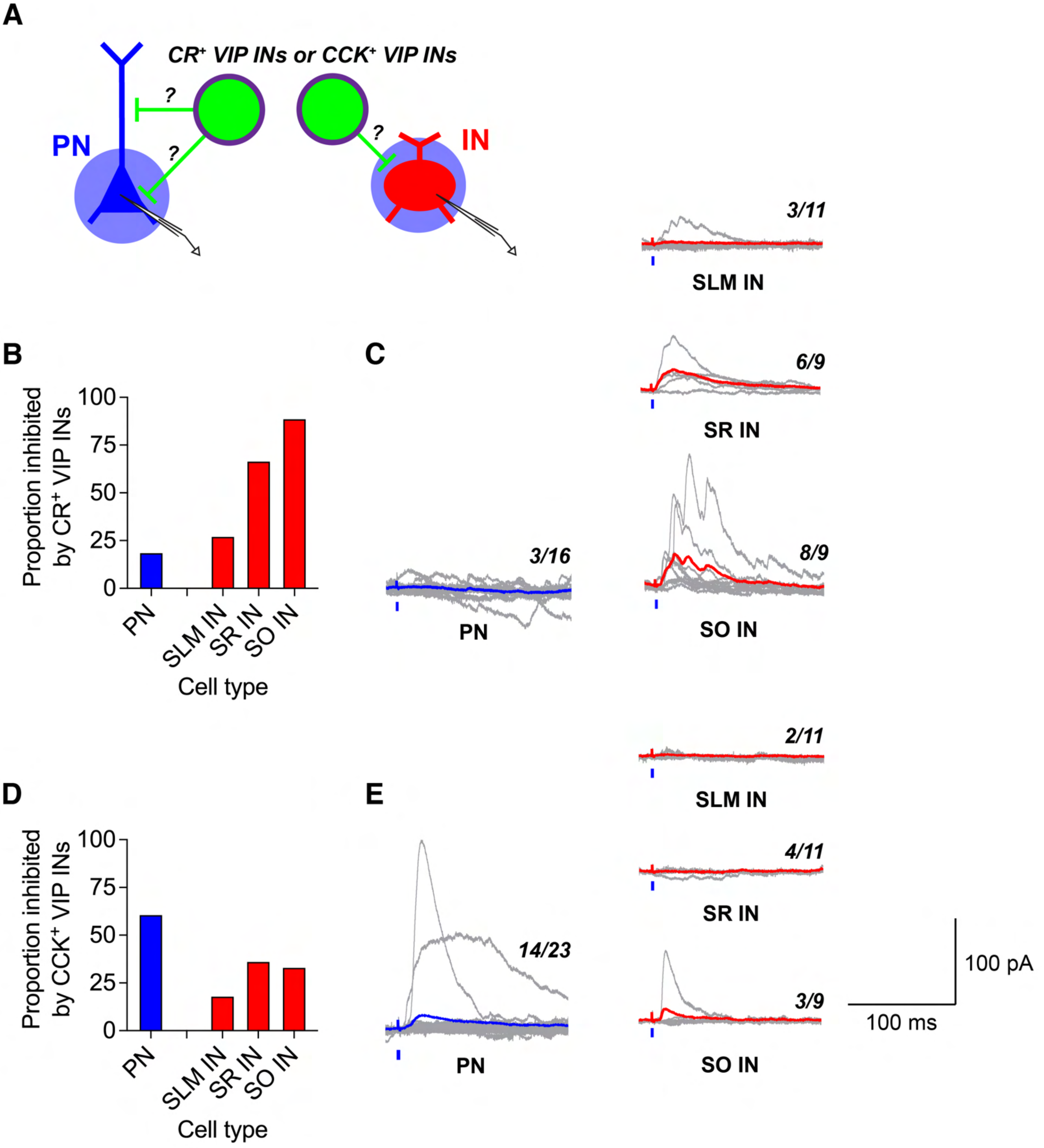
A
CR+ VIP INs or CCK+ VIP INs
PN
IN
?
?
?
B
Proportion inhibited by CR+ VIP INs
100
75
50
25
0
PN
SLM IN
SR IN
SO IN
Cell type
C
3/11
SLM IN
6/9
SR IN
3/16
PN
8/9
SO IN
D
Proportion inhibited by CCK+ VIP INs
100
75
50
25
0
PN
SLM IN
SR IN
SO IN
Cell type
E
2/11
SLM IN
4/11
SR IN
14/23
PN
3/9
SO IN
100 pA
100 ms

### Figure S9

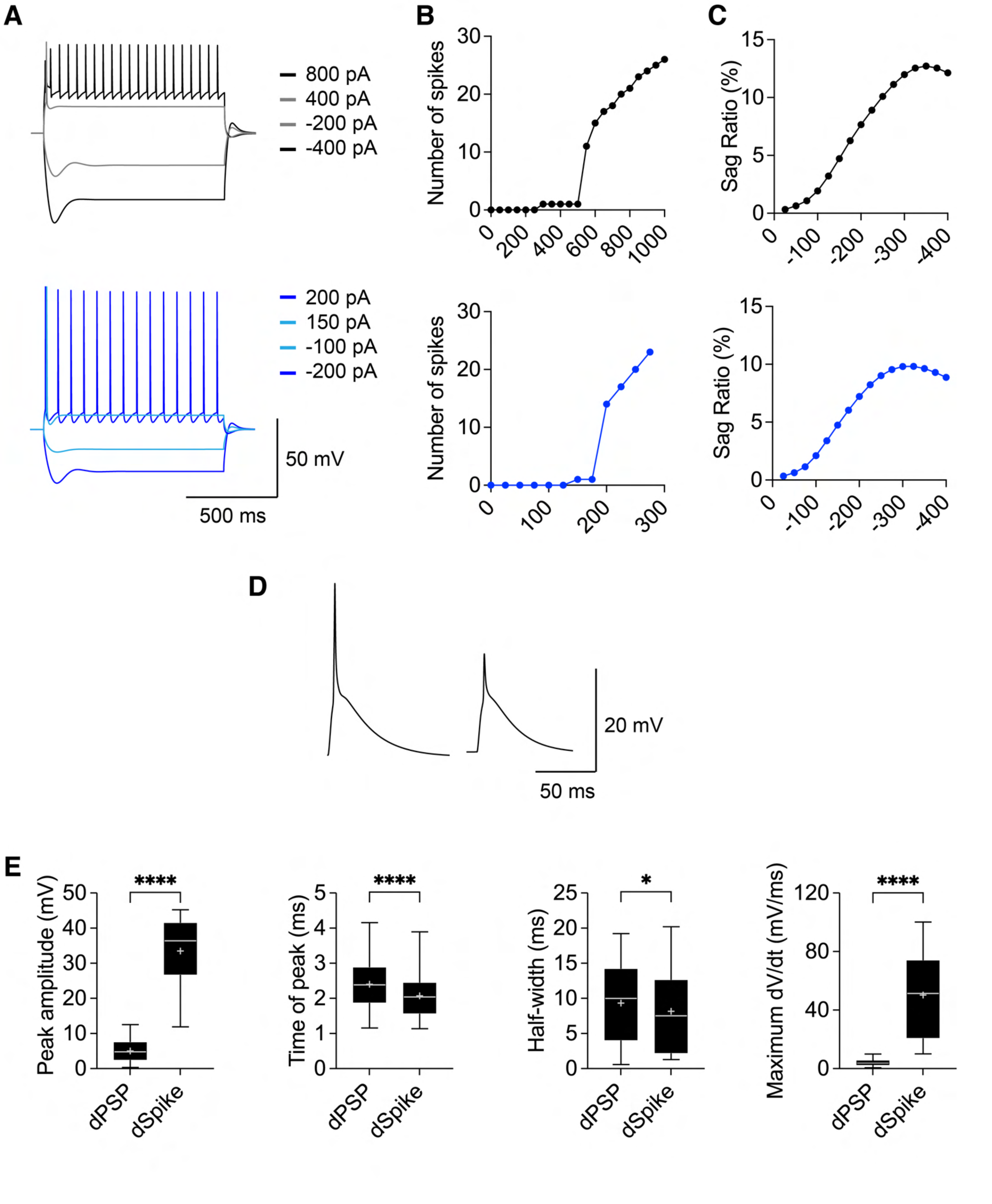
A
800 pA
400 pA
-200 pA
-400 pA
200 pA
150 pA
-100 pA
-200 pA
50 mV
500 ms
B
Number of spikes
C
Sag Ratio (%)
D
20 mV
50 ms
E
Peak amplitude (mV)
Time of peak (ms)
Half-width (ms)
Maximum dV/dt (mV/ms)
dPSP
dSpike
****
*

### Figure S10

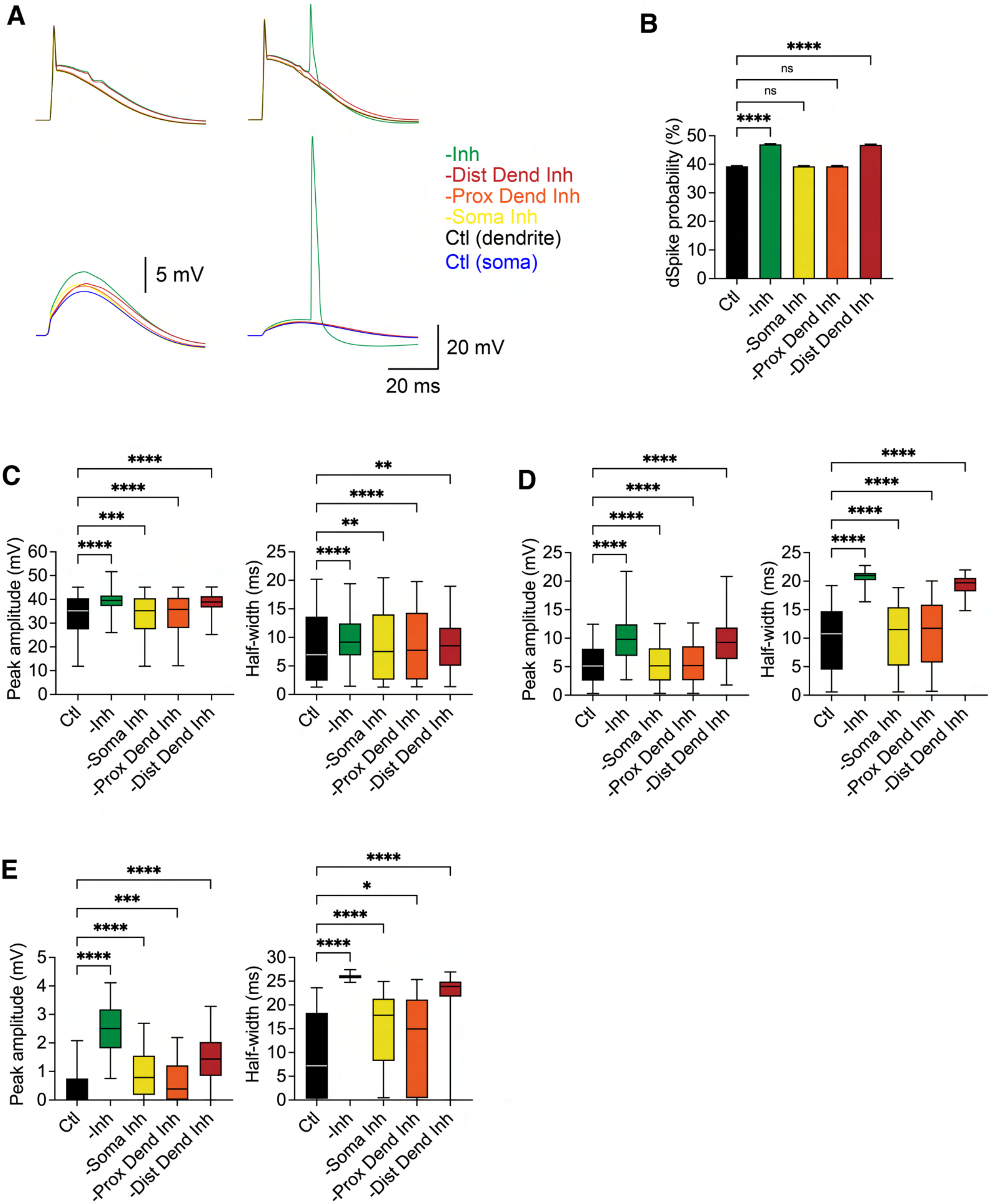
A
B
C
D
E
-Inh
-Dist Dend Inh
-Prox Dend Inh
-Soma Inh
Ctl (dendrite)
Ctl (soma)
5 mV
20 mV
20 ms
dSpike probability (%)
Peak amplitude (mV)
Half-width (ms)
Ctl
-Inh
-Soma Inh
-Prox Dend Inh
-Dist Dend Inh
ns
****
***
**
*

Figure S11

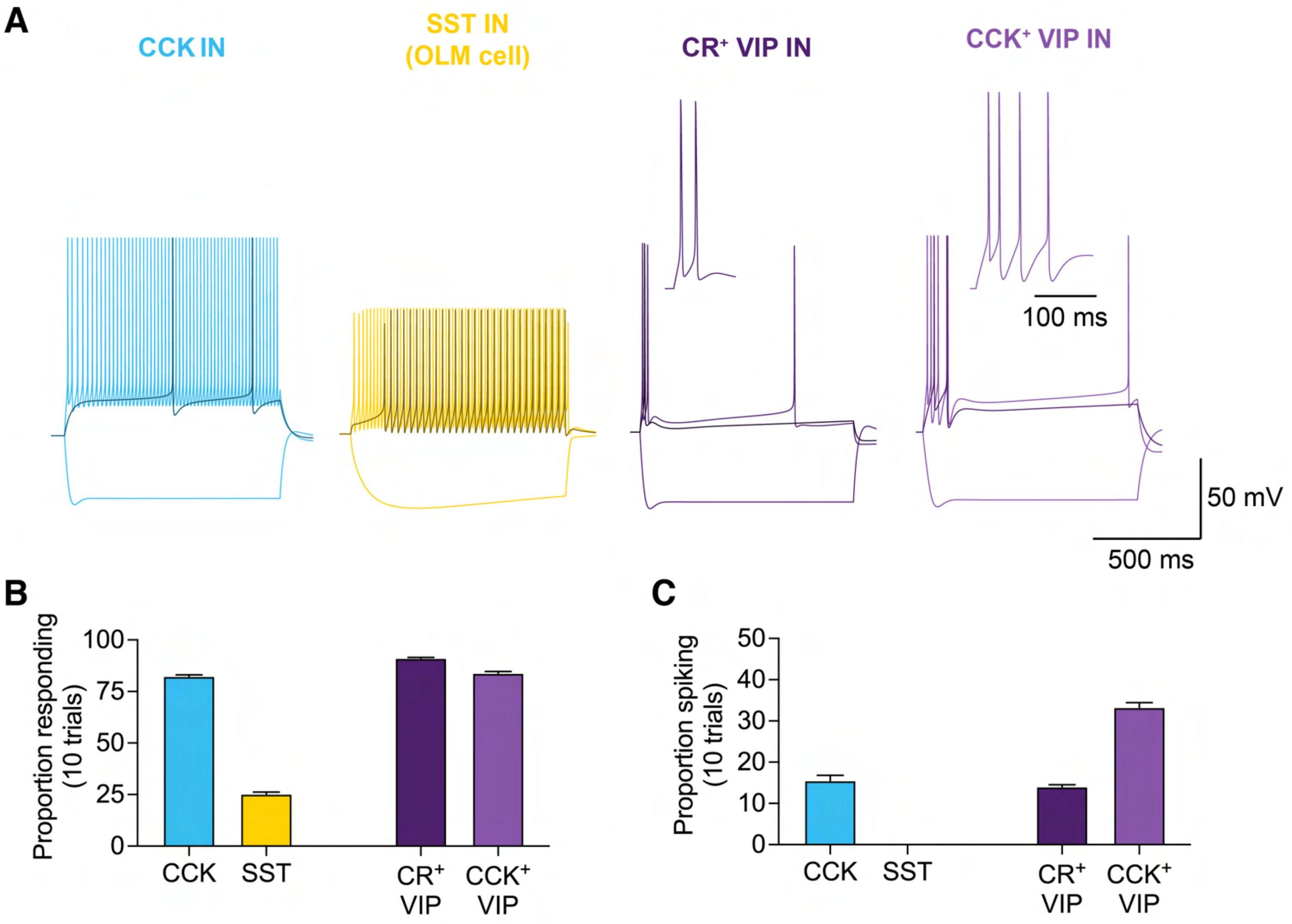

A
CCK IN
SST IN
(OLM cell)
CR+ VIP IN
CCK+ VIP IN
100 ms
50 mV
500 ms
B
Proportion responding
(10 trials)
100
75
50
25
0
CCK
SST
CR+
VIP
CCK+
VIP
C
Proportion spiking
(10 trials)
50
40
30
20
10
0
CCK
SST
CR+
VIP
CCK+
VIP


### Figure S12

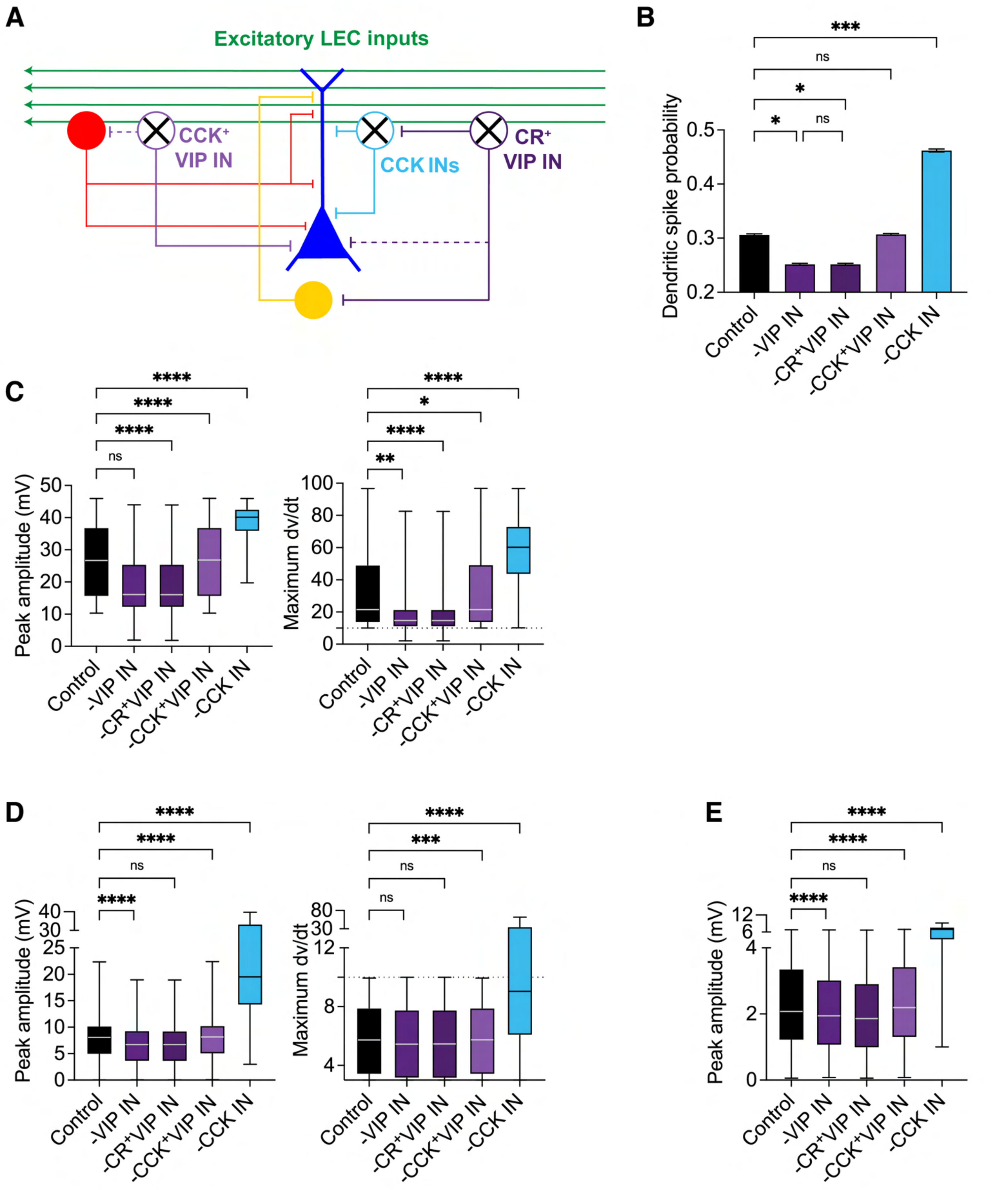
A
Excitatory LEC inputs
CCK+ VIP IN
CCK INs
CR+ VIP IN
B
Dendritic spike probability
Control
-VIP IN
-CR+VIP IN
-CCK+VIP IN
-CCK IN
C
Peak amplitude (mV)
Maximum dv/dt
D
Peak amplitude (mV)
Maximum dv/dt
E
Peak amplitude (mV)

Figure S13

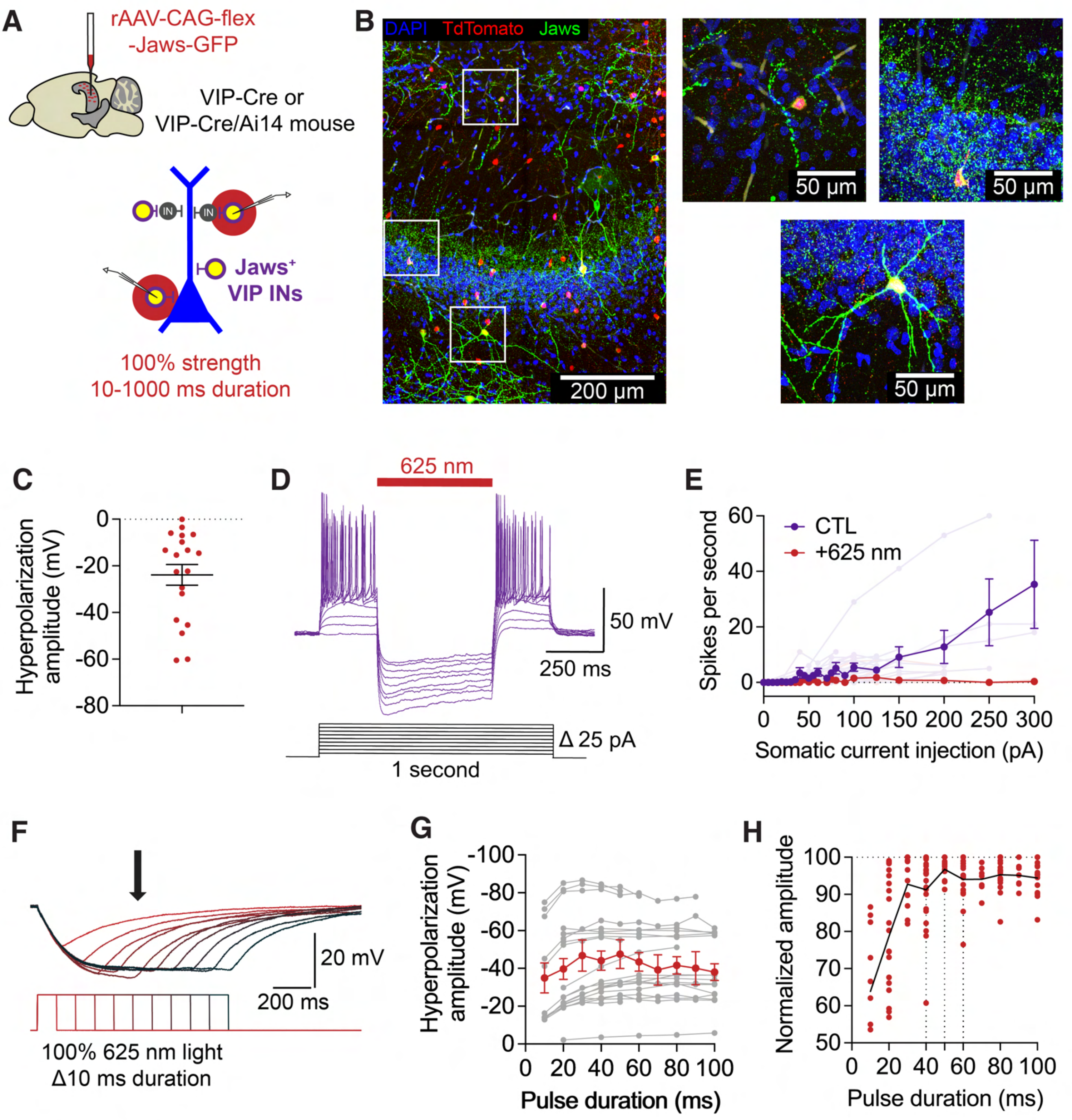
A
rAAV-CAG-flex
-Jaws-GFP
VIP-Cre or
VIP-Cre/Ai14 mouse
IN
IN
Jaws+
VIP INs
100% strength
10-1000 ms duration
B
DAPI TdTomato Jaws
200 µm
50 µm
50 µm
50 µm
C
Hyperpolarization amplitude (mV)
0
-20
-40
-60
-80
D
625 nm
50 mV
250 ms
Δ 25 pA
1 second
E
CTL
+625 nm
Spikes per second
Somatic current injection (pA)
F
20 mV
200 ms
100% 625 nm light
Δ10 ms duration
G
Hyperpolarization amplitude (mV)
Pulse duration (ms)
H
Normalized amplitude
Pulse duration (ms)

### Figure S14

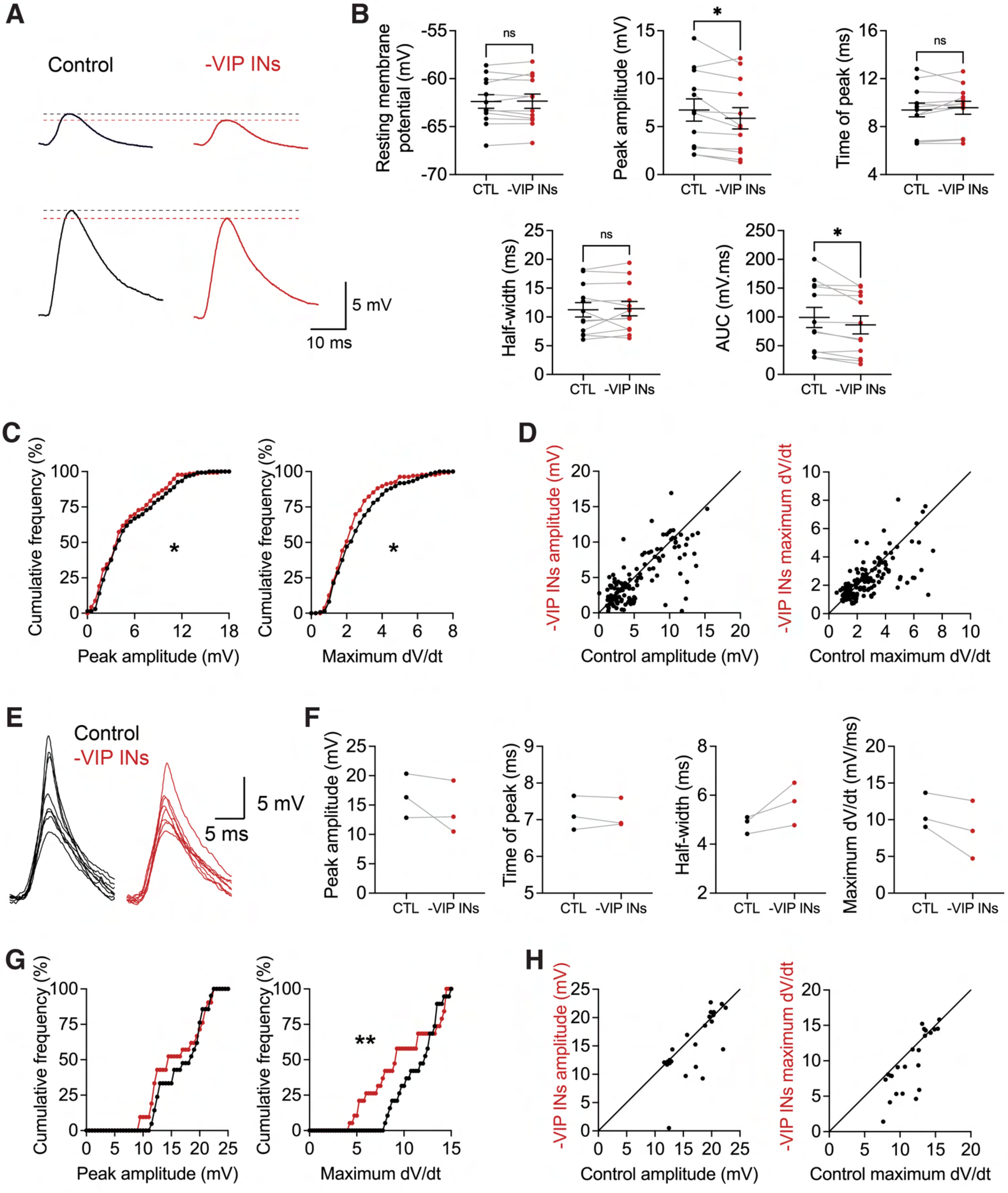
A
Control
-VIP INs
5 mV
10 ms
B
Resting membrane potential (mV)
ns
CTL
-VIP INs
Peak amplitude (mV)
*
Time of peak (ms)
Half-width (ms)
AUC (mV.ms)
C
Cumulative frequency (%)
Peak amplitude (mV)
Maximum dV/dt
D
-VIP INs amplitude (mV)
Control amplitude (mV)
-VIP INs maximum dV/dt
Control maximum dV/dt
E
Control
-VIP INs
5 mV
5 ms
F
Peak amplitude (mV)
Time of peak (ms)
Half-width (ms)
Maximum dV/dt (mV/ms)
G
Cumulative frequency (%)
Peak amplitude (mV)
**
Maximum dV/dt
H
-VIP INs amplitude (mV)
Control amplitude (mV)
-VIP INs maximum dV/dt
Control maximum dV/dt

### Figure S15

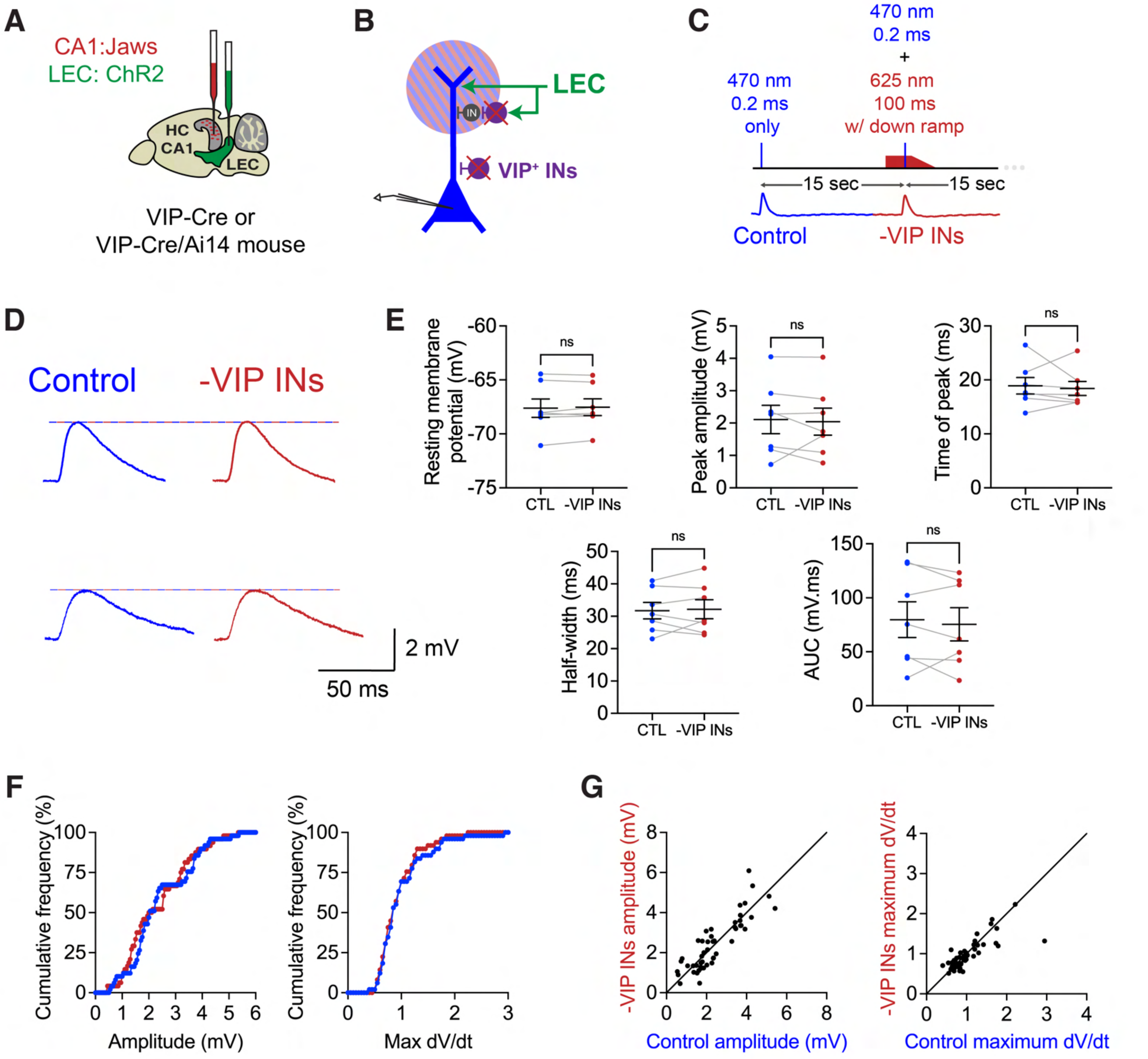
A
CA1:Jaws
LEC: ChR2
HC
CA1
LEC
VIP-Cre or
VIP-Cre/Ai14 mouse
B
LEC
IN
VIP⁺ INs
C
470 nm
0.2 ms
only
470 nm
0.2 ms
+
625 nm
100 ms
w/ down ramp
15 sec
15 sec
Control
-VIP INs
D
Control
-VIP INs
2 mV
50 ms
E
Resting membrane potential (mV)
Peak amplitude (mV)
Time of peak (ms)
Half-width (ms)
AUC (mV.ms)
ns
CTL
-VIP INs
F
Cumulative frequency (%)
Amplitude (mV)
Max dV/dt
G
-VIP INs amplitude (mV)
Control amplitude (mV)
-VIP INs maximum dV/dt
Control maximum dV/dt

### Figure S16

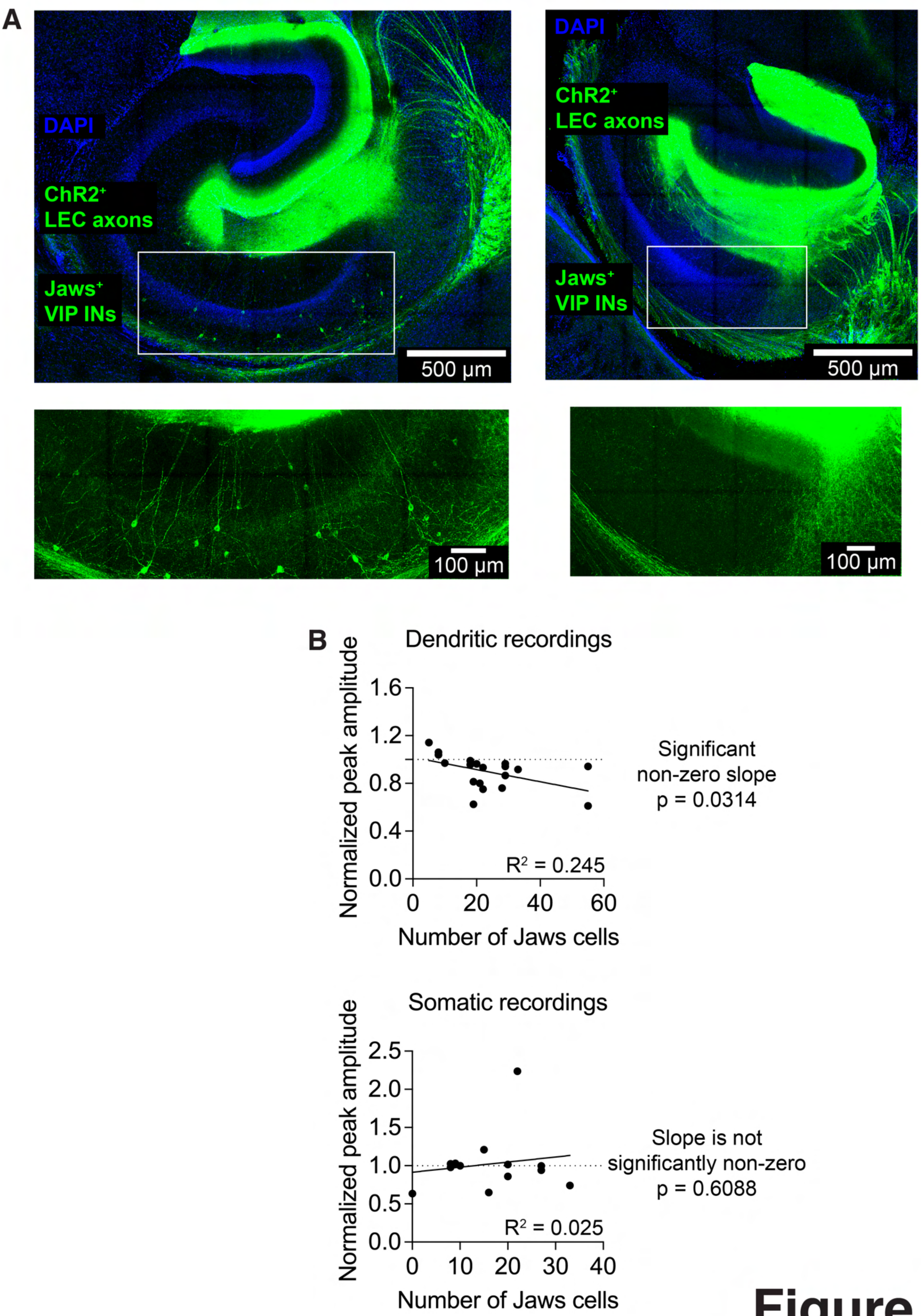
A
DAPI
ChR2+
LEC axons
Jaws+
VIP INs
500 µm
DAPI
ChR2+
LEC axons
Jaws+
VIP INs
500 µm
100 µm
100 µm
B
Dendritic recordings
Normalized peak amplitude
1.6
1.2
0.8
0.4
0.0
0
20
40
60
Number of Jaws cells
R2 = 0.245
Significant
non-zero slope
p = 0.0314
Somatic recordings
Normalized peak amplitude
2.5
2.0
1.5
1.0
0.5
0.0
0
10
20
30
40
Number of Jaws cells
R2 = 0.025
Slope is not
significantly non-zero
p = 0.6088

### Figure S17

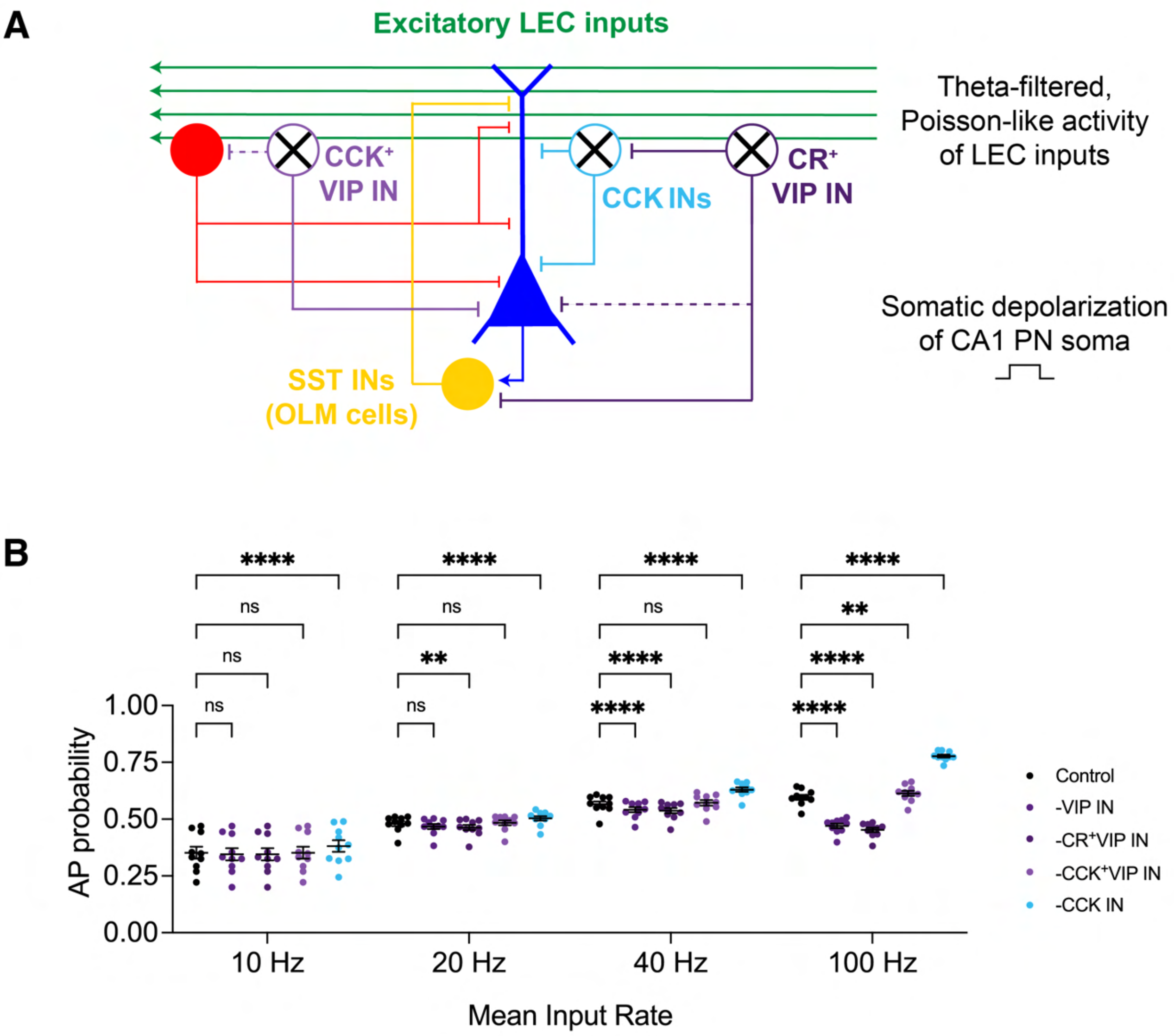

A
Excitatory LEC inputs
Theta-filtered, Poisson-like activity of LEC inputs
CCK+ VIP IN
CCK INs
CR+ VIP IN
Somatic depolarization of CA1 PN soma
SST INs (OLM cells)
B
AP probability
1.00
0.75
0.50
0.25
0.00
10 Hz
20 Hz
40 Hz
100 Hz
Mean Input Rate
ns
**
****
Control
-VIP IN
-CR+VIP IN
-CCK+VIP IN
-CCK IN


### Figure S18
